# MoveTraits – A database for integrating animal behaviour into trait-based ecology

**DOI:** 10.1101/2025.03.15.643440

**Authors:** L.T. Beumer, A.G. Hertel, R. Royauté, M.A. Tucker, J. Albrecht, R.S. Beltran, F. Cagnacci, S.C. Davidson, N. Dejid, R. Kays, A. Kölzsch, A. Lohr, E.L. Neuschulz, K. Safi, A.K. Scharf, M. Schleuning, M. Wikelski, T. Mueller

**Affiliations:** 1Senckenberg Biodiversity and Climate Research Centre (SBiK-F), Frankfurt am Main, Germany; Department of Arctic Biology, The University Centre in Svalbard, Longyearbyen, Svalbard; Behavioural Ecology, Department of Biology, Ludwig-Maximilians University of Munich, Planegg-Martinsried, Germany; Université Paris-Saclay, INRAE, AgroParisTech, UMR EcoSys, Palaiseau, France; Department of Environmental Science, Radboud Institute for Biological and Environmental Sciences, Radboud University, Nijmegen, Netherlands; Department of Ecology and Evolutionary Biology, University of California Santa Cruz, Santa Cruz, USA; Animal Ecology Unit, Research and Innovation Centre, Edmund Mach Foundation, Trento, Italy; Department of Migration, Max Planck Institute of Animal Behavior, Radolfzell, Germany; Department of Biology, University of Konstanz, Constance, Germany; North Carolina Museum of Natural Sciences, Raleigh, USA; Department of Forestry and Environmental Resources, North Carolina State University, Raleigh, USA; Ecology Department, Radboud Institute for Biological and Environmental Sciences, Radboud University, Nijmegen, Netherlands; Department of Biological Sciences, Goethe University, Frankfurt am Main, Germany

## Abstract

Trait-based approaches are key to understanding eco-evolutionary processes but rarely account for animal behaviour despite its central role in ecosystem dynamics. We propose integrating behaviour into trait-based ecology through movement traits - standardised and comparable measures of animal movement derived from biologging data, such as daily displacements or range sizes. Accounting for animal behaviour will advance trait-based research on species interactions, community structure, and ecosystem functioning. Importantly, movement traits allow for quantification of behavioural reaction norms, offering insights into species’ acclimation and adaptive capacity to environmental change. We outline a vision for a ’living’ global movement trait database that enhances trait data curation by (1) continuously growing alongside shared biologging data, (2) calculating traits directly from individual-level data using standardised, consistent methodology, and (3) providing information on multi-level (species, individual, within-individual) trait variation. We present a proof-of-concept ‘MoveTraits’ database with 55 mammal and 108 bird species, demonstrating calculation workflows for 5 traits across multiple time scales. Movement traits have significant potential to improve trait-based global change predictions and contribute to global biodiversity assessments as Essential Biodiversity Variables. By making animal movement data more accessible and interpretable, this database could bridge the gap between movement ecology and biodiversity policy, facilitating evidence-based conservation.

## Box 1

**Glossary**

**Animal biologging**: The use of miniaturized animal-attached tags (‘biologgers’) for recording data about an animal’s movements, behaviour, physiology and/or environment (Rutz & Hays 2009). Traditionally, biologging (tags logging data in memory) is distinguished from biotelemetry (tags transmitting data to a receiver or satellite) (Watanabe & Papastamatiou 2023). However, due to the emergence of hybrid devices and the increasing complementary use of both device types, this distinction is increasingly fading in practice. Here, we collectively refer to both approaches as biologging.

**Animal movement**: In movement ecology, animal movement is typically defined as the change in the spatial location of an individual in time (Nathan *et al*. 2008). It is a fundamental aspect of animal behaviour, operating across multiple scales - from fine-scale local movements to long-distance migrations. With animal biologging, additional dimensions of movement can be quantified, such as wing flaps, head movements, or lying vs. standing. For the purpose of this article, we focus on animal movements as location changes in geographic space.

**Movement metric**: A quantitative measure to describe and analyse the patterns of animal movement in space and time, inferred from biologgers. Typically, metrics either characterise an animal’s path (focussing on the one-dimension aspects of movement trajectories, such as displacement, or speed) or space use (focussing on the two-dimensional patterns, incl. home range size, or utilisation distribution).

**Movement trait**: An agreed-upon movement metric that is calculated in a comparable and standardised way across many taxa and therefore complies with operational trait definitions. We consider movement traits to be a subcategory of behavioural traits (see Fig. 2).

**Trait**: A well-defined, measurable property of organisms, usually measured at the individual level and used comparatively across species (Dawson *et al*. 2021; McGill *et al*. 2006). Common categories are morphological, physiological, phenological, life history, or behavioural traits; see Fig. 2.

**Functional trait**: Traits that influence the organism’s performance under different environmental conditions and/or its effect on ecosystem processes (e.g., seed dispersal, nutrient cycling) (Dawson *et al*. 2021; McGill *et al*. 2006; but see Sobral 2021).

## Introduction

Identifying the mechanisms underpinning organismal interactions is key to move from a simple description of observed patterns to predicting ecological and evolutionary processes (Funk *et al*. 2017). In this context, trait-based approaches have emerged as a promising tool, driving rapid progress over the last two decades (Green *et al*. 2022; McGill *et al*. 2006; Violle *et al*. 2007). Instead of solely focusing on species identity, trait-based ecology posits that an organism’s ecological role and distribution in space and time can be understood through the combination and interaction of a suite of measurable traits (Lavorel & Garnier 2002; McGill *et al*. 2006). Recognising that community and ecosystem functions ultimately arise from the collective interactions of individual organisms, it boosts our ability to identify generalisable principles governing eco-evolutionary processes across scales, for instance by facilitating macro-ecological and macro-evolutionary studies (Funk *et al*. 2017; Harfoot *et al*. 2014; Lavorel & Garnier 2002).

Initially, trait-based concepts, data collections, and insights into community assembly and ecosystem functioning focused almost entirely on plant traits (Kattge *et al*. 2020; Lavorel & Garnier 2002; McGill *et al*. 2006; Suding *et al*. 2008). Given animals’ crucial role in ecosystem functioning, especially in linking trophic levels (Schleuning *et al*. 2023), attention has recently turned to animal functional traits, leading to the curation of global-scale trait databases for various animal taxa, for example PanTHERIA (Jones *et al*. 2009), EltonTraits (Wilman *et al*. 2014), Amniote (Myhrvold *et al*. 2015), the SPI-Birds data hub (Culina *et al*. 2021), AnimalTraits (Herberstein *et al*. 2022), COMBINE (Soria *et al*. 2021), AVONET (Tobias *et al*. 2022), and FuTRES (Balk *et al*. 2022). These databases fill an important data gap to facilitate the integration of animal and plant trait data, critical for a broader understanding of plant-animal interactions as a fundamental pillar of ecological communities. While these global animal-trait databases have been influential (e.g., Newbold *et al*. 2015; Olival *et al*. 2017; Rigal *et al*. 2023; Tucker *et al*. 2023) they remain mostly restricted to morphological traits (e.g., body or brain size) or general ecological traits (e.g., trophic guild, diet), which are relatively stable between and within individuals and are typically sourced from the literature and museum collections (Balk *et al*. 2022; Schleuning *et al*. 2023).

One key aspect that remains underrepresented in trait-based ecology is animal behaviour, despite its recognised importance for many ecological processes ranging from individuals’ fitness to ecosystem-level functions (e.g., Nagelkerken & Munday 2016; Rahman & Candolin 2022; Réale *et al*. 2007; Wilson *et al*. 2020). This is a critical oversight, especially in the context of global change studies (Buchholz *et al*. 2019), where behavioural responses can significantly mediate species’ abilities to cope with changing environments. Evidence of human impacts on animal behaviour (e.g., Suraci *et al*. 2019; Tucker *et al*. 2018; Wong & Candolin 2015), and animals’ behavioural adjustments to environmental change (e.g., Abrahms *et al*. 2018; Johansson *et al*. 2024; Wong & Candolin 2015) is abundant. Such behavioural changes can trigger cascading effects on species interactions, community structure and ecosystem function (Wilson *et al*. 2020). Yet, these insights are often not incorporated into broader trait-based analyses. A primary impediment to integrating behaviour is the limited availability of standardised and comparable behavioural trait data collected from wild animals (Box 2).

A key facet of animal behaviour that can be quantified for wild animals is movement. Animal movements (see Glossary) represent the way animals interact with their biotic and abiotic environment. Their influence on individuals’ survival and reproduction, as well as ecological interactions within and across trophic levels, make them ecologically and evolutionarily relevant (Cagnacci *et al*. 2010; Jeltsch *et al*. 2013). Using bio-logging technologies, movement can be recorded in situ for an ever-growing number of species and individuals (Hussey *et al*. 2015; Kays *et al*. 2015; Nathan *et al*. 2022). However, even though a wealth of animal movement data is available and partly harmonised in shared databases (Davidson *et al*. 2025; Harcourt *et al*. 2019), animal movement is still rarely directly integrated into trait-based approaches. This is due to the fact that the billions of animal location records cannot be readily used as traits in their initial form but need to be processed into informative metrics first. In the past, comparative work mostly relied on sourcing such processed movement metrics from the literature (Tucker *et al*. 2014, but see e.g., Huang *et al*. 2021). A major shortcoming of this approach is that these estimates are derived using varying methodologies and calculated at different levels (e.g., individual versus species). In addition, they are based on data collected with varying sampling intervals, which may introduce error and bias as most methods for deriving metrics from animal movement data are sensitive to sampling rates (Calabrese *et al*. 2016). Therefore, a significant obstacle that remains is the lack of a comprehensive database assembling standardised movement metrics that represent comparable movement traits (see Glossary). This step is essential for animal movement data to be usable for trait ecologists in trait-based approaches.

Given the lack of standardised and comparable movement trait data, the relevance of animal mobility to ecosystem dynamics is often indirectly acknowledged in trait-based studies by incorporating animal movement behaviour via ‘proxies’. These are usually morphological or general ecological traits (hereafter referred to as ‘proxy traits’), available from the recently developed animal-focussed trait databases (Fig. 1). For example, animals’ capacity to disperse seeds is commonly approximated via body size or mass due to a suggested allometric scaling of dispersal movements (Jenkins *et al*. 2007; Sorensen *et al*. 2020; Stevens *et al*. 2014) and daily distance travelled (Carbone *et al*. 2005). While proxy traits may represent simple categorisations of or scale allometrically with species’ average movement behaviour (Sheard *et al*. 2020; Tucker *et al*. 2014), they may overlook critical movement variation by (1) discounting behavioural variation between species with similar proxy traits, (2) not adequately reflecting behavioural variation within species, (3) disregarding spatio-temporal patterns of animal movement and behaviour, and/or (4) not accounting for animals’ behavioural plasticity in responding to fluctuating biotic and abiotic conditions (Fig. 1). Hence, while proxy traits might explain some patterns (e.g. average home range size across mammals with large differences in body size; Tucker *et al*. 2014) and approximate a species’ movement ‘potential’, they may omit key information on the variability and context-dependence of trait effects that are critical to improve our understanding of species interactions, community structure, and ecosystem functioning.

**Fig. 1:**
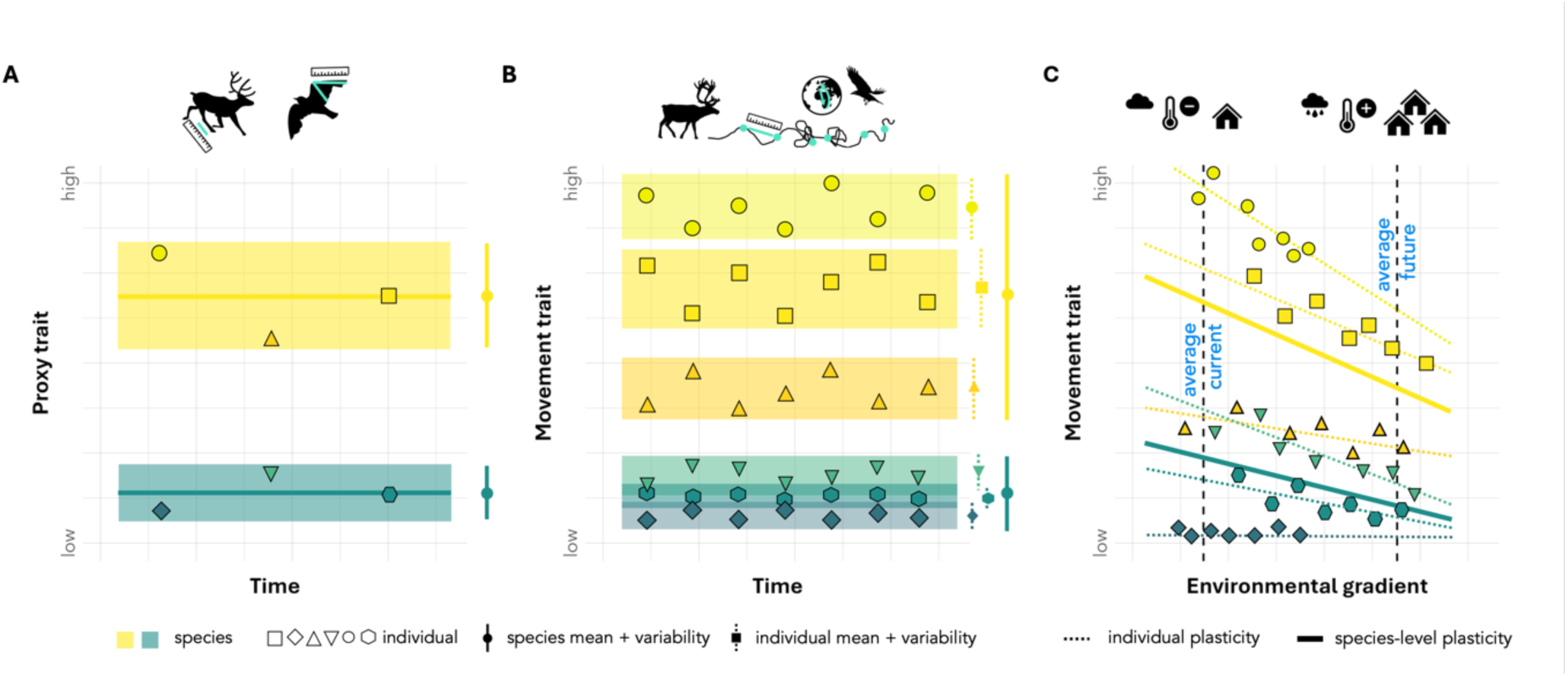
Conceptual advances provided by movement traits. (A) In trait-based ecological studies, animal movement behaviour is often incorporated via ‘proxy traits’ - typically morphological and ecological traits, such as body size/mass, wing morphology, or trophic guild, which are typically derived from museum collections and measured only once per individual. This technically allows for quantification of within-species trait variation, although most trait databases typically only contain trait values averaged at the species level, often without reporting the underlying trait measurements, variance measures, or sample sizes. (B) Biologging devices measure behavioural information repeatedly for the same individual in its natural environment and over ecologically meaningful time periods. Building on these repeated individual-level raw data, movement traits can be calculated in a standardised workflow with consistent methodology, achieving (1) more comparable measures including variance estimates and sample sizes, and (2) quantification of both between-species, between-individual and within-individual trait variation. (C) Finally, animals may display considerable plasticity in behavioural reactions to their encountered abiotic and biotic environment. Individual measures of movement traits along environmental gradients facilitate incorporation of *behavioural reaction norms*, combining consistent between-individual differences in behaviour independent of context (different intercepts) with within-individual behavioural plasticity (individuals adjusting behaviour adaptively to changing conditions; non-zero slopes). Accounting for behavioural reactions norms should enable better predictions of community dynamics and ecosystem processes under global change. Panel C adapted from Hertel *et al*. (2020).

To overcome these limitations, we outline a vision for the development of a movement trait repository as a ‘living’ database, continuously updated and growing alongside biologging databases. We suggest that such a database would provide a new approach for curating trait databases by (1) directly building on individual-level data and (2) facilitating trait calculation in a standardised workflow with consistent methodology. In addition, we propose a way to (3) provide information on multi-level (species, individual, intra-individual) trait variation thus far lacking from existing trait databases. We believe that this approach will achieve more comparable measures and therefore more generality in the inferred ecological insight, which is crucial for effective biodiversity conservation and environmental management. Demonstrating the practical workflow from biologging data to trait calculation, we present a first proof-of-concept animal movement trait dataset based on standardised analysis of biologging data and illustrate how these movement traits can fill critical trait data gaps. We conclude by discussing how such a movement trait database will unlock the potential of biologging data to integrate animal behaviour into trait-based ecology, global change prediction, environmental monitoring, and global biodiversity assessments, with the potential to inform Essential Biodiversity Variables (EBVs) and bridge the gap between movement ecology research and biodiversity policy.

### BOX 2

**Trait-based approaches in behavioural ecology**

Behavioural ecology aims to understand the ecological and evolutionary basis of animal behaviour, and - like trait-based approaches in general - concentrates on the individual as the focal unit of analysis (Carter *et al*. 2013). While in trait-based ecology, traits are typically used to capture differences between species, in behavioural ecology, the use of traits has mainly focussed on behavioural differences between individuals of the same species that are repeatable over time and across situations (i.e., animal temperament or personality, Réale *et al*. 2007). Analogue to the ‘big 5’ temperament traits in human psychological research, initial work focused on quantifying individuals’ aggressiveness, boldness, activity, exploration, and sociability (Réale *et al*. 2007), using experimental tests in the lab or controlled set-ups in the wild (Carter *et al*. 2013). While standardised test set-ups control for environmental confounds, they are impractical to quantify behavioural differences across a wide range of species, causing a gap of comparable behavioural trait studies in the wild. More recently, an alternative approach has emerged to study behavioural traits, purely based on statistical partitioning of behavioural variation into its environmental, among- and within-individual sources (Dall & Griffith 2014; Dingemanse *et al*. 2010; Dochtermann & Dingemanse 2013). This ‘statistical’ approach is not limited to the five major personality traits but is instead founded on the notion that individuals can differ in their average expression of any kind of behaviour (Dall & Griffith 2014; Dingemanse *et al*. 2010). Between-individual behavioural variation is quantified by repeatedly measuring individual behaviour across different biological contexts (Dingemanse *et al*. 2010). Using this approach, intrinsic individual variation (i.e., an individual’s average behaviour) can be discerned from reversible behavioural plasticity. Because behaviour is at least partially heritable (Dochtermann *et al*. 2019), between-individual behavioural variation is the substrate for natural selection and can speed up adaptation to environmental change (Réale *et al*. 2007; Wolf & Weissing 2012).

### Conceptual alignment: Animal movement reflects key behavioural traits

Traits are morphological, physiological, phenological, life history, or behavioural properties of organisms (Fig. 2), measured at the individual level and often averaged per species for comparative cross-species studies (Dawson et al. 2021; McGill et al. 2006; see Glossary). The term ‘functional trait’ refers to traits that influence the organism’s performance under different environmental conditions (Dawson *et al*. 2021; Garnier *et al*. 2015) and/or its effect on ecosystem processes (e.g., seed dispersal, nutrient cycling) (Dawson *et al*. 2021; McGill *et al*. 2006; but see Sobral 2021). Although traits are defined without reference to the environment, measured trait *values* need to be annotated with environmental conditions to facilitate the interpretation of ecological and evolutionary significance (Dawson *et al*. 2021; Garnier *et al*. 2015). Generally, traits, including behavioural traits, are assumed to be heritable to some degree, and thus under natural selection (Dochtermann *et al*. 2019, see Box 2).

**Fig. 2:**
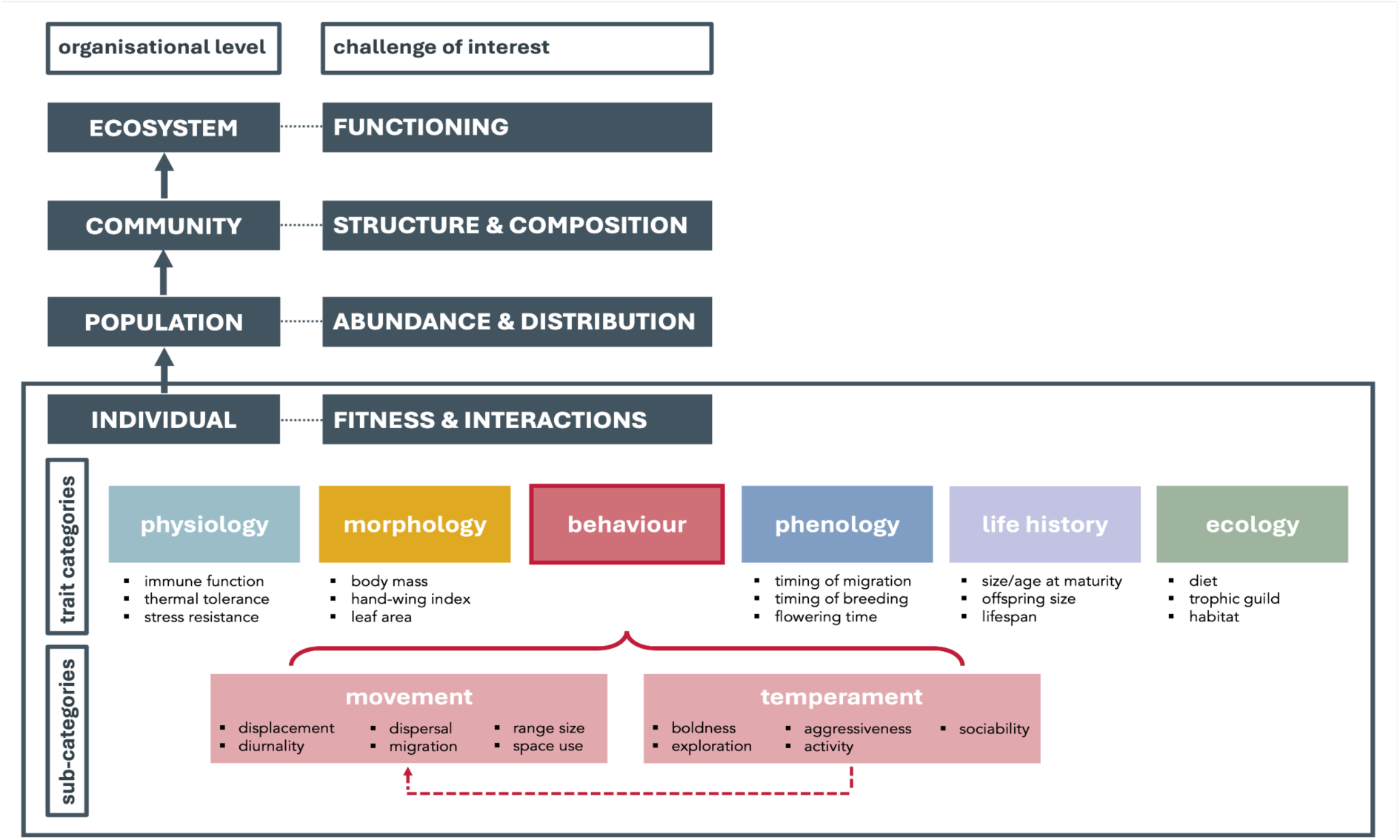
Trait-based approaches have been used in various ecological disciplines, leading to a plethora of trait concepts, definitions, and categories (Violle *et al*. 2007). Here, we suggest a ‘taxonomy of traits’ with movement traits as a subcategory of behavioural traits, as movement always reflects behaviour, but not all behaviour is expressed via movement. However, we note that there can be considerable overlap between trait categories and suggest a pragmatic approach to such classifications.

Metrics inferred from animal movement data can comply with these commonly used operational definitions of traits and have the potential to be useful in comparative approaches (see Glossary): Biologging-derived animal movement data are always collected at the individual level and usually recorded automatically and repeatedly at regular time intervals and without observer bias (Fig. 1). The inferred metrics are, in principle, precisely defined and clearly measurable. However, only those movement metrics that can be calculated in a standardised way and that can be averaged and compared across a number of species (Joly *et al*. 2019; Tucker *et al*. 2018) can serve as useful movement traits. As the recorded locations are spatio-temporally explicit, their annotation with environmental context is feasible. Recent studies demonstrate a genetic basis (i.e., heritability) for movement traits such as migration direction or habitat preferences (Bonar *et al*. 2022; Gervais *et al*. 2020), indicating the potential for evolution and adaptive responses to selective pressures. Given the importance of animal mobility for individual fitness and ecological processes, most movement traits would qualify as functional traits: Animal movement is a critical determinant of individual fitness, for instance by determining success in resource acquisition and risk avoidance (DeMars & Boutin 2018; Gaillard *et al*. 2010; McLoughlin *et al*. 2006; Mitchell *et al*. 2002). Likewise, it underpins key ecological processes of importance for (meta-)community dynamics and ecosystem functioning (Nathan *et al*. 2008; Schlägel *et al*. 2020). These include, for instance, seed dispersal (Graf *et al*. 2024; Kays *et al*. 2011), nutrient transfer (Doughty *et al*. 2016; McLoughlin *et al*. 2016), disease dynamics (Manlove *et al*. 2022; Scherer *et al*. 2020), spread of invasive species (Reynolds *et al*. 2015), and trophic interactions (DeMars & Boutin 2018; Mitchell *et al*. 2002). Consequently, animal movement provides critical ecosystem services and Nature Contributions to People (Kremen *et al*. 2007).

Given this conceptual alignment, we posit that movement data are an under-used source of trait information, i.e., can serve as movement traits to quantify animal behaviour for integration into trait-based ecology. We suggest that such movement traits are a subcategory of behavioural traits (Fig. 2), as movement is always a reflection of behaviour (e.g., foraging, territoriality, exploration) but not all behaviour is necessarily expressed via movements that result in a change of the individuals’ spatial location (e.g., sociability or aggressiveness expressed through facial expressions or other body language). We note, however, that there is considerable overlap between some trait categories. For instance, while migration distance is a movement trait (Fig. 2), the timing of migration is a phenological trait. Similarly, sprint performance or stamina would be a physiological, rather than a movement trait per se. We suggest a pragmatic approach to such trait classifications.

An interesting property of behavioural traits, including movement, is that they are labile traits, meaning that their expression may change from measurement to measurement (Blomberg *et al*. 2003; Scheiner 1993). Morphological traits, like structural size, tarsus or wing length, are usually more static both between and within individuals outside of ontogeny or organism growth periods. This property has several implications for trait measurements and the eco-evolutionary insights they afford: (1) Behavioural traits should always be based on repeated measures to approximate their centrality (e.g., mean or median) and should be accompanied by measures of variability (e.g., variance or standard deviation) (Niemelä & Dingemanse 2017). (2) If repeated individual measures are recorded, they also allow for the partitioning of behavioural variability into between-individual and environmental sources (Dingemanse & Dochtermann 2013). Between-individual variation in movement traits is generally high (Stuber *et al*. 2022) and enables natural selection on existing trait variation, facilitating faster adaptation to environmental change as compared to evolution via genetic mutation (Barrett & Schluter 2008; Wolf & Weissing 2012). Species with greater heritable between-individual variation therefore likely have a greater *adaptive capacity* to fast-paced environmental change. (3) Animals are able to adjust behaviour more readily than morphology to their environmental context, and this trait lability can support acclimation via reversible behavioural plasticity (Beever *et al*. 2017; Dingemanse *et al*. 2010). Individuals can thereby differ in their plasticity to experienced environmental gradients, a scenario under which plasticity can evolve (Nussey *et al*. 2007) – offering a lens into the *acclimation potential* of the species to environmental shifts outside the currently experienced range. Repeated measurements of labile traits like movement therefore allow estimation of the acclimation potential by quantifying changes in behavioural expression along biotic or abiotic environmental gradients, a concept called ‘*behavioural reaction norm*’ (Fig. 1; Dingemanse *et al*. 2010). As biologging devices measure behavioural information repeatedly for the same individual in its natural environment and over ecologically meaningful time periods (Fig. 1; Hertel *et al*. 2020), movement traits thus provide a powerful opportunity for trait-based research to investigate the adaptive capacity and acclimation potential of different species or populations to environmental change via a) the amount of between-individual variability, and b) the degree of between-individual variation in behavioural plasticity to environmental change (Hertel *et al*. 2020, 2021).

### MoveTraits v 0.1.0 - A first proof-of-concept movement trait database

To demonstrate the practical implementation of movement trait calculation, we provide a first proof-of-concept movement trait database (MoveTraits v 0.1.0) based on movement data harmonised and publicly available on Movebank (https://www.movebank.org/; 243 studies, effective March 2025, see Table S3) and open access data published by Tucker *et al*. (2023) (46 studies, Table S3). The dataset includes movement traits from 108 bird and 55 mammal species, represented by 6351 individuals (3660 birds, 2691 mammals).

Animal movement trajectories can be partitioned temporally at scales of steps, daily and seasonal routes, life-history level (e.g. home range), or even lifetime tracks (Kays *et al*. 2023; Nathan *et al*. 2008). To this end, a variety of movement metrics have been developed, facilitating direct comparisons of movements across individuals, populations and species. For example, Abrahms *et al*. (2017) identified several movement metrics useful for classifying broad-scale movement patterns for large marine and terrestrial vertebrate species (e.g. migratory versus range-resident). Similarly, metrics deployed by Tucker *et al*. (2018) proved insightful in determining movement responses to human activity across terrestrial mammal species, and Joly *et al*. (2019) provided simple, comparable, and repeatable metrics for quantifying annual migratory movement distances for terrestrial mammals. For our first proof-of concept version of MoveTraits, we decided on an initial suite of simple movement traits (Table 1) that we consider to (1) provide complementary insights on animals’ movement behaviour, (2) be relevant and comparable across a diverse range of taxa, movement modes, and environmental contexts, and (3) be relatively straightforward to infer from commonly collected animal movement data. This suite is not meant to be a static final set, does not apply to all taxonomic groups, and we hope will be expanded in the future.

**Table 1:** Initial suite of movement trait metrics estimated at the individual level and for different time scales. Metrics were calculated and summarised at the individual level using the entire tracking period of that individual (excluding data gaps) assuming that enough data were recorded to estimate the traits’ centrality and uncertainty (Table S1).

| Movement trait | Time scales | Definition/<br>Calculation | Ecological significance |
| --- | --- | --- | --- |
| Displacement distance [m] | Hourly (1 h),<br>Daily (24 h) | Straight-line Euclidean distance between subsequent locations | Realised movement distances at different time scales |
| Maximum displacement distance [m] | Daily (24 h),<br>Weekly (7 day) | Maximum distance observed across all pairwise distances in a given period | Long-distance movement capacity |
| Range size [km<sup>2</sup>] | Daily,<br>Weekly,<br>Monthly,<br>Annual | 95% Minimum Convex Polygon (MCP) | Overall space requirements |
| Intensity of area use | Daily,<br>Monthly | Ratio between total movement distance (cumulative sum of consecutive distances), divided by the square root of the range size (95% MCP) | Summarising linear and area-based extent of movement; lower values indicate more straight movements (e.g., migrations), higher values more clustered movements |
| Diurnality index [-1 to 1] | Entire tracking period | The relative activity during the day conditional on day length | Indicates whether movement occurs primarily during the day (1) or night (-1) |

First, we resampled the GPS data to regular time intervals of 1 hour (‘hourly’), 24 hours (‘daily’), and 7 days (‘weekly’). Then, we used the resampled data to derive movement metrics at different timescales: We calculated hourly and daily step length, i.e., the straight-line distance between locations. We also quantified daily, weekly, and annual maximum displacement distances as the maximum of all pairwise (i.e., not consecutive) distances within the sampling period. To summarise individuals’ space requirements, we delineated range size from the 95% Minimum Convex Polygon of all GPS locations at a daily, weekly, monthly, and annual timescale. In addition, we also calculated Intensity of Use, a metric that summarises the spread of movement from more linear to more clustered movement patterns (Almeida *et al*. 2010). Finally, we calculated a diurnality index, indicating whether movement (based on hourly steps) occurs mostly during the local night or day (Hoogenboom *et al*. 1984). For more details on data requirements and trait calculations, see Table S1 and Supplementary Methods S1.

MoveTraits v 0.1.0 provides the movement trait data at three hierarchical levels (Fig. 1, 4; Box 3): (1) summarised at the *species level*, facilitating interoperability with other species-level trait databases, (2) summarised at the *individual level* for studies on between-individual variation, and (3) the underlying repeated movement trait estimates for each individual over time to allow for research questions at the *intra-individual level*. We summarised the underlying repeated movement trait estimates at the individual level as mean, median, coefficient of variation, and 5^th^ as well as 95^th^ percentile and provided species means of centrality and variance (e.g., the mean species 95^th^ percentile summarized from individual 95^th^ percentiles for a given trait), facilitating different ecological inquiries. For example, the mean daily displacement distance may signify an individual’s daily routine movements, while the 95^th^ percentile represents an individual’s occasional daily long-distance movements. Each metric was only estimated for individuals with sufficient repeated measures to allow for meaningful measures of centrality and variance (Table S1). For instance, we only summarised daily displacement distance (Table 1) for individuals with at least 30 days of data. Therefore, not all traits are calculated for every individual. We annotated each individual’s trait summary with mean coordinates (to allow spatial annotation with environmental data), tracking period (length, start and end date; for temporal annotation with environmental data), species name and common name, taxonomic class (mammal, bird), movement mode (walk, fly, swim), individual information such as sex, age, and body mass (if available), and details of the data owner (see also Table S1).

This multi-level approach is novel: most trait databases only contain trait values averaged at the species level (but see Balk *et al*. 2022), reported without the underlying individual-level trait measurements, variance measures, or sample sizes (Balk *et al*. 2022; Beltran *et al*. 2025; Moran *et al*. 2016). Therefore, information critical for interpreting species-level trait values are usually missing, complicating error tracing, preventing discoverability, replicability, and interoperability of trait data, and limiting ecological insight (Balk *et al*. 2022; Bartomeus *et al*. 2016; Moran *et al*. 2016). Our multi-scale approach explicitly avoids such a reduction to species-mean values, while still facilitating easy interoperability with existing species-mean trait databases. The individual-level trait data and metadata ensure user flexibility, enabling aggregation to any desired organisational level (e.g., population).

#### BOX 3

**Examples of mammal movement traits at the species, individual, and within-individual level**

To showcase the utility of movement traits summarised at the three hierarchical levels, we present three examples.

*Example 1 (species level):* Morphological traits are commonly used as proxies to account for species movement capacity in trait-based studies. Animal movements, such as home range size (Tucker *et al*. 2014) or maximum dispersal distances (Stevens *et al*. 2014), have been shown to allometrically scale with body mass in terrestrial animals. For example, body size alone has been reported to explain 52% of the variance in home ranges size of terrestrial mammals (Tucker *et al*. 2014). Similarly, the hand-wing index, a metric of bird wing morphology linked to the wing aspect ratio, has been shown to correlate with flight efficiency, dispersal distance, and long-distance movements in birds (Lockwood *et al*. 1998; Sheard *et al*. 2020). To test how closely empirically derived movement traits relate to proxy traits, we linked the species-mean range size trait to the species-mean body mass trait from the PanTHERIA mammal trait database, for all terrestrial mammal species with at least one individual with 14 days of daily location data (n = 42; Table S1). While the general positive relationship between body mass and range size holds, it explains limited variance (R^2^ = 0.22) and substantial uncertainty remains around the trend line (Fig. 3A, see also Fig. S1 for a correlation between hand-wing index and flight distance). Accounting for movement capacity directly, when possible, instead of via proxy traits will thus allow refinement of trait-based studies.

*Example 2 (individual level):* Trait-based studies traditionally rely on species-averaged traits. However, between-individual variation can be substantial and is typically not accounted for. Importantly, between-individual variation may not be the same across species and is generally expected to increase with the trait mean. Using daily displacement distances summarised at the individual level from species with at least 10 unique individuals (n = 27; Table S1), we find great variation among species in their between-individual variance (Fig. 3B). These findings imply that species averages may reflect the movement capacity of some species (e.g., moose, Fig. 3B), while for others, these averages do not capture the breadth of behavioural diversity (e.g., blue wildebeest or reindeer, Fig. 3B). Disregarding between-individual variance may therefore impede correct ecological inference.

*Example 3 (within-individual level):* As movement data are collected at the individual level and repeatedly through time, they facilitate the observation of individual phenotypic expression of movement traits in response to the environment, i.e., individual phenotypic plasticity. The reaction norm framework enables the quantification of between-individual variation in phenotypic plasticity via random slopes. We here highlight the movement responses of two non-migratory mule deer along a gradient of human disturbance in the western United States (Utah and Wyoming), measured as Human Footprint Index (Fig. 3C). The mean daily location information that is provided at the within-individual level for repeated measurements of the trait ‘daily displacement distance’ allows for the annotation of movement traits with environmental covariates. In our example, the two individuals vary in their baseline daily displacement distance in undisturbed landscapes (420 m versus 1600 m at HFI 6) as well as in their responsiveness to human disturbance, with one individual decreasing daily displacements while the other one does not adjust displacements. While reductions in mammalian movements to human disturbance have been shown globally (Tucker *et al*. 2018), our example demonstrates the importance and potential of studying individual responses via repeated trait measures.

See Fig. S1 for corresponding relationships for bird species.

**Fig. 3:**
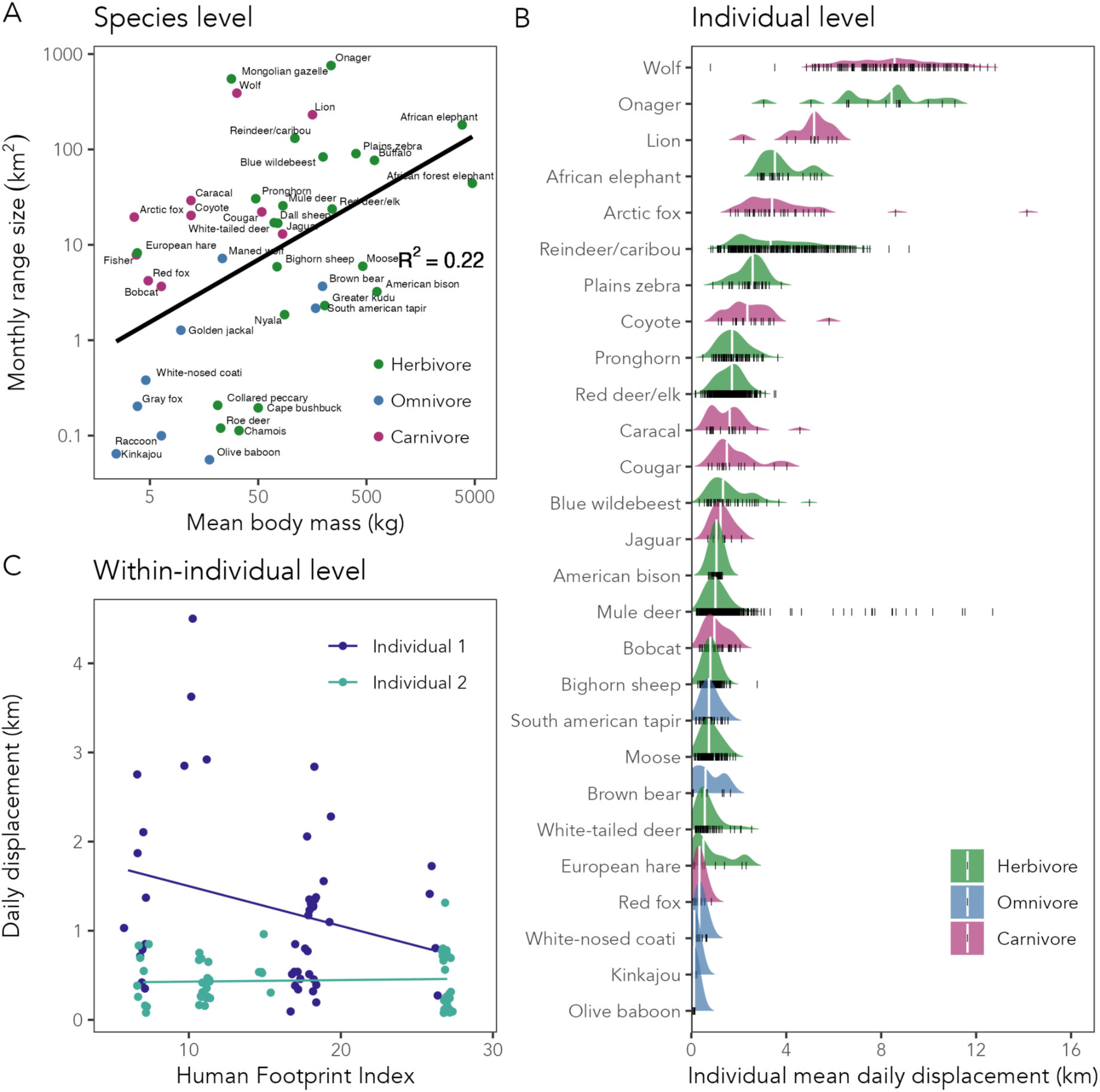
Examples showcasing the utility of movement traits at the species, individual, and within-individual level, extracted from our first proof-of-concept MoveTraits database. (A) Log-transformed species mean body mass against log-transformed monthly range size for 42 terrestrial mammal species for which monthly range size could be calculated. (B) Between-individual variation in mean daily displacement distance for 27 mammal species for which daily displacements were available. Individual means are indicated with black lines and their overall distribution at the species level with density ridges, with species means indicated by white vertical lines. (C) Illustrative example: Daily displacement distances of two non-migratory mule deer (*Odocoileus hemionus*) measured along a gradient of human disturbance in the central United States (Utah & Wyoming), quantified by the Human Footprint Index. A significant interaction between individuals and HFI (F(1, 2229679) = 4.863, p = 0.029) suggests that the two individuals adjust movement differently to HFI. Data were collected between March - May 2019 by Matthew Kauffman and Julie Young and were sourced from the Tucker *et al*. (2023) open-access dataset.

### Towards a global ‘living’ MoveTraits database

#### Envisioned Implementation

Ultimately, we envision an open, ‘living’ movement trait database established as an extension to one or several of the existing biologging databases. Many biologging databases exist (Davidson *et al*. 2025; Harcourt *et al*. 2019) with different regional, taxonomic (e.g., Urbano & Cagnacci 2021), or data/device type foci (e.g., Iverson et al. 2019). Combined, they store billions of animal locations and serve as ‘digital collections’ of animal movement behaviour (Kays *et al*. 2022; Wikelski *et al*. 2024). Automating the calculation of movement traits from these digital collections in a standardised workflow (Fig. 4) is the logical next step. Over time, the movement trait database could thus grow alongside these databases in terms of regions, species, number of individuals covered, and trait metrics extracted, advancing trait databases from relying on published reports to updating dynamically with incoming data. It could also extend trait calculations to additional dimensions of behaviour that can be captured by biologging, for example through measurements of acceleration, body temperature, and heart rate (Hussey *et al*. 2015; Kays *et al*. 2015). This automated trait curation process can be supported by parallel developments of data and metadata standards, and ‘stewardship’ for biologging data management (Davidson *et al*. 2025). Leveraging existing infrastructure and data to create and share movement traits will accelerate research in fields ranging from conservation biology to global change ecology, connecting researchers across fields.

**Fig. 4:**
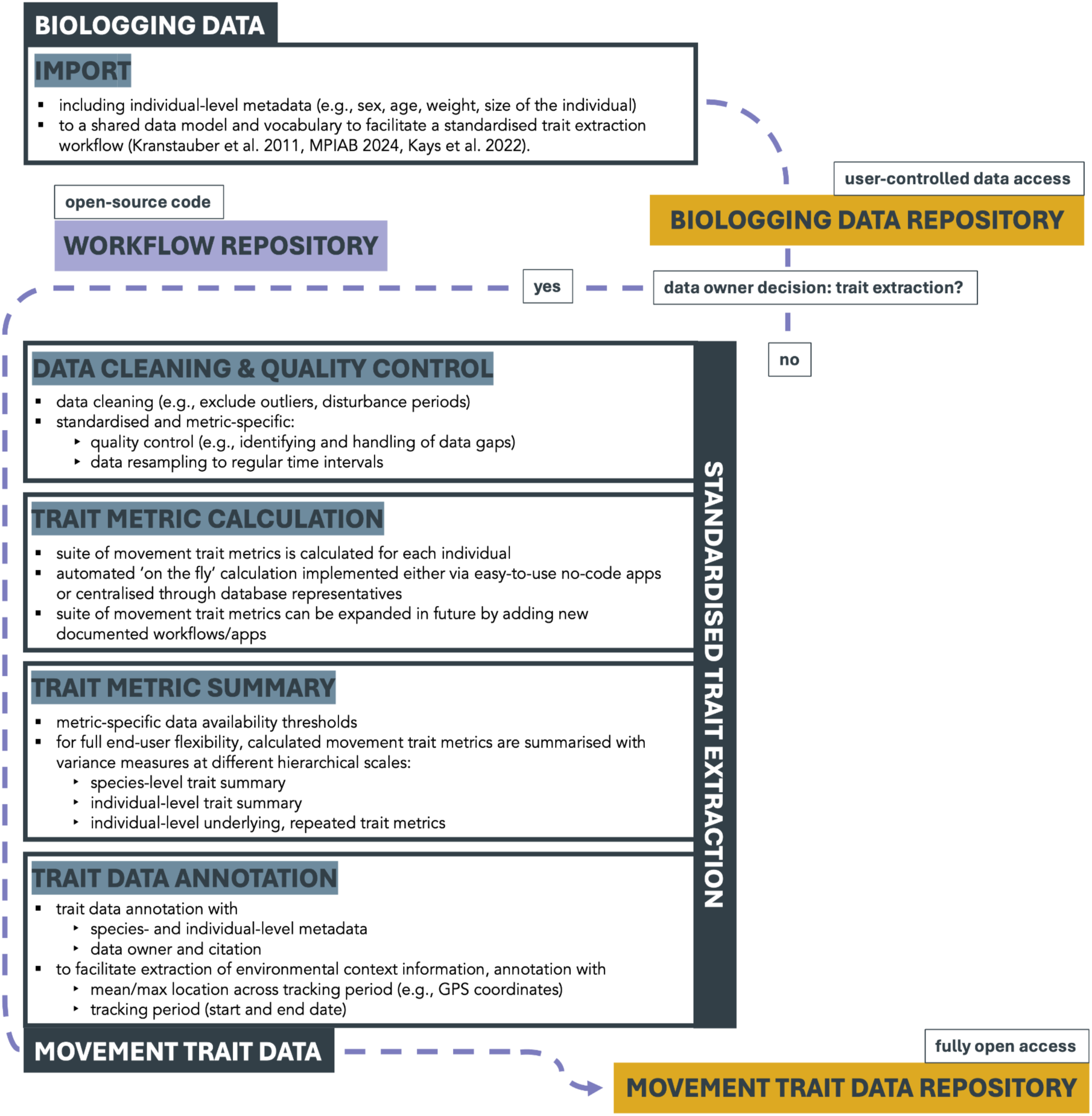
Envisioned standardised workflow from (1) animal biologging data import and organization in a biologging data repository, to (2) standardised and metric-specific data cleaning and quality control, trait metric calculation, summary, and annotation, to (3) trait data archiving in a connected movement trait data repository (‘MoveTraits’ database). All workflow steps are automated and stored in a workflow repository as open-source code. While data access in the biologging data repository is user-controlled (with embargo and access restriction options to protect research agendas and ensure compliance with conservation requirements), all movement trait data is published fully open access, following the FAIR principles (Wilkinson *et al*. 2016) to improve the findability, accessibility, interoperability, and reuse of data.

Currently, the lack of publicly available movement data is a key obstacle to achieve the promising integration of movement and trait data: While trait databases include data across almost 20,000 species, only 1.8% of these species also have publicly available movement data in Movebank (Beltran *et al*. 2025). In Movebank and many other biologging databases, access to a large majority of data is restricted, for example to support storage of real-time and sensitive species locations (Davidson *et al*. 2025). However, databases support restricted sharing, thus facilitating participation by owners of restricted-access data in biologging databases, who could choose to allow trait calculation from their tracked individuals without making their underlying biologging data public. For species- and individual-level trait data calculated from closed-access biologging data, contextualisation with spatio-temporally explicit environmental conditions would be enabled through the reporting of the study period, mean location and location extent. To facilitate environmental annotation at the within-individual level, we envision trait annotation with coordinates on a standardised spatial grid, e.g., at a 30 km resolution. This way, individuals’ exact locations are protected, while critical information about the general spatial context of the individuals’ movements can nonetheless be retrieved. Facilitating environmental data annotation is critical for movement traits, in particular because animals adjust movement first and foremost to the environment (Nathan *et al*. 2008).

Movebank is currently the largest animal movement database and provides a suite of tools for data cleaning, annotation, visualisation, and automated analysis workflows (Kays *et al*. 2022; Kölzsch *et al*. 2022). Its MoveApps platform supports no-code analysis using open-source modules, creating a possibility for MoveTraits to be based on repeatable trait calculation procedures that are open for contribution by other researchers. We therefore suggest Movebank as the basis for MoveTraits v 0.1.0 (Fig. 4) and propose integrating the calculation and storage of movement trait data as an extension to the Movebank ‘ecosystem’. However, if interoperability between biologging databases grows (Davidson *et al*. 2025; Sequeira *et al*. 2021), MoveTraits could be expanded to extract traits across databases and be hosted independently.

#### Implementation Challenges

The integration of movement traits into trait-based ecology faces several challenges. Foremost among these are size-based constraints of biologging technology, which have led to a significant data bias towards larger species (Beltran *et al*. 2025; Kays *et al*. 2015). This gap currently hampers cross-taxa generalisations but is expected to narrow as the miniaturisation of tags and the development of novel non-attached remote sensing devices (e.g., videography of flying insects, see Vo-Doan *et al*. 2024) expand the range of trackable species.

Another challenge is the scale-sensitivity of movement metrics. Most methods for deriving metrics from animal movement data are sensitive to sampling rate and sampling duration (Calabrese *et al*. 2016). This scale-sensitivity stems from interacting biases related to sampling frequency (De Solla *et al*. 1999), movement path tortuosity (Rowcliffe *et al*. 2012), tracking device measurement error (Ranacher *et al*. 2016), and multi-scale autocorrelation inherent in movement data (Fleming *et al*. 2014). Consequently, current discrete-time approaches risk reflecting sampling schedules rather than underlying movement processes. Here, novel continuous-time approaches offer promising solutions for scale-insensitive estimations (Calabrese *et al*. 2016), which should be adopted in future implementations.

Finally, biologging databases have been designed primarily to manage data for individual projects. Steps are needed to prepare these data for contribution to aggregate products like MoveTraits (Fig. 4; Davidson *et al*. 2025; Sequeira *et al*. 2021). For example, data owners may need incentives to make their data fit for use by providing sufficient metadata, applying study-specific outlier detection, and uniquely identifying individuals across projects and databases (Wikelski *et al*. 2024). In addition, MoveTraits will grow through approaches to incorporate the expertise of contributors and credit them in ways that are recognized by citation trackers, employers and funding agencies.

#### Novel Research Avenues

A comprehensive database with movement traits provided at distinct hierarchical levels, interoperable with other trait databases, opens up new opportunities to address key eco-evolutionary questions (Beltran *et al*. 2025; Box 4) and to improve global change predictions. First, this database will enable comparative studies to understand the *causes of variation in movement traits*, i.e., whether/to which degree there are generalisable patterns between movement and other traits. Secondly, the database will improve our ability to study the *ecological consequences of movement trait variation* across scales. In aiding the deciphering of ecological interaction networks in space and time, this will afford novel insights into crucial processes such as seed dispersal, nutrient transfer, or disease transmission across landscapes, linking individual-level behaviours to ecosystem-level processes (e.g., Graf *et al*. 2024) and animal species to ecosystem services (e.g., Savoca *et al*. 2021). Trait data at the within-individual level should promote better accounting for spatio-temporal patterns of animal movement which are typically masked in species-level trait data but important for accurate estimations of ecological interactions (e.g., Kays *et al*. 2011). Similarly, it will improve our capability to assess the *evolutionary consequences of movement trait variation.* Species that display greater between-individual variance, on which natural selection can act, are more likely to adapt to rapid environmental changes than species with little between-individual variance (Barrett & Schluter 2008; Wolf & Weissing 2012). Estimating between-individual variance is therefore essential to predict the evolutionary potential of movement traits and animal behaviour. Finally, movement traits will facilitate the *integration of animal behaviour into global change prediction and environmental management*. While changes in species abundance, distribution, physiology, morphology, and phenology in response to global change have been widely documented, the role of animal behaviour as a mechanism for acclimation and adaptation remains underexplored (Beever *et al*. 2017; Buchholz *et al*. 2019). This oversight is problematic as it neglects a key pathway through which animals can react to rapidly changing conditions. Therefore, incorporating animal behaviour into predictive models is essential for anticipating individual and population responses under future environmental change scenarios. Especially the ability to account for individual behavioural reaction norms will boost a more mechanistic understanding - critical to robust predictions of altered dynamics under novel conditions (Moran *et al*. 2016).

#### Relevance for Global Biodiversity Assessments

Movement traits derived from biologging data have significant potential to contribute to global biodiversity assessments. Movement traits such as migration distances, home range sizes, or daily movement rates could form the basis for new Essential Biodiversity Variables (EBVs) or indicators of biodiversity health, making movement data more policy-relevant (Jetz *et al*. 2019; Kissling *et al*. 2018; Pereira *et al*. 2013; Russo *et al*. 2025). Movement is indeed already listed as an EBV under ‘species traits’ (https://geobon.org/ebvs/what-are-ebvs/), but without clear definition and ignoring within-species variation. Our MoveTraits database could quantitatively characterise movement as an EBV at multiple scales. Since all traits at the individual- and within-individual level are time-annotated, monitoring these variables could provide crucial information about species’ responses to environmental changes, habitat fragmentation, or human disturbances. For instance, changes in migration patterns or home range sizes across populations or species could serve as early warning indicators of ecosystem stress or biodiversity loss. Daily movement rates might indicate habitat quality or resource availability. Such movement-based EBVs could complement existing biodiversity indicators, providing a more comprehensive picture of ecosystem health and functioning.

Moreover, standardised movement traits could enhance the assessment of ecosystem services and Nature Contributions to People that depend on animal movement (Kremen *et al*. 2007). This information could be invaluable for conservation planning and sustainable resource management. By making movement data more accessible, comparable, and interpretable in the context of biodiversity assessments, MoveTraits can help bridge the gap between movement ecology research and biodiversity policy (Jeltsch *et al*. 2013). It could provide policymakers with quantifiable, comparable metrics of animal behaviour and ecosystem functioning, facilitating evidence-based decision-making in conservation and environmental management.

##### BOX 4

**Key eco-evolutionary questions to be addressed with the MoveTraits database**

We foresee exciting research questions along four main themes that could be addressed through our MoveTraits database. We highlight a few examples below:

1. *Understanding causes of variation in animal movement*

a. Are some species more consistent in their movement trait expression while others show greater variability, akin to movement specialists and generalists?
b. Is the degree of species movement specialisation shaped by other species traits, such as longevity, social behaviour, trophic level, or breeding strategy?
2. *Ecological consequences of movement trait variation*

a. How does movement trait expression affect different types of species interactions, such as trophic, competitive, or mutualistic interactions?
b. Can movement traits capture species’ role in community structure and their contributions to ecosystem functions?
3. *Evolutionary consequences of movement trait variation*

a. Are species that display greater within- or between-individual behavioural variance more likely to acclimate/adapt to rapid environmental changes than species with less variance which selection can act upon?
b. What are the fitness consequences of divergent behavioural strategies and how stable are such strategies through time or across distinct populations?
4. *Integrating animal behaviour into global change predictions*

a. Can incorporating animal movement behaviour into predictive models help to better understand individual, population, community, and ecosystem responses to environmental change?
b. Which environmental stressors have the most profound impacts on movement traits?

## Conclusion

The creation of a movement trait database provides a unique opportunity to unlock a valuable and rapidly growing pool of standardised behavioural information for trait-based ecology. Our proposed database outlines a novel pathway to trait data curation that allows for semi-automated, continuous growth instead of one-off static summaries, enables greater data sharing through open-access trait data even if access to the underlying biologging data is restricted, and fosters an open-source collaborative development of reproducible and standardised trait calculation workflows. The integration of movement behaviour into trait-based ecology represents a significant step towards a more comprehensive understanding of community and ecosystem dynamics. By providing standardised, comparable measures of animal movement behaviour, movement traits will enrich eco-evolutionary research and enhance our ability to predict ecosystem responses to global change. Especially the new possibility to integrate individual plastic responses via behavioural reaction norms into trait-based global change ecology constitutes a major conceptual advance and an effective link between the fields of trait-based and behavioural ecology. Furthermore, the relevance of these movement traits extends beyond academic research into global biodiversity assessments and policy. As potential EBVs and indicators of ecosystem health, MoveTraits will make biologging data more accessible and meaningful to policymakers and conservation practitioners.

In an era of rapid environmental change, the ability to quantify and predict animal responses across scales – from individual behaviour to ecosystem functioning – is crucial. Our proposed movement trait database promises to be a valuable tool in this endeavour, representing a significant step towards a more mechanistic understanding of ecological systems, crucial for effective biodiversity conservation and environmental management.

## Acknowledgements

AGH has received funding from the German Science Foundation (HE 8857/1-1). TM and ND received funding from the German Federal Ministry of Education and Research (BMBF, 01LC2320A) through the project MoreStep. ELN has received funding from the German Science Foundation (NE 1863/2-2, NE 1863/3-2, NE 1863/4-1). SCD has received funding from NASA’s Ecological Forecasting Program (80NSSC21K1182).

We want to thank all data owners who allowed us to include their valuable data in the MoveTraits v 0.1.0 database. We emphasize that without these open-access datasets, we would not have been able to derive this first proof-of-concept database. In particular, we thank Matthew Kauffman and Julie Young for providing the data of two mule deer used in the illustrative example in Fig. 3C (data sourced from the open access dataset published in Tucker *et al*. 2023) and Wolfgang Fiedler and Daniel Schmidt-Rothmund for allowing us to use their Eurasian griffon vulture data for the illustrative example in Fig. S1C (Movebank study ‘Raptors NABU Moessingen public’, Study ID 186178781).

