## Supplementary Materials S1 for "MoveTraits – A database for integrating animal behaviour into trait-based ecology"

### APPENDIX S1

#### MoveTraits - Integrating animal behaviour into trait-based ecology

Beumer, L.T.[*,1,2,†], Hertel, A.G.[*,3], Royauté, R.[4], Tucker, M.A.[5], Albrecht, J.[1], Beltran, R.S.[6], Cagnacci, F.[7], Davidson, S.C.[8,9], Dejid, N.[1], Kays, R.[10,11], Kölzsch, A.[8,12], Lohr, A.[10], Neuschulz, E.L.[1], Safi, K.[8], Scharf, A.K.[8], Schleuning, M.[1], Wikelski, M.[8,9] and Mueller, T.[1,13]

*Shared first authorship

[1] Senckenberg Biodiversity and Climate Research Centre (SBiK-F), Frankfurt am Main, Germany

[2] Department of Arctic Biology, The University Centre in Svalbard, Longyearbyen, Svalbard

[3] Behavioural Ecology, Department of Biology, Ludwig-Maximilians University of Munich, Planegg-Martinsried, Germany

[4] Université Paris-Saclay, INRAE, AgroParisTech, UMR EcoSys, Palaiseau, France

[5] Department of Environmental Science, Radboud Institute for Biological and Environmental Sciences, Radboud University, Nijmegen, Netherlands

[6] Department of Ecology and Evolutionary Biology, University of California Santa Cruz, Santa Cruz, USA

[7] Animal Ecology Unit, Research and Innovation Centre, Edmund Mach Foundation, Trento, Italy

[8] Department of Migration, Max Planck Institute of Animal Behavior, Radolfzell, Germany

[9] Department of Biology, University of Konstanz, Constance, Germany

[10] North Carolina Museum of Natural Sciences, Raleigh, USA

[11] Department of Forestry and Environmental Resources, North Carolina State University, Raleigh, USA

[12] Ecology Department, Radboud Institute for Biological and Environmental Sciences, Radboud University, Nijmegen, Netherlands

[13] Department of Biological Sciences, Goethe University, Frankfurt am Main, Germany

*Table S1: Initial suite of movement trait metrics calculated at the individual level*

| **Trait metric** | **Definition/ Calculation** | **Fix rate threshold** | **Data availability threshold** | **Values reported in MoveTraits database** |
|---|---|---|---|---|
| Displacement distances | | | | |
| Hourly [m] | Straight-line displacement distance between subsequent 1h locations | hourly | 167 locations (7 days) | Mean, median, CV, 5th and 95th percentile |
| Daily [m] | Straight-line displacement distance between subsequent 24 h relocations. | daily | 30 locations (30 days) | Mean, median, CV, 5th and 95th percentile |
| Daily - maximum [m] | Observed maximum distance of all pairwise hourly distances in a 24 h period | hourly | Days with at least 12 locations; minimum 7 days | Mean, median, CV, 5th and 95th percentile<br><br>Summarized across all days within criteria |
| Weekly - maximum [m] | Observed maximum distance of all pairwise daily distances in a 1-week period | daily | Weeks with at least 5 relocations; minimum 14 weeks | Mean, median, CV, 5th and 95th percentile<br><br>Summarized across all weeks within criteria |
| Annual - maximum [m] | Observed maximum distance of all pairwise weekly distances in a 1-year period | weekly | Minimum 36 weeks, i.e. 9 months of data | Mean, median, CV, 5th and 95th percentile<br><br>Summarized across all years within criteria |
| Diel activity | | | | |
| Annual Diurnality index | ((Total movement distance during day / daylength) - (Total movement distance during night / nightlength)) / ((Total movement distance during day / daylength) + (Total movement distance during night / nightlength)) | hourly | Calculated for days with minimum 19 locations | Mean, median, CV, 5th and 95th percentile<br><br>Summarized for entire track |
| Range size | | | | |
| Daily | 95% Minimum | hourly | 12 locations/day | Mean, median, CV, 5th |

| | | | | |
|---|---|---|---|---|
| | Convex Polygon | | | and 95$^{th}$ percentile<br><br>Summarized across all days within criteria |
| Weekly | 95% Minimum Convex Polygon | hourly | 84 hourly relocations (3.5 days) | Mean, median, CV, 5$^{th}$ and 95$^{th}$ percentile<br><br>Summarized across all weeks within criteria |
| Monthly | 95% Minimum Convex Polygon | daily | 14 days of data | Mean, median, CV, 5$^{th}$ and 95$^{th}$ percentile<br><br>Summarized across all months within criteria |
| Annual | 95% Minimum Convex Polygon | weekly | 36 weeks of data | Mean, median, CV, 5$^{th}$ and 95$^{th}$ percentile<br><br>Summarized across all years within criteria |
| Range use/Tortuosity | | | | |
| Intensity of area use - daily | Cumulative hourly movement distance during 24 h, divided by the square root of the range size during the same 24 h | hourly | 12 locations/day | Mean, median, CV, 5$^{th}$ and 95$^{th}$ percentile |
| Intensity of area use - monthly | Cumulative daily movement distance during a given month, divided by the square root of the range size during the same month | daily | 14 days of data | Mean, median, CV, 5$^{th}$ and 95$^{th}$ percentile |
| Intensity of area use - annual | Cumulative weekly movement distance during a given year, divided by the square root of the range size during the same year | weekly | 36 weeks of data | Mean, median, CV, 5$^{th}$ and 95$^{th}$ percentile |

*Table S2: Suggested metadata to be included in future versions of a living animal movement trait database*

| **Metadata** | **Level** |
|---|---|
| Data owner | Study |
| Data owner contact details | Study |
| Study identifier (DOI) | Study |
| Licence type | Study |
| Citation | Study |
| Individual ID | Individual |
| Species | Individual |
| Species common name | Individual |
| Sex | Individual |
| Age [years, juvenile, adult] | Individual |
| Body mass [g] | Individual |
| Tracking start and end date | Individual |
| Most common sampling interval | Individual |
| Mean location - entire track | Individual |
| Mean location - daily, weekly, monthly, annual, either from the observed data or placed in a reference grid cell | Intra-individual |
| Movement mode [walk, fly, swim, arboreal] | Species |
| Sample size [n tracked individuals] | Species |
| Class (mammal, bird) | Species |

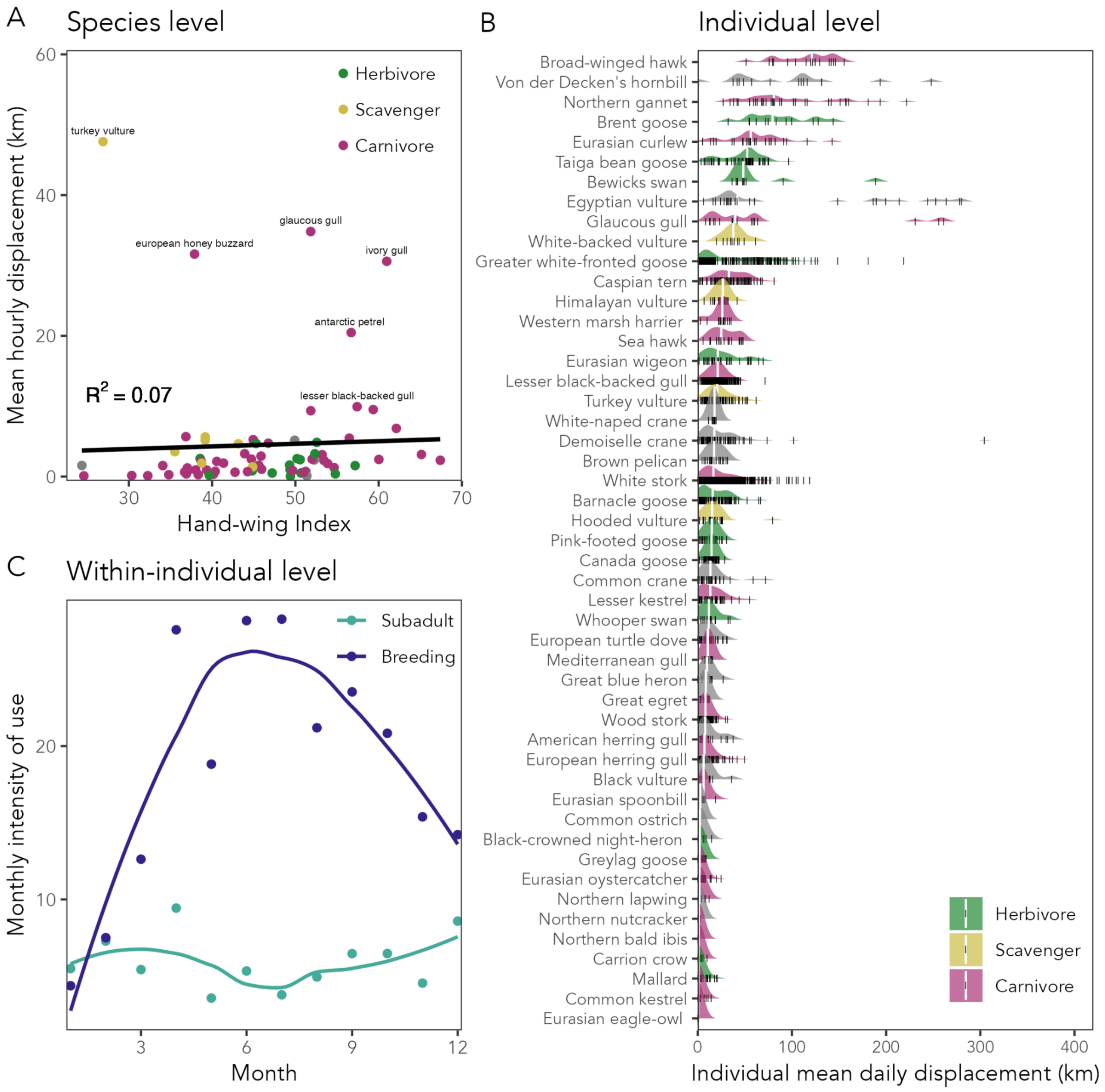


**Figure S1:** Illustrative case study examples for bird species in our MoveTraits database. (A) Hand-wing index is a common proxy for dispersal potential. Using 76 species for which hourly displacements were observed, the hand-wing index only correlated weekly with hourly displacements ($R^2$ = 0.07). (B) Between-individual variation in daily displacement across 49 bird species. Species are coloured according to their trophic position, as noted in the AVONET trait database. Gray shaded species were missing information for trophic position. (C) The monthly intensity of use for one Eurasian griffon vulture (*Gyps fulvus*) individual measures as a subadult in 2017 (green) and as a breeding adult in 2023 (purple). The monthly intensity of use is a measure of space use clusteredness. The individual shows more directed movements throughout the year as a subadult (green) while its breeding status is revealed through clustered movements throughout the summer (purple). Purple and green lines represent smoothers for each respective year over the twelve-month gradient. Data were sourced from the Movebank study ‘Raptors NABU Moessingen public’ (Study ID 186178781, Table S3).

#### Supplementary methods S1: Detailed description of database workflow

We have built a first proof-of-concept database for birds and mammals from open-access Movebank studies (licence types CC_0, CC_BY, CC_BY_NC, see below) and from an open-access dataset published alongside Tucker et al. (2023) and publicly available for download under: https://zenodo.org/records/7704108 (file 'Tucker_Road_Spatial.rds'). The workflow to reproduce this database and the MoveTraits database version 0.1.0 at the species, individual, and intra-individual level has been published under the Open Science Framework (https://doi.org/10.17605/OSF.IO/SP8Z6).

##### *Licence types*

We downloaded publicly available data from Movebank studies with the following open access licence types:

| Licence type | Licence terms |
|---|---|
| CC_0 | • No Copyright<br>• The person who associated a work with this deed has dedicated the work to the public domain by waiving all of his or her rights to the work worldwide under copyright law, including all related and neighbouring rights, to the extent allowed by law.<br>• You can copy, modify, distribute and perform the work, even for commercial purposes, all without asking permission. See Other Information below.<br>• Other Information:<br>○ In no way are the patent or trademark rights of any person affected by CC0, nor are the rights that other persons may have in the work or in how the work is used, such as publicity or privacy rights.<br>○ Unless expressly stated otherwise, the person who associated a work with this deed makes no warranties about the work, and disclaims liability for all uses of the work, to the fullest extent permitted by applicable law.<br>○ When using or citing the work, you should not imply endorsement by the author or the affirmer.<br>• https://creativecommons.org/publicdomain/zero/1.0/ |
| CC_BY | • You are free to:<br>○ Share — copy and redistribute the material in any medium or format for any purpose, even commercially.<br>○ Adapt — remix, transform, and build upon the material for any purpose, even commercially.<br>• The licensor cannot revoke these freedoms as long as you follow the licence terms.<br>• Under the following terms:<br>○ Attribution — You must give appropriate credit, provide a link to the licence, and indicate if changes were made. You may do so in any reasonable manner, but not in any way that suggests the licensor endorses you or your use.<br>○ No additional restrictions — You may not apply legal terms or technological measures that legally restrict others from doing anything the licence permits.<br>• https://creativecommons.org/licenses/by/4.0/ |

| | |
|---|---|
| CC_BY_NC | • You are free to:<br>○ Share — copy and redistribute the material in any medium or format<br>○ Adapt — remix, transform, and build upon the material<br>• The licensor cannot revoke these freedoms as long as you follow the licence terms.<br>• Under the following terms:<br>○ Attribution — You must give appropriate credit, provide a link to the licence, and indicate if changes were made . You may do so in any reasonable manner, but not in any way that suggests the licensor endorses you or your use.<br>○ NonCommercial — You may not use the material for commercial purposes.<br>○ No additional restrictions — You may not apply legal terms or technological measures that legally restrict others from doing anything the licence permits.<br>• https://creativecommons.org/licenses/by-nc/4.0/ |

We contacted all data owners about the use of their data in the MoveTraits database with the option to remove their data from the database. Traits from two studies were removed.

***Workflow details***

1. The workflow for all biologging data downloaded from Movebank entailed data download and cleaning (e.g., removal of records without location coordinates or necessary metadata such as taxon identity and filtering of duplicates, e.g., in cases where tracks of some individuals are uploaded under different study identifiers). Data downloaded from Tucker et al. (2023) was already cleaned. Given the different structure of the two data sources (individual 'move2' objects (Kranstauber *et al.* 2024) for Movebank and one dataframe for (Tucker *et al.* 2023), traits were processed separately and trait databases derived from the two sources were merged in the end.
2. We resampled each individual track to a 1 hour ('hourly') relocation interval with a 15 minute tolerance, to a 24 hour ('daily') relocation interval with a 1 hour tolerance, and to a weekly (168 hours) relocation interval with a 1 day tolerance using the 'amt' R package (Signer *et al.* 2019). Resampling was not, at this stage, stratified to a specific minute of the hour (for hourly resampling) or to a specific hour of the day (for daily resampling). The coordinate reference system was WGS84 (EPSG 4326) unless specified otherwise.
3. We extracted displacement distances (hourly and daily) in meters from the resampled track using the step_lengths() function in the amt package. For maximum displacement distances (daily, weekly, annual) we calculated pairwise Euclidean distances within the focal time frame using the st_distance() function from the 'sf' R package (Pebesma & Bivand 2023). For range size calculation (daily, weekly, monthly, annual) we transformed coordinates to the Mollweide

equal area projection. We used the mcp() function in the R package 'adehabitatHR' (Calenge 2006) to calculate the 95% Minimum Convex Polygon of all points in the focal time period. We extracted the range size in $m^2$. To quantify intensity of use, we used the resampled data (e.g., hourly, daily, weekly) to calculate the cumulative distance during the focal time period (e.g. during a given month) and divided these by the area used during the same period (e.g., MCP during the same week/month). Diurnality was currently exclusively calculated for individuals with hourly data and only for days with at least 20 hourly locations. In the future, diurnality calculations from coarser intervals of two or three hours might be implemented.

4. We summarized traits at the individual level as mean, median, coefficient of variation (CV), $5^{th}$ and $95^{th}$ percentile of the underlying individual movement traits. We summarized these individual level traits at the species level as the mean of each summary (e.g., the mean of all individual $95^{th}$ percentiles). We also provide a version of the database including the underlying individual movement traits (e.g., range size for every month the individual was tracked) alongside the individual's trait summary (e.g., mean monthly range size).
5. We finally merged the databases derived from Movebank data and from the Tucker et al. (2023) study and excluded duplicate individuals from the Tucker et al. (2023) database because it only contained traits based on three months of data (1 February - 15 May 2019).
6. We excluded any records from reptiles and fish as we were uncertain whether our traits would be informative for these taxonomic classes.
