## Supplementary Materials Table S3 for "MoveTraits – A database for integrating animal behaviour into trait-based ecology"

**Table S3.** Overview of studies and their data owners included in the MoveTraits database version 0.1.0. Licence types are explained in the Supplementary methods 1. All raw data were standardised to hourly, daily, and weekly relocation intervals. Fixes at finer intervals were discarded. Traits were only calculated for datasets that fulfilled the minimum data requirements (Table S1). Studies listed informed at least one trait metric with at least one individual.

| Source | Study ID | Species | Principal Investigator | Contact person | Citation | Recommended reference | License type |
|---|---|---|---|---|---|---|---|
| Movebank | 82684 | Bycanistes bucinator | Wolfgang Fiedler | Wolfgang Fiedler | Lenz J, Fiedler W, Caprano T, Friedrichs W, Gaese BH, Wikelski M, Böhning-Gaese K. 2011. Seed-dispersal distributions by trumpeter hornbills in fragmented landscapes. Proceedings of the Royal Society B Biological Sciences. https://doi.org/10.1098/rspb.2010.2383 | https://doi.org/10.1098/rspb.2010.2383 | CC_BY |
| Movebank | 446579 | Anas platyrhynchos | Martin Wikelski | Wolfgang Fiedler | Korner P, Sauter A, Fiedler W, Jenni L. 2016. Variable allocation of activity to daylight and night in the mallard. Animal Behaviour. 115: 69–79. https://doi.org/10.1016/j.anbehav.2016.02.026 | https://doi.org/10.1016/j.anbehav.2016.02.026 | CC_BY |
| Movebank | 481458 | Cathartes aura,Coragyps atratus | David Barber | David Barber | Bildstein KL, Barber D, Bechard MJ, Graña Grilli M, Therrien J. 2021. Data from: Study "Vultures Acopian Center USA GPS" (2003-2021). Movebank Data Repository. <a href="https://www.doi.org/10.5441/001/1.f3qt46r2" target="_blank">https://www.doi.org/10.5441/001/1.f3qt46r2</a> <br><br> Mallon JM, Bildstein KL, Fagan WF. 2021. Inclement weather forces stopovers and prevents migratory progress for obligate soaring migrants. Mov Ecol. 9:39. https://doi.org/10.1186/s40462-021-00274-6 <br><br> Graña Grilli M, Lambertucci SA, Therrien J-F, Bildstein KL. 2017. Wing size but not wing shape is related to migratory behavior in a soaring bird. J Avian Biol. 48(5):669-678. https://doi.org/10.1111/jav.01220 <br><br> Dodge S, Bohrer G, Bildstein K, Davidson SC, Weinzierl R, Mechard MJ, Barber D, Kays R, Brandes D, Han J, et al. 2014. Environmental drivers of variability in the movement ecology of turkey vultures (Cathartes aura) in North and South America. Philos T Roy Soc B. 369(1643):20130195. https://doi.org/10.1098/rstb.2013.0195 <br><br> Portions of these data are published with DOIs 10.5441/001/1.37r2b884 (study "Vultures Acopian Center USA 2003-2016") and 10.5441/001/1.46ft1k05 (study "Turkey vultures in North and South America (data from Dodge et al. 2014)") | https://doi.org/10.5441/001/1.f3qt46r2 | CC_BY |

| Source | Study ID | Species | Principal Investigator | Contact person | Citation | Recommended reference | License type |
|---|---|---|---|---|---|---|---|
| Movebank | 1764627 | Syncerus caffer | Paul Cross | Paul Cross | Cross PC, Bowers JA, Hay CT, Wolhuter J, Buss P, Hofmeyr M, du Toit JT, Getz WM. 2016. Data from: Nonparameteric kernel methods for constructing home ranges and utilization distributions. Movebank Data Repository. <a href="https://www.doi.org/10.5441/001/1.j900f88t" target="_blank">https://www.doi.org/10.5441/001/1.j900f88t</a> <br><br> Calabrese JM, Fleming CH, Gurarie E. 2016. ctmm: an r package for analyzing animal relocation data as a continuous-time stochastic process. Methods Ecol Evol. 7(9):1124-1132. https://doi.org/10.1111/2041-210X.12559 <br><br> Cross PC, Heisey DM, Bowers JA, Hay CT, Wolhuter J, Buss P, Hofmeyr M, Michel AL, Bengis RG, Bird TLF, et al. 2009. Disease, predation and demography: assessing the impacts of bovine tuberculosis on African buffalo by monitoring at individual and population levels. J Appl Ecol. 46(2):467-475. https://doi.org/10.1111/j.1365-2664.2008.01589.x <br><br> Getz WM, Fortmann-Roe S, Cross PC, Lyons AJ, Ryan SJ, Wilmers CC. 2007. LoCoH: Nonparameteric kernel methods for constructing home ranges and utilization distributions. PLoS ONE. 2(2):e207. https://doi.org/10.1371/journal.pone.0000207 | https://doi.org/10.5441/001/1.j900f88t | CC_0 |
| Movebank | 1818825 | Loxodonta cyclotis | Stephen Blake | Stephen Blake | | Movebank study 'Forest Elephant Telemetry Programme'. Accessed on [date]. | CC_BY |
| Movebank | 1902221 | Pecari tajacu | Roland Kays | Roland Kays | No papers published | Movebank study 'Collared Peccary on BCI'. Accessed on [date]. | CC_BY |
| Movebank | 2919708 | Gyps africanus,Torgos tracheliotus | Ran Nathan | Orr Spiegel | Spiegel O, Getz WM, Nathan R. 2014. Data from: Factors influencing foraging search efficiency: Why do scarce lappet-faced vultures outperform ubiquitous white-backed vultures? (V2). Movebank Data Repository. <a href="https://www.doi.org/10.5441/001/1.mf903197" target="_blank">https://www.doi.org/10.5441/001/1.mf903197</a> <br><br> Spiegel O, Getz WM, Nathan R. 2013. Factors influencing foraging search efficiency: why do scarce lappet-faced vultures outperform ubiquitous white-backed vultures? Am Nat. 181(5):E102-115. https://www.doi.org/10.1086/670009 | https://doi.org/10.5441/001/1.mf903197 | CC_0 |
| Movebank | 2927282 | Anas platyrhynchos,Anas penelope,Aythya ferina | Wolfgang Fiedler | Wolfgang Fiedler | Data by Max Planck Institute for Animal Behavior, Vogelwarte Radolfzell, Germany. | Movebank study 'Lake Constance Ducks'. Accessed on [date]. | CC_BY |

| Source | Study ID | Species | Principal Investigator | Contact person | Citation | Recommended reference | License type |
|---|---|---|---|---|---|---|---|
| Movebank | 2930072 | Steatornis caripensis | Martin Wikelski | Martin Wikelski | Holland RA, Wikelski M, Kuemmeth F, Bosque C. 2012. Data from: The secret life of oilbirds: new insights into the movement ecology of a unique avian frugivore. Movebank Data Repository. <a href="https://www.doi.org/10.5441/001/1.35fs26kq" target="_blank">https://www.doi.org/10.5441/001/1.35fs26kq</a> <br><br> Holland RA, Wikelski M, Kuemmeth F, Bosque C. 2009. The secret life of oilbirds: new insights into the movement ecology of a unique avian frugivore. PLoS ONE. 4(12):e8264. https://doi.org/10.1371/journal.pone.0008264 | https://doi.org/10.5441/001/1.35fs26kq | CC_0 |
| Movebank | 2943485 | Aquila chrysaetos,Haliaeetus leucocephalus,Pandion haliaetus,Falco peregrinus,Asio flammeus | Roland Kays | Roland Kays | Nye P, Hewitt G, Swenson T, Kays R. 2018. Data from: New York State bald eagle report 2010. Movebank Data Repository. <a href="https://www.doi.org/10.5441/001/1.s65q50j0" target="_blank">https://www.doi.org/10.5441/001/1.s65q50j0</a> <br><br> Li Z, Han J, Ding B, Kays R. 2012. Mining periodic behaviors of object movements for animal and biological sustainability studies. Dat Min Knowl Disc. 24(2):355-386. https://doi.org/10.1007/s10618-011-0227-9 <br><br> Nye P. 2010. New York State bald eagle report 2010. Albany (NY): New York State Department of Environmental Conservation. 43 p. <br><br> Martell MS, Henny CJ, Nye PE, Solensky MJ. 2001. Fall migration routes, timing, and wintering sites of North American ospreys as determined by satellite telemetry. Condor. 103(4):715-724. https://doi.org/10.1650/0010-5422(2001)103[0715:FMRTAW]2.0.CO;2 <br><br> Rodriguez F, Martell M, Nye P, Bildstein KL. 2001. Osprey migration through Cuba. In: Bildstein K, Klem D Jr, editors. Hawkwatching in the Americas. North Wales (PA): Hawk Migration Association of North America. p. 107-117. <br><br> Bautz L, Nye P. 1987. Spring movement of an adult bald eagle from southeastern New York to central Ontario. The Eyas. 10(1):32-33. | https://doi.org/10.5441/001/1.s65q50j0 | CC_0 |
| Movebank | 2980294 | Anas strepera | Andrea Gehrold | Wolfgang Fiedler | Gehrold A, Bauer H-G, Fiedler W, Wikelski M. 2014. Great flexibility in autumn movement patterns of European Gadwalls (Anas strepera). Journal of Avian Biology. 45: 131–139. https://doi.org/10.1111/j.1600-048X.2013.00248.x<br><br>A subset of these data have been published as<br> Gehrold A, Wikelski M. 2013. Data from: Great flexibility in autumn movement patterns of European Gadwalls (Anas strepera). Movebank Data Repository. <a href="https://doi.org/10.5441/001/1.26dg08hv" target="_blank">https://doi.org/10.5441/001/1.26dg08hv</a> | https://doi.org/10.5441/001/1.26dg08hv | CC_BY |
| Movebank | 2988309 | Phalacrocorax carbo | Wolfgang Fiedler | Wolfgang Fiedler | | Movebank study 'Great Cormorant Lake Constance MPIAB'. Accessed on [date]. | CC_BY |

| Source | Study ID | Species | Principal Investigator | Contact person | Citation | Recommended reference | License type |
|---|---|---|---|---|---|---|---|
| Movebank | 2988333 | Larus fuscus | Martin Wikelski | Martin Wikelski | Wikelski M, Arriero E, Gagliardo A, Holland R, Huttunen MJ, Juvaste R, Mueller I, Tertitski G, Thorup K, Wild M, Alanko M, Bairlein F, Cherenkov A, Cameron A, Flatz R, Hannila J, Hüppop O, Kangasniemi M, Kranstauber B, Penttinen M, Safi K, Semashko V, Schmid H, Wistbacka R. 2015. Data from: True navigation in migrating gulls requires intact olfactory nerves. Movebank Data Repository. <a href="https://doi.org/10.5441/001/1.q986rc29" target="_blank">https://doi.org/10.5441/001/1.q986rc29</a> <br><br> Wikelski M, Arriero E, Gagliardo A, Holand R, Huttunen MJ, Juvaste R, Mueller I, Tertitski G, Thorup K, Wild M, et al. 2015. True navigation in migrating gulls requires intact olfactory nerves. Sci Rep. 5:17061. https://doi.org/10.1038/srep17061 | https://doi.org/10.5441/001/1.q986rc29 | CC_0 |
| Movebank | 3109235 | Anas platyrhynchos | Jonas Waldenström | Jonas Waldenström | Bengtsson, D., Safi, K., Avril, A., Fiedler, W., Wikelski, M., Gunnarsson, G., Elmberg, J., Tolf, C., Olsen, B. & Waldenström, J. 2016. Does influenza A virus infection affect movement behaviour during stopover in its wild reservoir host? Royal Society Open Science 3: 150633. [http://dx.doi.org/10.1098/rsos.150633]. <br /><br /> Bengtsson, D., Avril, A., Gunnarsson, G., Elmberg, J., Söderquist, P., Norevik, G., Tolf, C., Safi, K., Fiedler, W., Wikelski, M., Olsen, B. & Waldenström, J. 2014. Movements, home-range size and habitat selection of Mallards during fall migration. PLOS ONE 9: 00764 [doi: 10.1371/journal.pone.0100764]<br /><br /> | https://doi.org/10.1098/rsos.150633 | CC_BY_NC |
| Movebank | 3780829 | Bubo bubo | Martin Wikelski | Wolfgang Fiedler | Reinhard Vohwinkel et al. in prep. | Movebank study 'Eagle owl Reinhard Vohwinkel MPIAB'. Accessed on [date]. | CC_BY |
| Movebank | 4124813 | Grus nigricollis | Sherub Sherub | Sherub Sherub | Sherub S, Wikelski M. 2024. Data from: Study "Black-necked crane Bhutan (UWICE-MPIAB)". Movebank Data Repository. <a href="https://doi.org/10.5441/001/1.601" target="_blank">https://doi.org/10.5441/001/1.601</a> <br><br> These data are described in<br> Yanco SW, Oliver RY, Iannarilli F, Carlson BS, Heine G, Mueller U, Richter N, Vorneweg B, Andryushchenko Y, Batbayar N, et al. 2024. Migratory birds modulate niche tradeoffs in rhythm with seasons and life history. Proc Natl Acad Sci USA. https://doi.org/10.1073/pnas.2316827121 | https://doi.org/10.5441/001/1.601 | CC_BY |
| Movebank | 6770990 | Fregata magnificens | Martin Wikelski | Martin Wikelski | Wikelski et al., birds and hurricanes, in prep. | Movebank study 'MPIAB PNIC hurricane frigate tracking'. Accessed on [date]. | CC_BY |
| Movebank | 6925808 | Martes pennanti | Scott LaPoint | Scott LaPoint | Paper: LaPoint, S, Gallery P, Wikelski M, Kays R (2013) Animal behavior, cost-based corridor models, and real corridors. Landscape Ecology, v 28 i 8, p 1615–1630. doi:10.1007/s10980-013-9910-0 LaPoint S, Gallery P, Wikelski M, Kays R. 2013. Data from: Animal behavior, cost-based corridor models, and real corridors. Movebank Data Repository. https://doi.org/10.5441/001/1.2tp2j43g | https://doi.org/10.5441/001/1.2tp2j43g | CC_BY_NC |
| Movebank | 7002955 | Ciconia ciconia | Ran Nathan | Shay Rotics | Rotics et al. in prep | Movebank study 'HUJ MPIAB White Stork GSM E-Obs'. Accessed on [date]. | CC_BY |

| Source | Study ID | Species | Principal Investigator | Contact person | Citation | Recommended reference | License type |
|---|---|---|---|---|---|---|---|
| Movebank | 7023252 | Papio anubis | Meg Crofoot | Meg Crofoot | Crofoot MC, Kays RW, Wikelski M. 2021. Data from: Study "Collective movement in wild baboons". Movebank Data Repository. <a href="https://www.doi.org/10.5441/001/1.3q2131q5" target="_blank">https://www.doi.org/10.5441/001/1.3q2131q5</a> <br><br> Harel R, Loftus JC, Crofoot MC. 2021. Locomotor compromises maintain group cohesion in baboon troops on the move. P Roy Soc B. 288(1955):20210839. https://doi.org/10.1098/rspb.2021.0839 <br><br> Farine DR, Strandburg-Peshkin A, Couzin ID, Berger-Wolf TY, Crofoot MC. 2017. Individual variation in local interaction rules can explain emergent patterns of spatial organization in wild baboons. P Roy Soc B. 284(1853):20162243. https://doi.org/10.1098/rspb.2016.2243 <br><br> Strandburg-Peshkin A, Farine DR, Crofoot MC, Couzin ID. 2017. Habitat and social factors shape individual decisions and emergent group structure during baboon collective movement. eLife. 6:e19505. https://doi.org/10.7554/eLife.19505 <br><br> Farine DR, Strandburg-Peshkin A, Berger-Wolf T, Ziebart B, Brugere I, Li J, Crofoot MG. 2016. Both nearest neighbours and long-term affiliates predict individual locations during collective movement in wild baboons. Sci Rep. 6:27704. https://doi.org/10.1038/srep27704 <br><br> Strandburg-Peshkin A, Farine DR, Couzin ID, Crofoot MC. 2015. Shared decision-making drives collective movement in wild baboons. Science. 348–6241:1358–1361. https://doi.org/10.1126/science.aaa5099 | https://doi.org/10.5441/001/1.3q2131q5 | CC_0 |
| Movebank | 7431347 | Ciconia ciconia | Movebank Administrator | Wolfgang Fiedler | Berthold P, Kaatz C, Kaatz M, Querner U, van den Bossche W, Chernetsov N, Fiedler W, Wikelski M. 2022. Data from: Study "MPIAB Argos white stork tracking (1991-2017)". Movebank Data Repository. <a href="https://www.doi.org/10.5441/001/1.k29d81dh" target="_blank">https://www.doi.org/10.5441/001/1.k29d81dh</a> | https://doi.org/10.5441/001/1.k29d81dh | CC_BY |
| Movebank | 8019591 | Bison bison | Stephen Blake | Stephen Blake | | Movebank study 'Missouri Bison Tracking Project'. Accessed on [date]. | CC_BY |
| Movebank | 8849813 | Ardea alba | Roland Kays | Roland Kays | Brzorad, J. N., M. C. Allen, S. Jennings, E. Condeso, S. Elbin, R. Kays, D. Lumpkin, S. Schweitzer, N. Tsipoura, and A. D. Maccarone. 2022. Seasonal Patterns in Daily Flight Distance and Space Use by Great Egrets (Ardea alba). Waterbirds 44:343–355. | https://doi.org/10.1675/063.044.0309 | CC_0 |
| Movebank | 8927992 | Anas platyrhynchos,Fulica atra | Martin Wikelski | Martin Wikelski | Wikelski et al., unpublished | Movebank study 'LifeTrack - Evros Delta'. Accessed on [date]. | CC_BY |
| Movebank | 9493881 | Ciconia ciconia | Martin Wikelski | Wolfgang Fiedler | Data from study "LifeTrack White Stork Uzbekistan" in www.movebank.org by Max Planck institute of Animal Behavior (Radolfzell, Germany) and cooperation partners. | Movebank study 'LifeTrack White Stork Uzbekistan'. Accessed on [date]. | CC_BY |

| Source | Study ID | Species | Principal Investigator | Contact person | Citation | Recommended reference | License type |
|---|---|---|---|---|---|---|---|
| Movebank | 9651291 | Neophron percnopterus | Evan Buechley | Evan Buechley | Buechley ER, Şekercioğlu CH. 2019. Data from: Satellite tracking a wide-ranging endangered vulture species to target conservation actions in the Middle East and East Africa. Movebank Data Repository. <a href="https://www.doi.org/10.5441/001/1.385gk270" target="_blank">https://www.doi.org/10.5441/001/1.385gk270</a> <br><br> Phipps WL, López-López P, Buechley ER, Oppel S, Álvarez E, Arkumarev V, Bekmansurov R, Berger-Tal O, Bermejo A, Bounas A, Alanís IC, de la Puente J, Dobrev V, Duriez O, Efrat R, Fréchet G, García J, Galán M, García-Ripollés C, Gil A, Iglesias-Lebrija JJ, Jambas J, Karyakin IV, Kobierzycki E, Kret E, Loercher F, Monteiro A, Etxebarria JM, Nikolov SC, Pereira J, Peške L, Ponchon C, Realinho E, Saravia V, Şekercioğlu ÇH, Skartsi T, Tavares J, Teodósio J, Urios V, Vallverdú N. 2019. Spatial and temporal variability in migration of a soaring raptor across three continents. Front Ecol Evol. 7:323. https://doi.org/10.3389/fevo.2019.00323 <br><br> Buechley ER, McGrady MJ, Çoban E, Şekercioğlu ÇH. 2018. Satellite tracking a wide-ranging endangered vulture species to target conservation actions in the Middle East and East Africa. Biodivers Conserv. 27(9):2293-2310. https://doi.org/10.1007/s10531-018-1538-6 <br><br> Buechley ER, Oppel S, Beatty WS, Nikolov SC, Dobrev V, Arkumarev V, Saravia V, Bougain C, Bounas A, Kret E, Skartsi T, Aktay L, Ahababyan K, Frehner E, Şekercioğlu ÇH. 2018. Identifying critical migratory bottlenecks and high-use areas for an endangered migratory soaring bird across three continents. J Avian Biol. 49(7):e01629. https://doi.org/10.1111/jav.01629 | https://doi.org/10.5441/001/1.385gk270 | CC_BY |
| Movebank | 10006517 | Uria lomvia | Grant Gilchrist | Kyle Elliott | | Movebank study 'Thick-billed Murres; Gilchrist; Digges Island, Canada'. Accessed on [date]. | CC_0 |
| Movebank | 10157679 | Ciconia ciconia | Azafzaf.H | Wolfgang Fiedler | Azafzaf, H., Feltrup-Azafzaf, C., Flack, A., Wikelski, M. & Fiedler, W. (year of access): GPS Tracking data from study "LifeTrack White Stork Tunisia" in www.movebank.org.. | https://doi.org/10.1126/sciadv.1500931 | CC_BY |
| Movebank | 10204361 | Pandion haliaetus | Barb Jensen | Barb Jensen | | Movebank study 'Pandion haliaetus Osprey - SouthEast Michigan'. Accessed on [date]. | CC_0 |

| Source | Study ID | Species | Principal Investigator | Contact person | Citation | Recommended reference | License type |
|---|---|---|---|---|---|---|---|
| Movebank | 10236270 | Ciconia ciconia | Wolfgang Fiedler | Wolfgang Fiedler | Parts of this dataset are also available in the study "MPIO white stork lifetime tracking data (2013-2014)" and published as<br> Flack A, Fiedler W, Blas J, Pokrovski I, Mitropolsky B, Kaatz M, Aghababyan K, Khachatryan A, Fakriadis I, Makrigianni E, Jerzak L, Shamin M, Shamina C, Azafzaf H, Feltrup-Azafzaf C, Mokotjomela TM, Wikelski M. 2015. Data from: Costs of migratory decisions: a comparison across eight white stork populations. Movebank Data Repository. <a href="https://www.doi.org/10.5441/001/1.78152p3q" target="_blank">https://www.doi.org/10.5441/001/1.78152p3q</a> <br><br> Flack A, Fiedler W, Blas J, Pokrovski I, Kaatz M, Mitropolsky M, Aghababyan K, Fakriadis Y, Makrigianni E, Jerzak L, et al. 2016. Costs of migratory decisions: a comparison across eight white stork populations. Science Advances. 2(1): e1500931. https://doi.org/10.1126/sciadv.1500931 <br><br> Kays R, Davidson SC, Berger M, Bohrer G, Fiedler W, Flack A, Hirt J, Hahn C, Gauggel D, Russell B, et al. 2021. The Movebank system for studying global animal movement and demography. Methods Ecol Evol. https://doi.org/10.1111/2041-210X.13767 | https://doi.org/10.1111/2041-210X.13767 | CC_BY |
| Movebank | 10449318 | Ciconia ciconia,Homo sapiens | Martin Wikelski | Wolfgang Fiedler | Data from study "LifeTrack White Stork Loburg" in www.movebank.org by Max Planck Institute of Animal Behavior (Radolfzell, Germany), Hebrew University of Jerusalem (Israel), University of Potsdam (Germany) and Storchenhof Loburg (Germany) | Movebank study 'LifeTrack White Stork Loburg'. Accessed on [date]. | CC_BY |
| Movebank | 10449535 | Ciconia ciconia | Martin Wikelski | vasilis elias | Parts of this dataset are also available in the study "MPIO white stork lifetime tracking data (2013-2014)" and published as<br> Flack A, Fiedler W, Blas J, Pokrovski I, Mitropolsky B, Kaatz M, Aghababyan K, Khachatryan A, Fakriadis I, Makrigianni E, Jerzak L, Shamin M, Shamina C, Azafzaf H, Feltrup-Azafzaf C, Mokotjomela TM, Wikelski M. 2015. Data from: Costs of migratory decisions: a comparison across eight white stork populations. Movebank Data Repository. <a href="https://www.doi.org/10.5441/001/1.78152p3q" target="_blank">https://www.doi.org/10.5441/001/1.78152p3q</a> <br><br> Flack A, Fiedler W, Blas J, Pokrovski I, Kaatz M, Mitropolsky M, Aghababyan K, Fakriadis Y, Makrigianni E, Jerzak L, et al. 2016. Costs of migratory decisions: a comparison across eight white stork populations. Science Advances. 2(1): e1500931. https://doi.org/10.1126/sciadv.1500931<br><br>Kays R, Davidson SC, Berger M, Bohrer G, Fiedler W, Flack A, Hirt J, Hahn C, Gauggel D, Russell B, et al. 2021. The Movebank system for studying global animal movement and demography. Methods Ecol Evol. https://doi.org/10.1111/2041-210X.13767 | https://doi.org/10.1111/2041-210X.13767 | CC_BY |
| Movebank | 10449698 | Ciconia ciconia | Ran Nathan | Shay Rotics | Rotics et al. unpublished data | Movebank study 'HUJ MPIAB White Stork GSM 2013'. Accessed on [date]. | CC_BY |

| Source | Study ID | Species | Principal Investigator | Contact person | Citation | Recommended reference | License type |
|---|---|---|---|---|---|---|---|
| Movebank | 10596067 | Ciconia ciconia | Martin Wikelski | Andrea Flack | Parts of this dataset are also available in the study "MPIO white stork lifetime tracking data (2013-2014)" and published as<br> Flack A, Fiedler W, Blas J, Pokrovski I, Mitropolsky B, Kaatz M, Aghababyan K, Khachatryan A, Fakriadis I, Makrigianni E, Jerzak L, Shamin M, Shamina C, Azafzaf H, Feltrup-Azafzaf C, Mokotjomela TM, Wikelski M. 2015. Data from: Costs of migratory decisions: a comparison across eight white stork populations. Movebank Data Repository. <a href="https://www.doi.org/10.5441/001/1.78152p3q" target="_blank">https://www.doi.org/10.5441/001/1.78152p3q</a> <br><br> Flack A, Fiedler W, Blas J, Pokrovski I, Kaatz M, Mitropolsky M, Aghababyan K, Fakriadis Y, Makrigianni E, Jerzak L, et al. 2016. Costs of migratory decisions: a comparison across eight white stork populations. Science Advances. 2(1): e1500931. https://doi.org/10.1126/sciadv.1500931<br><br>Kays R, Davidson SC, Berger M, Bohrer G, Fiedler W, Flack A, Hirt J, Hahn C, Gauggel D, Russell B, et al. 2021. The Movebank system for studying global animal movement and demography. Methods Ecol Evol. https://doi.org/10.1111/2041-210X.13767 | https://doi.org/10.1111/2041-210X.13767 | CC_BY |
| Movebank | 10722328 | Grus grus | Ramūnas Žydelis | Ramūnas Žydelis | Žydelis R, Desholm M, Månsson J, Nilsson L, Skov H. 2024. Data from: Study "GPS telemetry of Common Cranes, Sweden". Movebank Data Repository. <a href="https://doi.org/10.5441/001/1.597" target="_blank">https://doi.org/10.5441/001/1.597</a> <br><br> These data are described in<br> Yanco SW, Oliver RY, Iannarilli F, Carlson BS, Heine G, Mueller U, Richter N, Vorneweg B, Andryushchenko Y, Batbayar N, et al. 2024. Migratory birds modulate niche tradeoffs in rhythm with seasons and life history. Proc Natl Acad Sci USA. https://doi.org/10.1073/pnas.2316827121 | https://doi.org/10.5441/001/1.597 | CC_BY |
| Movebank | 10763606 | Ciconia ciconia | Martin Wikelski | Wolfgang Fiedler | Flack A, Fiedler W, Blas J, Pokrovsky I, Kaatz M, Mitropolsky M, Aghababyan K, Fakriadis I, Makrigianni E, Jerzak L, et al. 2016. Costs of migratory decisions: A comparison across eight white stork populations. Sci Adv. 2(1):e1500931. https://doi.org/10.1126/sciadv.1500931. | https://doi.org/10.1126/sciadv.1500931 | CC_BY |
| Movebank | 11223924 | Anser fabalis | Tomas Aarvak | Tomas Aarvak | | Movebank study 'Bean Goose Anser fabalis Finnmark.'. Accessed on [date]. | CC_BY |
| Movebank | 11948467 | Didelphis virginiana,Procyon lotor | Roland Kays | Roland Kays | | Movebank study 'NCSU Mammalogy Campus Carnivores'. Accessed on [date]. | CC_0 |
| Movebank | 13978569 | Phalcoboenus australis | David Barber | David Barber | | Movebank study 'Striated Caracara Falkland Islands'. Accessed on [date]. | CC_BY |
| Movebank | 14288429 | Equus hemionus | Petra Kaczensky | Petra Kaczensky | Kaczensky, P., Walzer, C. 2009. Gobi khulan GPS-ARGOS 2002-2008 tracking dataset. | Movebank study 'Khulan Mongolia GPS-ARGOS 2002-2008'. Accessed on [date]. | CC_BY_NC |
| Movebank | 14671003 | Necrosyrtes monachus | David Barber | David Barber | | Movebank study 'Hooded Vulture Africa'. Accessed on [date]. | CC_BY |

| Source | Study ID | Species | Principal Investigator | Contact person | Citation | Recommended reference | License type |
|---|---|---|---|---|---|---|---|
| Movebank | 15661938 | Physeter macrocephalus | Daniel Palacios | Barb Lagerquist | Irvine LM, Follett TM, Winsor MH, Mate BR, Palacios DM. 2020. Data from: Study "Sperm whales Gulf of Mexico 2011-2013 - Argos data". Movebank Data Repository. doi: <a href="https://www.doi.org/10.5441/001/1.tj291471">10.5441/001/1.tj291471</a> <br><br> Irvine LM, Winsor MH, Follett TM, Mate BR, Palacios DM. 2020. An at-sea assessment of Argos location accuracy for three species of large whales, and the effect of deep-diving behavior on location error. Animal Biotelemetry. 8:20. doi: <a href="https://doi.org/10.1186/s40317-020-00207-x">10.1186/s40317-020-00207-x</a> | https://doi.org/10.5441/001/1.44cb3946 | CC_BY |
| Movebank | 16880941 | Cathartes aura | David Barber | David Barber | A more complete version of this dataset is stored in study "Vultures Acopian Center USA GPS".<br><br>Bildstein K, Barber D, Bechard MJ. 2014. Data from: Environmental drivers of variability in the movement ecology of turkey vultures (Cathartes aura) in North and South America. Movebank Data Repository. <a href="https://doi.org/10.5441/001/1.46ft1k05" target="_blank">https://doi.org/10.5441/001/1.46ft1k05</a> <br><br> Dodge S, Bohrer G, Bildstein K, Davidson SC, Weinzierl R, Mechard MJ, Barber D, Kays R, Brandes D, Han J. 2014. Environmental drivers of variability in the movement ecology of turkey vultures (Cathartes aura) in North and South America. Philos T Roy Soc B. 369(1643):20130195. https://doi.org/10.1098/rstb.2013.0195 | https://doi.org/10.5441/001/1.46ft1k05 | CC_0 |
| Movebank | 17469219 | Ardeidae ,Ardea herodias,Egretta thula,Ardea alba | John Brzorad | John Brzorad | | Movebank study 'Egrets & Herons'. Accessed on [date]. | CC_BY_NC |
| Movebank | 19186107 | Fregata magnificens | Patrick Jodice | Patrick Jodice | Jodice, P.G.R., K. Meyer, S. Zaluski, and L. Soanes | Movebank study 'Magnificent Frigatebird_BVI_GPS-PTT_2014-2016'. Accessed on [date]. | CC_BY |

| Source | Study ID | Species | Principal Investigator | Contact person | Citation | Recommended reference | License type |
|---|---|---|---|---|---|---|---|
| Movebank | 19411459 | Panthera onca | Ronaldo Morato | Ronaldo Morato | Morato RG, Kantek DLZ, Miyazaki S, Deluque T, de Paula RC. 2021. Data from: Jaguar movement database: a GPS-based movement dataset of an apex predator in the Neotropics. Movebank Data Repository. <a href="https://www.doi.org/10.5441/001/1.3c4fv0m4" target="_blank">https://www.doi.org/10.5441/001/1.3c4fv0m4</a> <br><br> These data are described in the following written publications:<br><br>Morato RG Thompson JJ, Paviolo A, De la Torre JA, Lima F, McBride Jr RT, De Paula RC, Cullen Jr L, Silveira L, Kantek DLZ, et al. 2018. Jaguar movement database: a GPS-based movement dataset of an apex predator in the Neotropics. Ecology. 99(7):1691. https://doi.org/10.1002/ecy.2379 <br><br> Morato RG, Connette GM, Stabach JA, De Paula RC, Ferraz KMPM, Kantek DLZ, Miyazaki SS, Pereira TDC, Silva LC, Paviolo A, et al. 2018. Resource selection in an apex predator and variation in response to local landscape characteristics. Biol Conserv. 228:233–40. https://doi.org/10.1016/j.biocon.2018.10.022 <br><br> Morato RG, Stabach JA, Fleming CH, Calabrese JM, De Paula RC, Ferraz KMPM, Kantek DLZ, Miyazaki SS, Pereira TDC, Araujo GR, et al. 2016. Space use and movement of a neotropical top predator: the endangered jaguar. PLoS ONE. 11(12):e0168176. https://doi.org/10.1371/journal.pone.0168176 <br><br> In addition, a version of these data are included as part of the following published datasets: <br><br> Morato RG, Thompson JJ, Paviolo A, de la Torre JA, Lima F, McBride Jr RT, Paula RC, Cullen Jr L, Silveira L, Kantek DLZ, et al. 2019. Data from: Jaguar Movement Database: a GPS-based movement dataset of an apex predator in the Neotropics. Dryad. https://doi.org/10.5061/dryad.2dh0223 <br><br> miltinhoastronauta. 2018. LEEClab/jaguar_movement: Jaguar Movement Database v1.0 released! (v1.0). Zenodo. https://doi.org/10.5281/zenodo.1345119 <br><br> https://github.com/LEEClab/jaguar_movement/tree/v1.0 | https://doi.org/10.5441/001/1.3c4fv0m4 | CC_0 |

| Source | Study ID | Species | Principal Investigator | Contact person | Citation | Recommended reference | License type |
|---|---|---|---|---|---|---|---|
| Movebank | 20202974 | Gyps himalayensis | Martin Wikelski | Sherub Sherub | Portions of these data are published as<br> Sherub S and Wikelski M. 2021. Data from: Longer days enable higher diurnal activity for migratory birds [Himalayan griffons]. Movebank Data Repository. <a href="https://www.doi.org/10.5441/001/1.4n2501f5" target="_blank">https://www.doi.org/10.5441/001/1.4n2501f5</a> <br><br> Sherub S, Wikelski M, Fiedler W, Davidson SC. 2016. Data from: Behavioural adaptations to flight into thin air. Movebank Data Repository. https://doi.org/10.5441/001/1.143v2p2k (study "High-altitude flights of Himalayan vultures (data from Sherub et al. 2016)") <br><br> Papers<br> Pokrovsky I, Kölzsch A, Sherub S, Fiedler W, Glazov P, Kulikova O, Wikelski M, Flack A. 2021. Longer days enable higher diurnal activity for migratory birds. J Anim Ecol. https://doi.org/10.1111/1365-2656.13484 <br><br> Sherub S, Bohrer G, Wikelski M, Weinzierl R. 2016. Behavioural adaptations to flight into thin air. Biology Letters. 12(10):20160432. https://doi.org/10.1098/rsbl.2016.0432 | https://doi.org/10.5441/001/1.4n2501f5 | CC_BY |
| Movebank | 20873986 | Haliaeetus leucocephalus | Roland Kays | Roland Kays | Roland Kays and Ted Simons | Movebank study 'LifeTrack Bald Eagle'. Accessed on [date]. | CC_0 |
| Movebank | 21231406 | Ciconia ciconia | Martin Wikelski | Wolfgang Fiedler | Fiedler W, Flack A, Schäfle W, Keeves B, Quetting M, Eid B, Schmid H, Wikelski M. 2024. Data from: Study "LifeTrack White Stork SW Germany" (2013-2023). Movebank Data Repository. <a href="https://doi.org/10.5441/001/1.ck04mn78_2" target="_blank">https://doi.org/10.5441/001/1.ck04mn78_2</a> <br><br> These data are described in<br> Aikens EO, Nourani E, Fiedler W, Wikelski M, Flack A. 2024. Learning shapes the development of migratory behavior. Proc Natl Acad Sci USA. 121(12):e2306389121. https://doi.org/10.1073/pnas.2306389121<br><br>Kays R, Davidson SC, Berger M, Bohrer G, Fiedler W, Flack A, Hirt J, Hahn C, Gauggel D, Russell B, et al. 2022. The Movebank system for studying global animal movement and demography. Methods Ecol Evol. 13(2):419-431. https://doi.org/10.1111/2041-210X.13767 <br><br> Cheng Y, Fiedler W, Wikelski M, Flack A. 2019. "Closer-to-home" strategy benefits juvenile survival in a long-distance migratory bird. Ecology and Evolution. 9(16): 8945–8952. https://doi.org/10.1002/ece3.5395 <br><br> Flack A, Fiedler W, Blas J, Pokrovski I, Kaatz M, Mitropolsky M, Aghababyan K, Fakriadis Y, Makrigianni E, Jerzak L, et al. 2016. Costs of migratory decisions: a comparison across eight white stork populations. Science Advances. 2(1): e1500931. https://doi.org/10.1126/sciadv.1500931 <br><br> Weinzierl R, Bohrer G, Kranstauber B, Fiedler W, Wikelski M, Flack A. 2016. Wind estimation based on thermal soaring of birds. Ecology and Evolution. 6(24): 8706–8718. https://doi.org/10.1002/ece3.2585 <br><br> Parts of this dataset are also available in studies "Fall migration of white storks in 2014" (published as DOI 10.5441/001/1.bj96m274) and "MPIO white stork lifetime tracking data (2013-2014)" (published as DOI 10.5441/001/1.78152p3q). | https://doi.org/10.5441/001/1.ck04mn78_2 | CC_BY |

| Source | Study ID | Species | Principal Investigator | Contact person | Citation | Recommended reference | License type |
|---|---|---|---|---|---|---|---|
| Movebank | 24442409 | Ciconia ciconia | Wolfgang Fiedler | Wolfgang Fiedler | Fiedler W, Leppelsack E, Leppelsack H, Stahl T, Wieding O, Wikelski M. 2024. Data from: Study "LifeTrack White Stork Bavaria" (2014-2023). Movebank Data Repository. <a href="https://doi.org/10.5441/001/1.v1cs4nn0_2" target="_blank">https://doi.org/10.5441/001/1.v1cs4nn0_2</a> <br><br> Aikens EO, Nourani E, Fiedler W, Wikelski M, Flack A. 2024. Learning shapes the development of migratory behavior. Proc Natl Acad Sci USA. 121(12):e2306389121. https://doi.org/10.1073/pnas.2306389121<br><br> Brønnvik H, Nourani E, Fiedler W, Flack A. 2024. Experience reduces reliance on availability of conspecifics for route selection by a collectively migrating soaring bird. Curr Biol. 34(9):2030-2037.e3. https://doi.org/10.1016/j.cub.2024.03.052<br><br> Kays R, Davidson SC, Berger M, Bohrer G, Fiedler W, Flack A, Hirt J, Hahn C, Gauggel D, Russell B, et al. 2022. The Movebank system for studying global animal movement and demography. Methods Ecol Evol. 13(2):419-431. https://doi.org/10.1111/2041-210X.13767<br><br> Cheng Y, Fiedler W, Wikelski M, Flack A. 2019. "Closer-to-home" strategy benefits juvenile survival in a long-distance migratory bird. Ecology and Evolution. 9(16): 8945–8952. https://doi.org/10.1002/ece3.5395 | https://doi.org/10.5441/001/1.v1cs4nn0_2 | CC_BY |
| Movebank | 28691134 | Buteo platypterus | Laurie Goodrich | David Barber | contact PI | Movebank study 'Broad-winged Hawk habitat use, range, and movement ecology'. Accessed on [date]. | CC_BY |

| Source | Study ID | Species | Principal Investigator | Contact person | Citation | Recommended reference | License type |
|---|---|---|---|---|---|---|---|
| Movebank | 29799425 | Branta leucopsis | Klaus-Michael Exo | Michael Exo | van der Jeugd H, Oosterbeek K, Ens BJ, Shamoun-Baranes J, Exo K. 2014. Data from: Forecasting spring from afar? Timing of migration and predictability of phenology along different migration routes of an avian herbivore [Barents Sea data]. Movebank Data Repository. <a href="https://www.doi.org/10.5441/001/1.ps244r11" target="_blank">https://www.doi.org/10.5441/001/1.ps244r11</a> <br><br> Shariati-Najafabadi M et al. 2016. Environmental parameters linked to the last migratory stage of barnacle geese en route to their breeding grounds. Animal Behavior. 118:81-95. https://doi.org/10.1016/j.anbehav.2016.05.018<br><br> Shariati-Najafabadi M et al. 2015. Satellite-versus temperature-derived green wave indices for predicting the timing of spring migration of avian herbivores. Ecological Indicators 58:322–331. https://doi.org/10.1016/j.ecolind.2015.06.005<br><br> Kölzsch A et al. 2015. Forecasting spring from afar? Timing of migration and predictability of phenology along different migration routes of an avian herbivore. J Anim Ecol. 84(1):272-283. https://doi.org/10.1111/1365-2656.12281<br><br> Shariatinajafabadi M et al. 2014. Migratory herbivorous waterfowl track satellite-derived green wave index. PLoS ONE 9(9):e108331. https://doi.org/10.1371/journal.pone.0108331<br><br> Ens BJ et al. 2008. Tracking of individual birds—report on WP 3230 (bird tracking sensor characterization) and WP 4130 (sensor adaptation and calibration for bird tracking system) of the FlySafe basic activities project. SOVON-onderzoeksrapport 2008/10, SOVON Vogelonderzoek Nederland, Beek-Ubbergen 78 p. | https://doi.org/10.5441/001/1.ps244r11 | CC_0 |
| Movebank | 33622846 | Uria lomvia | Grant Gilchrist | Kyle Elliott | | Movebank study 'Thick-billed Murres; Gilchrist; Cape Graham Moore, Canada'. Accessed on [date]. | CC_0 |
| Movebank | 33643212 | Gyps fulvus,Gyps africanus,Torgos tracheliotus | Ran Nathan | Movement Ecology Lab at the Hebrew University of Jerusalem, Israel | Spiegel OM, Harel R, Centeno-Cuadros A, Hatzofe O, Getz WM, Nathan R. 2015. Data from: Moving beyond curve-fitting: using complementary data to assess alternative explanations for long movements of three vulture species. Movebank Data Repository. <a href="https://www.doi.org/10.5441/001/1.8c56f72s" target="_blank">https://www.doi.org/10.5441/001/1.8c56f72s</a> <br><br> Spiegel OM, Harel R, Centeno-Cuadros A, Hatzofe O, Getz WM, Nathan R. 2015. Moving beyond curve-fitting: using complementary data to assess alternative explanations for long movements of three vulture species. Am Nat. 185(2). https://www.doi.org/10.1086/679314 | https://doi.org/10.5441/001/1.8c56f72s | CC_0 |

| Source | Study ID | Species | Principal Investigator | Contact person | Citation | Recommended reference | License type |
|---|---|---|---|---|---|---|---|
| Movebank | 37350671 | Buceros bicornis,Rhyticeros undulatus | Aparajita Datta and Rohit Naniwadekar | Rohit | Naniwadekar R, Rathore A, Shukla U, Chaplod S, Datta A. 2019. Data from: How far do Asian forest hornbills disperse seeds? Movebank Data Repository. <a href="https://www.doi.org/10.5441/001/1.14sm8k1d" target="_blank">https://www.doi.org/10.5441/001/1.14sm8k1d</a> <br><br> Naniwadekar R, Rathore A, Shukla U, Chaplod S, Datta A. 2019. How far do Asian forest hornbills disperse seeds? Acta Oecol. 101:103482. https://doi.org/10.1016/j.actao.2019.103482 | https://doi.org/10.5441/001/1.14sm8k1d | CC_0 |
| Movebank | 40906102 | Lemur catta | Teague O'Mara | Teague O'Mara | O'Mara. 2012. Development of feeding in ring-tailed lemurs. PhD Thesis. Arizona State University. | https://doi.org/http://hdl.handle.net/2286/R.A.93648 | CC_0 |
| Movebank | 42451582 | Numenius americanus | Jay Carlisle | Jay Carlisle | | Movebank study 'Long-billed Curlew Migration from the Intermountain West'. Accessed on [date]. | CC_BY_NC |
| Movebank | 47450376 | Grus antigone | Robert van Zalinge | Robert van Zalinge | | Movebank study 'Sarus Crane, van Zalinge, Cambodia'. Accessed on [date]. | CC_BY |
| Movebank | 53460105 | Papio anubis | Meg Crofoot | Meg Crofoot | Crofoot MC, Kays RW, Wikelski M. 2015. Data from: Shared decision-making drives collective movement in wild baboons. Movebank Data Repository. <a href="https://www.doi.org/10.5441/001/1.kn0816jn" target="_blank">https://www.doi.org/10.5441/001/1.kn0816jn</a> <br><br> Strandburg-Peshkin A, Farine DR, Couzin ID, Crofoot MC. 2015. Shared decision-making drives collective movement in wild baboons. Science. 348–6241: 1358–1361. https://doi.org/10.1126/science.aaa5099 <br><br> A larger dataset containing the data in this study is available as https://doi.org/10.5441/001/1.3q2131q5 and in the study "Collective [illegible] | https://doi.org/10.5441/001/1.kn0816jn | CC_BY |
| Movebank | 67281010 | Mycteria americana | Roland Kays | Roland Kays | Schweitzer S, Bryan AL Jr, Brzorad J, Kays R. 2023. Data from: Study "NC Wood Stork Tracking". Movebank Data Repository. <a href="https://doi.org/10.5441/001/1.303" target="_blank">https://doi.org/10.5441/001/1.303</a> | Movebank study 'NC Wood Stork Tracking'. Accessed on [date]. | CC_BY |
| Movebank | 69724677 | Branta bernicla,Branta leucopsis | Stefan Garthe | Stefan Garthe | | Movebank study 'FTZ Geese Wadden Sea'. Accessed on [date]. | CC_0 |
| Movebank | 74496970 | Ciconia ciconia | Andrea Flack | Andrea Flack | Flack A, Fiedler W, Blas J, Pokrovski I, Mitropolsky B, Kaatz M, Aghababyan K, Khachatryan A, Fakriadis I, Makrigianni E, et al. 2015. Data from: Costs of migratory decisions: a comparison across eight white stork populations. Movebank Data Repository. <a href="https://www.doi.org/10.5441/001/1.78152p3q" target="_blank">https://www.doi.org/10.5441/001/1.78152p3q</a> <br><br> Flack A, Fiedler W, Blas J, Pokrovski I, Kaatz M, Mitropolsky M, Aghababyan K, Fakriadis Y, Makrigianni E, Jerzak L, et al. 2016. Costs of migratory decisions: a comparison across eight white stork populations. Science Advances. 2(1): e1500931. https://doi.org/10.1126/sciadv.1500931<br><br>Kays R, Crofoot MC, Jetz W, Wikelski M. 2015. Terrestrial animal tracking as an eye on life and planet. Science. 348(6240):aaa2478. https://doi.org/10.1126/science.aaa2478 | https://doi.org/10.5441/001/1.78152p3q | CC_BY |

| Source | Study ID | Species | Principal Investigator | Contact person | Citation | Recommended reference | License type |
|---|---|---|---|---|---|---|---|
| Movebank | 76367850 | Ciconia ciconia,Homo sapiens | Wolfgang Fiedler | Wolfgang Fiedler | Fiedler W, Hilsendegen C, Reis C, Lehmann J, Hilsendegen P, Schmid H, Wikelski M. 2024. Data from: Study "LifeTrack White Stork Rheinland-Pfalz" (2015-2023). Movebank Data Repository. <a href="https://doi.org/10.5441/001/1.4192t2j4_2" target="_blank">https://doi.org/10.5441/001/1.4192t2j4_2</a> <br><br>Aikens EO, Nourani E, Fiedler W, Wikelski M, Flack A. 2024. Learning shapes the development of migratory behavior. Proc Natl Acad Sci USA. 121(12):e2306389121. https://doi.org/10.1073/pnas.2306389121<br><br>Brønnvik H, Nourani E, Fiedler W, Flack A. 2024. Experience reduces reliance on availability of conspecifics for route selection by a collectively migrating soaring bird. Curr Biol. 34(9):2030-2037.e3. https://doi.org/10.1016/j.cub.2024.03.052<br><br>Kays R, Davidson SC, Berger M, Bohrer G, Fiedler W, Flack A, Hirt J, Hahn C, Gauggel D, Russell B, et al. 2022. The Movebank system for studying global animal movement and demography. Methods Ecol Evol. 13(2):419-431. https://doi.org/10.1111/2041-210X.13767<br><br>Cheng Y, Fiedler W, Wikelski M, Flack A. 2019. "Closer-to-home" strategy benefits juvenile survival in a long-distance migratory bird. Ecology and Evolution. 9(16): 8945–8952. https://doi.org/10.1002/ece3.5395 | https://doi.org/10.5441/001/1.4192t2j4_2 | CC_BY |
| Movebank | 79206236 | Ciconia ciconia | Martin Wikelski | Wolfgang Fiedler | Grasso R, Gagliardo A, Zafarana M, Müller I, Schmid H, Fiedler W, Wikelski M. 2021. Data from: Study "LifeTrack White Stork Sicily". Movebank Data Repository. <a href="https://www.doi.org/10.5441/001/1.4v8q16qf" target="_blank">https://www.doi.org/10.5441/001/1.4v8q16qf</a> <br><br> Kays R, Davidson SC, Berger M, Bohrer G, Fiedler W, Flack A, Hirt J, Hahn C, Gauggel D, Russell B, et al. 2021. The Movebank system for studying global animal movement and demography. Methods Ecol Evol. https://doi.org/10.1111/2041-210X.13767 | https://doi.org/10.5441/001/1.4v8q16qf | CC_BY |
| Movebank | 92261778 | Cygnus cygnus | Dmitrijs Boiko | Wolfgang Fiedler | Dimitris Boiko, Wolfgang Fiedler & Martin Wikelski (2017): Project "Life Track Whooper Swan Latvia" at www.movebank.org | Movebank study 'LifeTrack Whooper Swan Latvia'. Accessed on [date]. | CC_BY |
| Movebank | 133992043 | Anser albifrons | Andrea Kölzsch | Andrea Kölzsch | Kölzsch A, Kruckenberg H, Glazov P, Müskens GJDM, Wikelski M. 2016. Data from: Towards a new understanding of migration timing: slower spring than autumn migration in geese reflects different decision rules for stopover use and departure. Movebank Data Repository. <a href="https://www.doi.org/10.5441/001/1.31c2v92f" target="_blank">https://www.doi.org/10.5441/001/1.31c2v92f</a> <br><br> Kölzsch A, Müskens GJDM, Kruckenberg H, Glazov P, Weinzierl R, Nolet BA, Wikelski M. 2016. Towards a new understanding of migration timing: slower spring than autumn migration in geese reflects different decision rules for stopover use and departure. Oikos. 125(10):1496–1507. https://doi.org/10.1111/oik.03121 | https://doi.org/10.5441/001/1.31c2v92f | CC_BY |

| Source | Study ID | Species | Principal Investigator | Contact person | Citation | Recommended reference | License type |
|---|---|---|---|---|---|---|---|
| Movebank | 146932094 | Grus grus | Mindaugas Dagys & Ramūnas Žydelis | Ramūnas Žydelis | Dagys M, Žydelis R. 2024. Data from: Study "Common Crane Lithuania GPS, 2015-2016". Movebank Data Repository. <a href="https://doi.org/10.5441/001/1.604" target="_blank">https://doi.org/10.5441/001/1.604</a> <br><br> These data are described in<br> Yanco SW, Oliver RY, Iannarilli F, Carlson BS, Heine G, Mueller U, Richter N, Vorneweg B, Andryushchenko Y, Batbayar N, et al. 2024. Migratory birds modulate niche tradeoffs in rhythm with seasons and life history. Proc Natl Acad Sci USA. https://doi.org/10.1073/pnas.2316827121 | https://doi.org/10.5441/001/1.604 | CC_BY |
| Movebank | 150597370 | Geronticus eremita | Didone Frigerio | Konrad Lorenz Research Station | Puehringer-Sturmayr V, Krejci J, Schuster R, et al. Space use and site fidelity in the endangered Northern Bald Ibis Geronticus eremita: Effects of age, season, and sex. Bird Conservation International. 2023;33:e10. doi:10.1017/S0959270922000053 | https://doi.org/10.1017/S0959270922000053 | CC_BY_NC |
| Movebank | 154820583 | Gyps fulvus | Goran Susic | Wolfgang Fiedler | Life Track Griffon Vulture Croatia Study | Movebank study 'LifeTrack Griffon Vulture Croatia'. Accessed on [date]. | CC_BY |
| Movebank | 172255794 | Steatornis caripensis | David A. Rodríguez. | Bernd Vorneweg | | Movebank study 'Lifetrack Oilbirds Costa Rica'. Accessed on [date]. | CC_0 |
| Movebank | 172972156 | Anthropoides virgo | Nyambayar Batbayar | Martin Wikelski | Batbayar N, Galtbalt B, Natsagdorj T, Sukhbaatar T, Wikelski M. 2024. Data from: Study "LifeTrack Mongolia Demoiselle cranes" Movebank Data Repository. <a href="https://doi.org/10.5441/001/1.599" target="_blank">https://doi.org/10.5441/001/1.599</a> <br><br> These data are described in<br> Yanco SW, Oliver RY, Iannarilli F, Carlson BS, Heine G, Mueller U, Richter N, Vorneweg B, Andryushchenko Y, Batbayar N, et al. 2024. Migratory birds modulate niche tradeoffs in rhythm with seasons and life history. Proc Natl Acad Sci USA. https://doi.org/10.1073/pnas.2316827121<br><br> Galtbalt B, Batbayar N, Sukhbaatar T, Vorneweg B, Heine G, Müller U, Wikelski M, Klaassen M. 2022. Differences in on-ground and aloft conditions explain seasonally different migration paths in demoiselle crane. Movement Ecology. https://doi.org/10.1186/s40462-022-00302-z <br><br> The processed data from Galtbalt et al. (2022) are published in Dryad at https://doi.org/10.5061/dryad.cnp5hqc1r | https://doi.org/10.5441/001/1.599 | CC_BY_NC |

| Source | Study ID | Species | Principal Investigator | Contact person | Citation | Recommended reference | License type |
|---|---|---|---|---|---|---|---|
| Movebank | 173641633 | Ciconia ciconia | Wolfgang Fiedler | Wolfgang Fiedler | Fiedler W, Niederer W, Schönenberger A, Flack A, Wikelski M. 2024. Data from: Study "LifeTrack White Stork Vorarlberg" (2016-2023). Movebank Data Repository. <a href="https://doi.org/10.5441/001/1.71r7pp6q_2" target="_blank">https://doi.org/10.5441/001/1.71r7pp6q_2</a> <br><br> These data are described in<br> Aikens EO, Nourani E, Fiedler W, Wikelski M, Flack A. 2024. Learning shapes the development of migratory behavior. Proc Natl Acad Sci USA. 121(12):e2306389121. https://doi.org/10.1073/pnas.2306389121<br><br>Brønnvik H, Nourani E, Fiedler W, Flack A. 2024. Experience reduces reliance on availability of conspecifics for route selection by a collectively migrating soaring bird. Curr Biol. 34(9):2030-2037.e3. https://doi.org/10.1016/j.cub.2024.03.052<br><br> Cheng Y, Fiedler W, Wikelski M, Flack A. 2019. "Closer-to-home" strategy benefits juvenile survival in a long-distance migratory bird. Ecology and Evolution. 9(16): 8945–8952. https://doi.org/10.1002/ece3.5395 | https://doi.org/10.5441/001/1.71r7pp6q_2 | CC_BY |
| Movebank | 175720577 | Ciconia ciconia | Wolfgang Fiedler | Wolfgang Fiedler | Maxhuni Q, Gashi A, Hoxha L, Wolf G, Fiedler W. 2021. Data from: Study "LifeTrack White Stork Kosova". Movebank Data Repository. <a href="https://www.doi.org/10.5441/001/1.s367rd3k" target="_blank">https://www.doi.org/10.5441/001/1.s367rd3k</a> <br><br> Kays R, Davidson SC, Berger M, Bohrer G, Fiedler W, Flack A, Hirt J, Hahn C, Gauggel D, Russell B, et al. 2021. The Movebank system for studying global animal movement and demography. Methods Ecol Evol. https://doi.org/10.1111/2041-210X.13767 | https://doi.org/10.5441/001/1.s367rd3k | CC_BY |
| Movebank | 178979729 | Canis lupus | Dave Latham, Stan Boutin | Maria Cecilia Latham | Latham ADM, Boutin S. 2019. Data from: Wolf ecology and caribou-primary prey-wolf spatial relationships in low productivity peatland complexes in northeastern Alberta. Movebank Data Repository. <a href="https://www.doi.org/10.5441/001/1.7vr1k987" target="_blank">https://www.doi.org/10.5441/001/1.7vr1k987</a> <br><br> Latham AD. 2009. Wolf ecology and caribou-primary prey-wolf spatial relationships in low productivity peatland complexes in northeastern Alberta [dissertation]. [Alberta (CA)]: ProQuest Dissertations Publishing, University of Alberta. NR55419. 197 p. http://search.proquest.com/docview/305051214 <br><br> Latham AD, Latham MC, Boyce MS, Boutin S. 2011. Movement responses by wolves to industrial linear features and their effect on woodland caribou in northeastern Alberta. Ecol Appl. 21(8):2854–2865. https://doi.org/10.1890/11-0666.1 | https://doi.org/10.5441/001/1.7vr1k987 | CC_BY_NC |

| Source | Study ID | Species | Principal Investigator | Contact person | Citation | Recommended reference | License type |
|---|---|---|---|---|---|---|---|
| Movebank | 180290122 | Thalassoica antarctica | Descamps | Descamps | Descamps S, Tarroux A, Cherel Y, Delord K, Godø OR, Kato A, Krafft BA, Lorentsen S, Ropert-Coudert Y, Skaret G, Varpe Ø. 2016. Data from: At-sea distribution and prey selection of Antarctic petrels and commercial krill fisheries. Movebank Data Repository. <a href="https://www.doi.org/10.5441/001/1.q4gn4q56" target="_blank">https://www.doi.org/10.5441/001/1.q4gn4q56</a><br><br>These data are described in<br>Descamps S, Tarroux A, Cherel Y, Delord K, Godø OR, Kato A, Krafft BA, Lorentsen S-H, Ropert-Coudert Y, Skaret G, et al. 2016. At-sea distribution and prey selection of Antarctic petrels and commercial krill fisheries. PLoS ONE. 11(8):e0156968. https://doi.org/10.1371/journal.pone.0156968 | https://doi.org/10.5441/001/1.q4gn4q56 | CC_BY |
| Movebank | 182459847 | Cygnus cygnus | Martin Wikelski | Wolfgang Fiedler | Boiko D, Wikelski M, Fiedler W. 2019. Data from: Moulting sites of Latvian whooper swan Cygnus cygnus cygnets fitted with GPS-GSM transmitters. Movebank Data Repository. <a href="https://www.doi.org/10.5441/001/1.f89984gn" target="_blank">https://www.doi.org/10.5441/001/1.f89984gn</a> <br><br>Boiko D, Wikelski M. 2019. Moulting sites of Latvian whooper swan Cygnus cygnus cygnets fitted with GPS-GSM transmitters. Wildfowl. 5: 229-241. https://wildfowl.wwt.org.uk/index.php/wildfowl/article/view/2714<br><br> Boiko D, Wikelski M. 2018. Abstract: Moulting sites of Latvian Whooper Swan Cygnus cygnus cygnets tagged with transmitters in summer 2016. p. 40: http://conference.emu.ee/userfiles/swan2018/Luigekonverents148X[illegible] | https://doi.org/10.5441/001/1.f89984gn | CC_BY |
| Movebank | 182746263 | Gyps himalayensis | Martin Wikelski | Sherub Sherub | Sherub S, Wikelski M, Fiedler W, Davidson SC. 2016. Data from: Behavioural adaptations to flight into thin air. Movebank Data Repository. <a href="https://www.doi.org/10.5441/001/1.143v2p2k" target="_blank">https://www.doi.org/10.5441/001/1.143v2p2k</a> <br><br> Sherub S, Bohrer G, Wikelski M, Weinzierl R. 2016. Behavioural adaptations to flight into thin air. Biology Letters. 12(10):20160432. https://doi.org/10.1098/rsbl.2016.0432 | https://doi.org/10.5441/001/1.143v2p2k | CC_0 |
| Movebank | 183209639 | Larus fuscus | Stefan Garthe | Stefan Garthe | Garthe S, Schwemmer P, Paiva VH, Corman A-M, Fock HO, Voigt CC, Adler S (2016) Terrestrial and marine foraging strategies of an opportunistic seabird species breeding in the Wadden Sea. PLoS ONE 11(8): e0159630. doi:10.1371/journal.pone.0159630 | https://doi.org/10.5441/001/1.nk286sc0 | CC_BY_NC |
| Movebank | 185780950 | Geronticus eremita | Didone Frigerio | Konrad Lorenz Research Station | Puehringer-Sturmayr V, Krejci J, Schuster R, et al. Space use and site fidelity in the endangered Northern Bald Ibis Geronticus eremita: Effects of age, season, and sex. Bird Conservation International. 2023;33:e10. doi:10.1017/S0959270922000053 | https://doi.org/10.1017/S0959270922000053 | CC_BY_NC |
| Movebank | 186178781 | Pernis apivorus,Gyps fulvus | Daniel Schmidt-Rothmund | Wolfgang Fiedler | | Movebank study 'Raptors NABU Moessingen public'. Accessed on [date]. | CC_BY |

| Source | Study ID | Species | Principal Investigator | Contact person | Citation | Recommended reference | License type |
|---|---|---|---|---|---|---|---|
| Movebank | 193545363 | Canis latrans,Puma concolor | Julie Young, PhD | Peter Mahoney | Mahoney PJ, Ebinger M, Jaeger M, Shivik JA, Young JK. 2017. Data from: Uncovering behavioural states from animal activity and site fidelity patterns. Movebank Data Repository. <a href="https://www.doi.org/10.5441/001/1.7d8301h2" target="_blank">https://www.doi.org/10.5441/001/1.7d8301h2</a> <br><br>These data are described in<br>Mahoney PJ, Young JK. 2017. Uncovering behavioural states from animal activity and site fidelity patterns. Methods Ecol Evol. 8(2):174-183. https://doi.org/10.1111/2041-210X.12658 | https://doi.org/10.5441/001/1.7d8301h2 | CC_0 |
| Movebank | 208413731 | Connochaetes taurinus | Randall Boone | Jared Stabach | Stabach JA, Hughey L, Reid RS, Worden JS, Leimgruber P, Boone RB. 2020. Data from: Study "White-bearded wildebeest in Kenya". Movebank Data Repository. <a href="https://www.doi.org/10.5441/001/1.h0t27719" target="_blank">https://www.doi.org/10.5441/001/1.h0t27719</a> <br><br> Stabach JA, Hughey LF, Crego RD, Fleming CH, Hopcraft JGC, Leimgruber P, Morrison TA, Ogutu JO, Reid RS, Worden JS, Boone RB. 2022. Increasing anthropogenic disturbance restricts wildebeest movement across East African grazing systems. Front Ecol Evol. https://doi.org/10.3389/fevo.2022.846171 <br><br> Walton T, Boone R. 2017. Analyzing wildebeest behavior and effects of humans on activity in southwest Kenya. Poster [stored in this study; see Files] <br><br> Stabach JA. 2015. Movement, resource selection, and the physiological stress response of white-bearded wildebeest [dissertation]. [Fort Collins (CO, USA)]: Colorado State University. https://doi.org/10217/167207 <br><br> Boone RB, Reid RS, Lilieholm RJ, Worden JS, Ogutu JO. 2009. Wildebeest forage acquisition in fragmented landscapes under variable climates. National Science Foundation (NSF) DEB Grant 0919383. | https://doi.org/10.5441/001/1.h0t27719 | CC_BY |
| Movebank | 209824313 | Canis lupus | Mark Hebblewhite | Mark Hebblewhite | Hebblewhite M, Merrill EH (2008) Modelling wildlife-human relationships for social species with mixed-effects resource selection models. Journal of Applied Ecology 45:834-844. doi:10.1111/j.1365-2664.2008.01466.x <br> <br> Hebblewhite M, Merrill EH (2007) Multiscale wolf predation risk for elk: does migration reduce risk? Oecologia 152:377-387. doi:10.1007/s00442-007-0661-y | https://doi.org/10.1111/j.1365-2664.2008.01466.x | CC_BY |

| Source | Study ID | Species | Principal Investigator | Contact person | Citation | Recommended reference | License type |
|---|---|---|---|---|---|---|---|
| Movebank | 212096177 | Ciconia ciconia | Wolfgang Fiedler | Wolfgang Fiedler | Fiedler W, Flack A, Schmid A, Reinhard U, Wikelski M. 2024. Data from: Study "LifeTrack White Stork Oberschwaben" (2014-2023). Movebank Data Repository. <a href="https://doi.org/10.5441/001/1.c42j3js7_2" target="_blank">https://doi.org/10.5441/001/1.c42j3js7_2</a> <br><br> These data are described in<br> Aikens EO, Nourani E, Fiedler W, Wikelski M, Flack A. 2024. Learning shapes the development of migratory behavior. Proc Natl Acad Sci USA. 121(12):e2306389121. https://doi.org/10.1073/pnas.2306389121<br><br> Brønnvik H, Nourani E, Fiedler W, Flack A. 2024. Experience reduces reliance on availability of conspecifics for route selection by a collectively migrating soaring bird. Curr Biol. 34(9):2030-2037.e3. https://doi.org/10.1016/j.cub.2024.03.052<br><br> Kays R, Davidson SC, Berger M, Bohrer G, Fiedler W, Flack A, Hirt J, Hahn C, Gauggel D, Russell B, et al. 2021. The Movebank system for studying global animal movement and demography. Methods Ecol Evol. https://doi.org/10.1111/2041-210X.13767<br><br> Cheng Y, Fiedler W, Wikelski M, Flack A. 2019. "Closer-to-home" strategy benefits juvenile survival in a long-distance migratory bird. Ecology and Evolution. 9(16): 8945–8952. https://doi.org/10.1002/ece3.5395 | https://doi.org/10.5441/001/1.c42j3js7_2 | CC_BY |
| Movebank | 216040785 | Rangifer tarandus | Dale R. Seip | Sarah Davidson | Seip DR, Price E. 2019. Data from: Science update for the South Peace Northern Caribou (Rangifer tarandus caribou pop. 15) in British Columbia. Movebank Data Repository. <a href="https://www.doi.org/10.5441/001/1.p5bn656k" target="_blank">https://www.doi.org/10.5441/001/1.p5bn656k</a> <br><br> Seip D, Jones E. 2017. Population status of Central Mountain caribou herds in British Columbia and response to recovery management actions. Ministry of Environment and Climate Change Strategy. 18 p. https://www2.gov.bc.ca/assets/gov/environment/plants-animals-and-ecosystems/wildlife-wildlife-habitat/regional-wildlife/northeast-region/caribou/central_mountains_caribou_population_report_2017.pdf <br><br> Johnson CJ, Ehlers LPW, Seip DR. 2015. Witnessing extinction: cumulative impacts across landscapes and the future loss of an evolutionarily significant unit of woodland caribou in Canada. Biol Conserv. 186:176-186. https://doi.org/10.1016/j.biocon.2015.03.012 <br><br> BC Ministry of Environment. 2014. Science update for the South Peace Northern caribou (Rangifer tarandus caribou pop. 15) in British Columbia. Victoria, BC. 43 p. https://www2.gov.bc.ca/assets/gov/environment/plants-animals-and-ecosystems/wildlife-wildlife-habitat/caribou/science_update_final_from_web_jan_2014.pdf <br><br> Jones ES, Gillingham MP, Seip DR, Heard DC. 2007. Comparison of seasonal habitat selection between threatened woodland caribou ecotypes in central British Columbia. Rangifer. 27(4):111-128. https://doi.org/10.7557/2.27.4.325 | https://doi.org/10.5441/001/1.p5bn656k | CC_BY |

| Source | Study ID | Species | Principal Investigator | Contact person | Citation | Recommended reference | License type |
|---|---|---|---|---|---|---|---|
| Movebank | 217784323 | Cathartes aura,Coragyps atratus | David Barber | David Barber | A more complete version of this dataset is stored in study "Vultures Acopian Center USA GPS".<br><br> Bildstein KL, Barber D, Bechard MJ, Graña Grilli M. 2016. Data from: Wing size but not wing shape is related to migratory behavior in a soaring bird. Movebank Data Repository. <a href="https://www.doi.org/10.5441/001/1.37r2b884" target="_blank">https://www.doi.org/10.5441/001/1.37r2b884</a> <br><br> Graña Grilli M, Lambertucci SA, Therrien J-F, Bildstein KL. 2017. Wing size but not wing shape is related to migratory behavior in a soaring bird. J Avian Biol. 48(5):669-678. https://doi.org/10.1111/jav.01220 <br><br> Dodge S, Bohrer G, Bildstein K, Davidson SC, Weinzierl R, Mechard MJ, Barber D, Kays R, Brandes D, Han J, et al. 2014. Environmental drivers of variability in the movement ecology of turkey vultures (Cathartes aura) in North and South America. Philos T Roy Soc B. 369(1643):20130195. https://doi.org/10.1098/rstb.2013.0195 | https://doi.org/10.5441/001/1.37r2b884 | CC_BY |
| Movebank | 220078181 | Thalassarche chrysostoma | David Thompson | Rachael Orben | Thompson DR, Torres LG, Sagar PM, Kroeger CE, Orben RA. 2017. Data from: Classification of animal movement behavior through residence in space and time. Movebank Data Repository. <a href="https://www.doi.org/10.5441/001/1.694p666h" target="_blank">https://www.doi.org/10.5441/001/1.694p666h</a><br><br> Torres LG, Orben RA, Tolkova I, Thompson DR. 2017. Classification of animal movement behavior through residence in space and time. PLoS ONE. 12(1):e0168513. https://doi.org/10.1371/journal.pone.0168513 | https://doi.org/10.5441/001/1.694p666h | CC_BY_NC |
| Movebank | 227843945 | Physeter macrocephalus | Daniel Palacios | Barb Lagerquist | Irvine L, Palacios DM, Urbán, Mate B. 2017. Sperm whale dive behavior characteristics derived from intermediate-duration archival tag data. Ecology and Evolution. 7:7822–7837. doi: <a href="https://doi.org/10.1002/ece3.3322">10.1002/ece3.3322</a> | https://doi.org/10.5441/001/1.rj857167 | CC_BY |
| Movebank | 233141114 | Grus vipio | Nyambayar Batbayar | Nyambayar Batbayar | Batbayar N, Galtbalt B, Natsagdorj T, Sukhbaatar T, Wikelski M. 2024. Data from: Study "White-naped crane Mongolia WSCC" Movebank Data Repository. <a href="https://doi.org/10.5441/001/1.600" target="_blank">https://doi.org/10.5441/001/1.600</a> <br><br> These data are described in<br> Yanco SW, Oliver RY, Iannarilli F, Carlson BS, Heine G, Mueller U, Richter N, Vorneweg B, Andryushchenko Y, Batbayar N, et al. 2024. Migratory birds modulate niche tradeoffs in rhythm with seasons and life history. Proc Natl Acad Sci USA. https://doi.org/10.1073/pnas.2316827121 | https://doi.org/10.5441/001/1.600 | CC_BY_NC |
| Movebank | 236953686 | Anas platyrhynchos | Wolfgang Fiedler | Wolfgang Fiedler | Duck data Max Planck Institute for Animal Behavior | Movebank study 'LifeTrack Ducks Lake Constance'. Accessed on [date]. | CC_0 |
| Movebank | 249737372 | Uria lomvia | Kyle Elliott | Allison Patterson | | Movebank study 'Thick-billed murre Gilchrist and Elliott Digges 2016'. Accessed on [date]. | CC_0 |

| Source | Study ID | Species | Principal Investigator | Contact person | Citation | Recommended reference | License type |
|---|---|---|---|---|---|---|---|
| Movebank | 268904527 | Wallabia bicolor | Manuela Fischer | Manuela Fischer | Fischer M, Di Stefano J, Gras P, Kramer-Schadt S, Sutherland DR, Coulson G, Stillfried M (2020) Data from: Circadian rhythms enable efficient resource selection in a human-modified landscape. Movebank Data Repository. <a href="https://www.doi.org/10.5441/001/1.6ss053tn" target="_blank">https://www.doi.org/10.5441/001/1.6ss053tn</a> <br><br> Fischer M, Di Stefano J, Gras P, Kramer-Schadt S, Sutherland DR, Coulson G, Stillfried M (2019) Circadian rhythms enable efficient resource selection in a human-modified landscape. Ecology and Evolution 9(13): 7509–7527. https://doi.org/10.1002/ece3.5283 | https://doi.org/10.5441/001/1.6ss053tn | CC_0 |
| Movebank | 270079388 | Uria lomvia | Grant Gilchrist | Allison Patterson | | Movebank study 'Thick-billed murre Gilchrist Cape Graham Moore 2016'. Accessed on [date]. | CC_0 |
| Movebank | 277815715 | Ichthyaetus audouinii | Faggio | G Faggio | Recorbet, B., Faggio, G. & Boutten, W. (2017. Etude par géolocalisation de l'utilisation de l'espace par le goéland d'Audouin reproducteur des ZPS du Golfe d'Ajaccio. CEN-Corse | Movebank study 'Ichthyaetus audouinii Corsica'. Accessed on [date]. | CC_BY_NC |
| Movebank | 295134472 | Equus quagga | Chamaille-Jammes | Chamaille-Jammes | | Movebank study 'Plains zebra Chamaill√©-Jammes Hwange NP'. Accessed on [date]. | CC_BY_NC |
| Movebank | 312267867 | Larus marinus,Larus fuscus,Larus argentatus | Grigori Tertitski | Grigori Tertitski | | Movebank study 'Solovki Larus Track'. Accessed on [date]. | CC_0 |
| Movebank | 326568799 | Gyps fulvus | Wolfgang Fiedler | Wolfgang Fiedler | Dieter Haas, Wolfgang Fiedler u.a. | Movebank study 'Griffon Vulture Albstadt Salzburg (Gypsi)'. Accessed on [date]. | CC_BY |
| Movebank | 358865092 | Uria lomvia | Kyle Elliott | Allison Patterson | | Movebank study 'Thick-billed murre Gilchrist and Elliott Digges 2015'. Accessed on [date]. | CC_0 |
| Movebank | 384868221 | Haliaeetus albicilla | Pawel; Mirski | Pawel; Mirski | Mirski P., unpubl. data. | Movebank study 'White-tailed Eagle Poland.'. Accessed on [date]. | CC_BY_NC |
| Movebank | 384882516 | Aquila pomarina | Pawel; Mirski | Pawel; Mirski | Mirski P., unpubl. | Movebank study 'Spotted eagles NE Poland'. Accessed on [date]. | CC_BY_NC |
| Movebank | 439735878 | Hypsignathus monstrosus | Sarah Olson | Sarah Olson | Olson SH, Bounga G, Ondzie A, Bushmaker T, Seifert SN, Kuisma E, et al. (2019) Lek-associated movement of a putative Ebolavirus reservoir, the hammer-headed fruit bat (Hypsignathus monstrosus), in northern Republic of Congo. PLoS ONE 14(10): e0223139. https://doi.org/10.1371/journal.pone.0223139 | https://doi.org/10.1371/journal.pone.0223139 | CC_BY |

| Source | Study ID | Species | Principal Investigator | Contact person | Citation | Recommended reference | License type |
|---|---|---|---|---|---|---|---|
| Movebank | 467107447 | Anser albifrons | Helmut Kruckenberg | Andrea Kölzsch | Kruckenberg H, Müskens GJDM, Ebbinge BS. 2018. Data from: A periodic Markov model to formalise animal migration on a network [white-fronted goose data]. Movebank Data Repository. <a href="https://www.doi.org/10.5441/001/1.kk38017f" target="_blank">https://www.doi.org/10.5441/001/1.kk38017f</a> <br><br> Kölzsch A, Kleyheeg E, Kruckenberg H, Kaatz M, Blasius B. 2018. A periodic Markov model to formalise animal migration on a network. R Soc Open Sci. 5(6):180438. https://doi.org/10.1098/rsos.180438 <br><br> Kölzsch A, Müskens GJDM, Kruckenberg H, Glazov P, Weinzierl R, Nolet BA, Wikelski M. 2016. Towards a new understanding of migration timing: slower spring than autumn migration in geese reflects different decision rules for stopover use and departure. Oikos. 125(10):1496-1507. https://doi.org/10.1111/oik.03121 <br><br> Van Wijk RE, Kölzsch A, Kruckenberg H, Ebbinge BS, Müskens GJDM, Nolet BA. 2012. Individually tracked geese follow peaks of temperature acceleration during spring migration. Oikos. 121(5):655-664. https://doi.org/10.1111/j.1600-0706.2011.20083.x | https://doi.org/10.5441/001/1.kk38017f | CC_BY |
| Movebank | 468460067 | Nasua narica,Pecari tajacu,Ateles geoffroyi,Cebus capucinus,Potos flavus | Margaret Crofoot | Rasmus Havmoeller | Kays R, Hirsch B, Caillaud D, Mares R, Alavi S, Havmøller RW, Crofoot M. 2023. Data from: Multi-scale movement syndromes for comparative analyses of animal movement patterns. Movebank Data Repository. <a href="https://doi.org/10.5441/001/1.295" target="_blank">https://doi.org/10.5441/001/1.295</a> <br><br> These data are described in<br> Kays R, Hirsch B, Caillaud D, Mares R, Alavi S, Havmøller RW, Crofoot M. 2023. Multi-scale movement syndromes for comparative analyses of animal movement patterns. Move Ecol. 11:61. https://doi.org/10.1186/s40462-022-00365-y | https://doi.org/10.5441/001/1.295 | CC_BY |
| Movebank | 492444603 | Canis lupus | Stan Boutin | Amanda Droghini | Bohm, H., A. Droghini, E. W. Neilson, and S. Boutin. 2018. Northern Alberta Grey Wolf, Study ID 492444603. Available on Movebank (movebank.org). Accessed on [access date]. | https://doi.org/10.1371/journal.pone.0205742 | CC_0 |
| Movebank | 496130871 | Anthropoides virgo | Elena Ilyashenko | Ivan Pokrovskiy | Ilyashenko EI, Pokrovsky I, Ilyashenko VY, Mudrik EA, Korepov M, Mnatsekanov RA, Politov D, Fiedler W, Wikelski M. 2024. Data from: Study "1000 Cranes. Russia. Taman. Azov.". Movebank Data Repository. <a href="https://doi.org/10.5441/001/1.594" target="_blank">https://doi.org/10.5441/001/1.594</a> <br><br> These data are described in<br> Yanco SW, Oliver RY, Iannarilli F, Carlson BS, Heine G, Mueller U, Richter N, Vorneweg B, Andryushchenko Y, Batbayar N, et al. 2024. Migratory birds modulate niche tradeoffs in rhythm with seasons and life history. Proc Natl Acad Sci USA. https://doi.org/10.1073/pnas.2316827121 | https://doi.org/10.5441/001/1.594 | CC_BY |

| Source | Study ID | Species | Principal Investigator | Contact person | Citation | Recommended reference | License type |
|---|---|---|---|---|---|---|---|
| Movebank | 496136836 | Anthropoides virgo | Elena Ilyashenko | Ivan Pokrovskiy | Ilyashenko EI, Pokrovsky I, Ilyashenko VY, Mudrik EA, Korepov M, Politov D, Gugueva E, Fiedler W, Wikelski M. 2024. Data from: Study "1000 Cranes. Russia. Volga-Ural Interfluve.". Movebank Data Repository. <a href="https://doi.org/10.5441/001/1.595" target="_blank">https://doi.org/10.5441/001/1.595</a> <br><br> These data are described in<br> Yanco SW, Oliver RY, Iannarilli F, Carlson BS, Heine G, Mueller U, Richter N, Vorneweg B, Andryushchenko Y, Batbayar N, et al. 2024. Migratory birds modulate niche tradeoffs in rhythm with seasons and life history. Proc Natl Acad Sci USA. https://doi.org/10.1073/pnas.2316827121 | https://doi.org/10.5441/001/1.595 | CC_BY |
| Movebank | 497087673 | Anthropoides virgo | Elena Ilyashenko | Ivan Pokrovskiy | Ilyashenko EI, Pokrovsky I, Mudrik EA, Politov D, Postelnykh K, Ilyashenko VY, Fiedler W, Wikelski M. 2024. Data from: Study "1000 Cranes. Russia. Altai.". Movebank Data Repository. <a href="https://doi.org/10.5441/001/1.596" target="_blank">https://doi.org/10.5441/001/1.596</a> <br><br> These data are described in<br> Yanco SW, Oliver RY, Iannarilli F, Carlson BS, Heine G, Mueller U, Richter N, Vorneweg B, Andryushchenko Y, Batbayar N, et al. 2024. Migratory birds modulate niche tradeoffs in rhythm with seasons and life history. Proc Natl Acad Sci USA. https://doi.org/10.1073/pnas.2316827121 | https://doi.org/10.5441/001/1.596 | CC_BY |
| Movebank | 497091077 | Grus vipio,Anthropoides virgo | Elena Ilyashenko | Ivan Pokrovskiy | Ilyashenko EI, Pokrovsky I, Goroshko OA, Mudrik EA, Politov D, Ilyashenko VY, Fiedler W, Wikelski M. 2024. Data from: Study "1000 Cranes. Russia. Transbaikalia.". Movebank Data Repository. <a href="https://doi.org/10.5441/001/1.592" target="_blank">https://doi.org/10.5441/001/1.592</a> <br><br> These data are described in<br> Yanco SW, Oliver RY, Iannarilli F, Carlson BS, Heine G, Mueller U, Richter N, Vorneweg B, Andryushchenko Y, Batbayar N, et al. 2024. Migratory birds modulate niche tradeoffs in rhythm with seasons and life history. Proc Natl Acad Sci USA. https://doi.org/10.1073/pnas.2316827121 | https://doi.org/10.5441/001/1.592 | CC_BY |
| Movebank | 497138661 | Anthropoides virgo | Elena Ilyashenko | Ivan Pokrovskiy | Ilyashenko EI, Pokrovsky I, Gavrilov AE, Zaripova S, Fiedler W, Wikelski M. 2024. Data from: Study "1000 Cranes. Southern Kazakhstan.". Movebank Data Repository. <a href="https://doi.org/10.5441/001/1.591" target="_blank">https://doi.org/10.5441/001/1.591</a> <br><br> These data are described in<br> Yanco SW, Oliver RY, Iannarilli F, Carlson BS, Heine G, Mueller U, Richter N, Vorneweg B, Andryushchenko Y, Batbayar N, et al. 2024. Migratory birds modulate niche tradeoffs in rhythm with seasons and life history. Proc Natl Acad Sci USA. https://doi.org/10.1073/pnas.2316827121 | https://doi.org/10.5441/001/1.591 | CC_BY |
| Movebank | 505156776 | Anser anser | Harald Grabenhofer | Harald Grabenhofer | Grabenhofer, H. et al, unpublished Data | Movebank study 'Graugans Zugverhalten Neusiedler See'. Accessed on [date]. | CC_BY_NC |

| Source | Study ID | Species | Principal Investigator | Contact person | Citation | Recommended reference | License type |
|---|---|---|---|---|---|---|---|
| Movebank | 542653863 | Grus grus | Elena Ilyashenko | Ivan Pokrovskiy | Ilyashenko EI, Pokrovsky I, Ilyashenko VY, Korepov M, Fiedler W, Wikelski M. 2024. Data from: Study "1000 Cranes. Russia. Common Crane.". Movebank Data Repository. <a href="https://doi.org/10.5441/001/1.593" target="_blank">https://doi.org/10.5441/001/1.593</a> <br><br> These data are described in<br> Yanco SW, Oliver RY, Iannarilli F, Carlson BS, Heine G, Mueller U, Richter N, Vorneweg B, Andryushchenko Y, Batbayar N, et al. 2024. Migratory birds modulate niche tradeoffs in rhythm with seasons and life history. Proc Natl Acad Sci USA. https://doi.org/10.1073/pnas.2316827121 | https://doi.org/10.5441/001/1.593 | CC_BY |
| Movebank | 560041066 | Ciconia ciconia | Ran Nathan | Shay Rotics | Rotics S, Kaatz M, Turjeman S, Zurell D, Wikelski M, Sapir N, Eggers U, Fiedler W, Jeltsch F, Nathan R. 2018. Data from: Early arrival at breeding grounds: causes, costs and a trade-off with overwintering latitude. Movebank Data Repository. <a href="https://www.doi.org/10.5441/001/1.v8d24552" target="_blank">https://www.doi.org/10.5441/001/1.v8d24552</a> <br><br> Rotics S, Kaatz M, Turjeman S, Zurell D, Wikelski M, Sapir N, Eggers U, Fiedler W, Jeltsch F, Nathan R. 2018. Early arrival at breeding grounds: causes, costs and a trade-off with overwintering latitude. J Anim Ecol. 87(6):1627-1638. https://doi.org/10.1111/1365-2656.12898 | https://doi.org/10.5441/001/1.v8d24552 | CC_BY |
| Movebank | 581719332 | Pelecanus occidentalis | Jordan Karubian | Jordan Karubian | | Movebank study 'Brown Pelican Raccoon Island LA 2014-2017'. Accessed on [date]. | CC_BY_NC |
| Movebank | 604806671 | Circus aeruginosus | Anny Anselin | Tanja Milotic | Anselin A, Desmet P, Milotic T, Janssens K, T'Jollyn F, De Bruyn L, Bouten W (2019) MH_WATERLAND - Western marsh harriers (Circus aeruginosus, Accipitridae) breeding near the Belgium-Netherlands border. Dataset. <a target="_blank" href="https://doi.org/10.5281/zenodo.3532940">https://doi.org/10.5281/zenodo.3532940</a> | https://doi.org/10.1007/s10336-020-01785-6 | CC_0 |
| Movebank | 631036041 | Anthropoides virgo | Nyambayar Batbayar | Nyambayar Batbayar | Batbayar N, Galtbalt B, Natsagdorj T, Sukhbaatar T, Wikelski M. 2024. Data from: Study "1000 Cranes. Mongolia." Movebank Data Repository. <a href="https://doi.org/10.5441/001/1.598" target="_blank">https://doi.org/10.5441/001/1.598</a> <br><br> These data are described in<br> Yanco SW, Oliver RY, Iannarilli F, Carlson BS, Heine G, Mueller U, Richter N, Vorneweg B, Andryushchenko Y, Batbayar N, et al. 2024. Migratory birds modulate niche tradeoffs in rhythm with seasons and life history. Proc Natl Acad Sci USA. https://doi.org/10.1073/pnas.2316827121 | https://doi.org/10.5441/001/1.598 | CC_BY_NC |

| Source | Study ID | Species | Principal Investigator | Contact person | Citation | Recommended reference | License type |
|---|---|---|---|---|---|---|---|
| Movebank | 657674643 | Anser albifrons | Andrea Kölzsch | Andrea Kölzsch | Kölzsch A, Müskens GJDM, Moonen S, Kruckenberg H, Glazov P, Wikelski M. 2019. Data from: Flyway connectivity and exchange primarily driven by moult migration in geese [North Sea population]. Movebank Data Repository. <a href="https://www.doi.org/10.5441/001/1.ct72m82n" target="_blank">https://www.doi.org/10.5441/001/1.ct72m82n</a> <br><br> Kölzsch A, Müskens GJDM, Szinai P, Moonen S, Glazov P, Kruckenberg H, Wikelski M, Nolet BA. 2019. Flyway connectivity and exchange primarily driven by moult migration in geese. Move Ecol. 7:3. https://doi.org/10.1186/s40462-019-0148-6 | https://doi.org/10.5441/001/1.ct72m82n | CC_BY |
| Movebank | 657965212 | Anser albifrons | Andrea Kölzsch | Andrea Kölzsch | Müskens GJDM, Szinai P, Sapi T, Kölzsch A, Wikelski M, Nolet BA. 2019. Data from: Flyway connectivity and exchange primarily driven by moult migration in geese [Pannonic population]. Movebank Data Repository. <a href="https://www.doi.org/10.5441/001/1.46b0mq21" target="_blank">https://www.doi.org/10.5441/001/1.46b0mq21</a> <br><br> Kölzsch A, Müskens GJDM, Szinai P, Moonen S, Glazov P, Kruckenberg H, Wikelski M, Nolet BA. 2019. Flyway connectivity and exchange primarily driven by moult migration in geese. Move Ecol. 7:3. https://doi.org/10.1186/s40462-019-0148-6 | https://doi.org/10.5441/001/1.46b0mq21 | CC_BY |
| Movebank | 672882373 | Milvus milvus | Patrick Scherler | Patrick Scherler |  | Movebank study 'Milvus_milvus_atlantismarcuard'. Accessed on [date]. | CC_BY_NC |
| Movebank | 673728219 | Ovis dalli | James P Lawler | James P Lawler | Lawler JP, Griffith B, Johnson D, Burch J. 2005. The effects of military jet overflights on Dall's Sheep in Interior Alaska. Report to the Department of the Air Force. National Park Service, Alaska Region Natural Resource Technical Report NPS/AR/NRTR-2005-51. 181 p. | https://doi.org/http://purl.access.gpo.gov/GPO/LPS64886 | CC_BY |
| Movebank | 681056756 | Pteropus lylei | Julien Cappelle | Julien Cappelle | Choden K, Ravon S, Epstein JH, Hoem T, Furey N, Gely M, Jolivot A, Hul V, Neung C, Tran A, Cappelle J (2020) Data from: Pteropus lylei primarily forages in residential areas in Kandal, Cambodia. Movebank Data Repository. <a href="https://www.doi.org/10.5441/001/1.j25661td" target="_blank">https://www.doi.org/10.5441/001/1.j25661td</a> <br><br> Choden K, Ravon S, Epstein JH, Hoem T, Furey N, Gely M, Jolivot A, Hul V, Neung C, Tran A, Cappelle J (2019) Pteropus lylei primarily forages in residential areas in Kandal, Cambodia. Ecology and Evolution 9(7): 4181–4191. https://doi.org/10.1002/ece3.5046 | https://doi.org/10.5441/001/1.j25661td | CC_0 |
| Movebank | 736029750 | Loxodonta africana | Abi Tamim Vanak | Abi Tamim Vanak | Slotow R, Thaker M, Vanak AT (2019) Data from: Fine-scale tracking of ambient temperature and movement reveals shuttling behavior of elephants to water. Movebank Data Repository. <a href="https://www.doi.org/10.5441/001/1.403h24q5" target="_blank">https://www.doi.org/10.5441/001/1.403h24q5</a> <br><br> Thaker M, Gupte PR, Prins HHT, Slotow R, Vanak AT. 2019. Fine-scale tracking of ambient temperature and movement reveals shuttling behavior of elephants to water. Front Ecol Evol. 7:4. https://doi.org/10.3389/fevo.2019.00004 | https://doi.org/10.5441/001/1.403h24q5 | CC_BY_NC |

| Source | Study ID | Species | Principal Investigator | Contact person | Citation | Recommended reference | License type |
|---|---|---|---|---|---|---|---|
| Movebank | 897981076 | Cervus elaphus | Mark Hebblewhite | Mark Hebblewhite | Data for 2001-2020 are published as<br>Hebblewhite M, Merrill EH, Martin H, Berg JE, Bohm H, Eggeman SL. 2020. Data from: Study "Ya Ha Tinda elk project, Banff National Park, 2001-2020 (females)". Movebank Data Repository. <a href="https://www.doi.org/10.5441/001/1.5g4h5t6c" target="_blank">https://www.doi.org/10.5441/001/1.5g4h5t6c</a> <br><br> Tucker MA, Böhning-Gaese K, Fagan WF, Fryxell JM, Van Moorter B, Alberts SC, Ali AH, Allen AM, Attias N, Avgar T, et al. 2018. Moving in the Anthropocene: global reductions in terrestrial mammalian movements. Science 359(6374):466–469. https://www.doi.org/10.1126/science.aam9712 <br><br> Eggeman S, Hebblewhite M, Bohm H, Whittington J, Merrill E. 2016. Behavioral flexibility in migratory behavior in a long-lived large herbivore. J Anim Ecol 85(3):785-797. https://www.doi.org/10.1111/1365-2656.12495 <br><br> Hebblewhite M, Merrill EH, Morgantini LE, White CA, Allen JR, Bruns E, Thurston L, Hurd TE. 2006. Is the migratory behaviour of montane elk herds in peril? The case of Alberta's Ya Ha Tinda elk herd. Wildlife Soc B 34(5):1280-1294. https://www.doi.org/10.2193/0091-7648(2006)34[1280:ITMBOM]2.0.CO;2 <br><br> Hebblewhite M, Merrill EH. 2009. Trade-offs between predation risk and forage differ between migrant strategies in a migratory ungulate. Ecology 90(12):3445-3454. https://www.doi.org/10.1890/08-2090.1 <br><br> Hebblewhite M, Merrill EH. 2008. Modelling wildlife-human relationships for social species with mixed-effects resource selection models. J Appl Ecol 45(3):834-844. https://www.doi.org/10.1111/j.1365-2664.2008.01466.x <br><br> Hebblewhite M, Merrill EH, McDermid G. 2008. A multi-scale test of the forage maturation hypothesis in a partially migratory ungulate population. Ecol Monogr 78(2):141-166. https://www.doi.org/10.1890/06-1708.1 <br><br> Hebblewhite M, Merrill EH. 2007. Multiscale wolf predation risk for elk: does migration reduce risk? Oecologia 152(2):377-387. https://www.doi.org/10.1007/s00442-007-0661-y <br><br> Hebblewhite M. 2006. Linking predation risk and forage to ungulate population dynamics [dissertation]. Edmonton, Alberta: University of Alberta. | https://doi.org/10.5441/001/1.5g4h5t6c | CC_0 |
| Movebank | 909521569 | Pernis apivorus | Patrik Byholm | Elham Nourani | Nourani E, Vansteelant WMG, Byholm P, Safi K. 2020 Dynamics of the energy seascape can explain intra-specific variations in sea-crossing behaviour of soaring birds. Biol. Lett. 16: 20190797. http://dx.doi.org/10.1098/rsbl.2019.0797 | https://doi.org/10.1098/rsbl.2019.0797 | CC_BY_NC |

| Source | Study ID | Species | Principal Investigator | Contact person | Citation | Recommended reference | License type |
|---|---|---|---|---|---|---|---|
| Movebank | 914913822 | Pelecanus onocrotalus | Ran Nathan | Ron Efrat | Efrat R, Hatzofe O, Nathan R. 2019. Data from: Landscape-dependent time versus energy optimisations in pelicans migrating through a large ecological barrier. Movebank Data Repository. <a href="https://www.doi.org/10.5441/001/1.hs79pk45" target="_blank">https://www.doi.org/10.5441/001/1.hs79pk45</a> <br><br> Efrat R, Hatzofe O, Nathan R. 2019. Landscape-dependent time versus energy optimisations in pelicans migrating through a large ecological barrier. Funct Ecol. 33(11):2161-2171. https://doi.org/10.1111/1365-2435.13426 | https://doi.org/10.5441/001/1.hs79pk45 | CC_0 |
| Movebank | 922263102 | Circus aeruginosus | Almut Schlaich & Tonio Schaub | Tanja Milotic | Koks B, Schlaich A, Schaub T, Klaassen R, Anselin A, Desmet P, Milotic T, Janssens K, Bouten W (2019) H_GRONINGEN - Western marsh harriers (Circus aeruginosus, Accipitridae) breeding in Groningen (the Netherlands). Dataset. <a target="_blank" href="https://doi.org/10.5281/zenodo.3552507">https://doi.org/10.5281/zenodo.3552507</a> | https://doi.org/10.3897/zookeys.947.52570 | CC_0 |
| Movebank | 924418031 | Gallinago gallinago,Larus canus | Marcin | Marcin | www.interrex-tracking.com | Movebank study 'Common Snipe GPS-GSM logger Poland-Russia'. Accessed on [date]. | CC_0 |

| Source | Study ID | Species | Principal Investigator | Contact person | Citation | Recommended reference | License type |
|---|---|---|---|---|---|---|---|
| Movebank | 933711994 | Cervus elaphus | Mark S. Boyce | Sarah Davidson | Boyce MS and Ciuti S (2020) Data from: Human selection of elk behavioural traits in a landscape of fear. Movebank Data Repository. <a href="https://www.doi.org/10.5441/001/1.j484vk24" target="_blank">https://www.doi.org/10.5441/001/1.j484vk24</a> <br><br> Paton DG, Ciuti S, Quinn M, Boyce MS (2017) Hunting exacerbates the response to human disturbance in large herbivores while migrating through a road network. Ecosphere 8(6): e01841. https://doi.org/10.1002/ecs2.1841 <br><br> Prokopenko CM, Boyce MS, Avgar T (2017) Characterizing wildlife behavioural responses to roads using integrated step selection analysis. Journal of Applied Ecology 54: 470-479. https://doi.org/10.1111/1365-2664.12768 <br><br> Prokopenko CM, Boyce MS, Avgar T (2017) Extent-dependent habitat selection in a migratory large herbivore: road avoidance across scales. Landscape Ecology 32: 313-325. https://doi.org/10.1007/s10980-016-0451-1 <br><br> Roberts DR, Bahn V, Ciuti S, Boyce MS, Elith J, Guillera-Arroita G, Hauenstein S, Lahoz-Monfort JJ, Schöder B, Thuiller W, Warton DI, Wintle BA, Hartig F, Dormann CF (2017) Cross-validation strategies for data with temporal, spatial, hierarchical, or phylogenetic structure. Ecography 40: 913-929. https://doi.org/10.1111/ecog.02881 <br><br> Thurfjell H, Ciuti S, Boyce MS (2017) Learning from the mistakes of others: How female elk (Cervus elaphus) adjust behaviour with age to avoid hunters. PLoS ONE 12(6): e0178082. https://doi.org/10.1371/journal.pone.0178082 <br><br> Benz RA, Boyce MS, Thurfjell H, Paton DG, Musiani M, Dormann CF, et al. (2016) Dispersal ecology informs design of large-scale wildlife corridors. PLoS ONE 11(9): e0162989. https://doi.org/10.1371/journal.pone.0162989 <br><br> Ensing EP, Ciuti S, de Wijs FALM, Lentferink DH, ten Hoedt A, Boyce MS, Hut RA (2014) GPS based daily activity patterns in European red deer and North American elk (Cervus elaphus): indication for a weak circadian clock in ungulates. PLoS ONE 9(9): e106997. https://doi.org/10.1371/journal.pone.0106997 <br><br> Killeen J, Thurfjell H, Ciuti S, Paton D, Musiani M, Boyce MS (2014) Habitat selection during ungulate dispersal and exploratory movement at broad and fine scale with implications for conservation management. Movement Ecology 2:15. https://doi.org/10.1186/s40462-014-0015-4 <br><br> Thurfjell H, Ciuti S, Boyce MS (2014) Applications of step-selection functions in ecology and conservation. Movement Ecology 2:4. https://doi.org/10.1186/2051-3933-2-4 <br><br> Ciuti S, Muhly TB, Paton DG, McDevitt AD, Musiani M, Boyce MS (2012) Human selection of elk behavioural traits in a landscape of fear. Proceedings of the Royal Society B 279(1746): 4407-4416 | https://doi.org/10.5441/001/1.j484vk24 | CC_BY |
| Movebank | 938783961 | Buteo buteo,Circus aeruginosus | Geert Spanoghe | Tanja Milotic | Spanoghe G, Desmet P, Milotic T, Janssens K, De Regge N, Vanoverbeke J, Bouten W (2019) MH_ANTWERPEN - Western marsh harriers (Circus aeruginosus, Accipitridae) breeding near Antwerp (Belgium). Dataset. <a target="_blank" href="https://doi.org/10.5281/zenodo.3550093">https://doi.org/10.5281/zenodo.3550093</a> | https://doi.org/10.3897/zookeys.947.52570 | CC_0 |

| Source | Study ID | Species | Principal Investigator | Contact person | Citation | Recommended reference | License type |
|---|---|---|---|---|---|---|---|
| Movebank | 943824007 | Balaenoptera physalus,Balaenoptera musculus | Daniel Palacios | Barb Lagerquist | Irvine LM, Palacios DM, Lagerquist BA, Mate BR, Follett TM. 2019. Data from: Scales of blue and fin whale feeding behavior off California, USA, with implications for prey patchiness. Movebank Data Repository. <a href="https://www.doi.org/10.5441/001/1.47h576f2" target="_blank">https://www.doi.org/10.5441/001/1.47h576f2</a> <br><br> Irvine LM, Palacios DM, Lagerquist BA, Mate BR. 2019. Scales of blue and fin whale feeding behavior off California, USA, with implications for prey patchiness. Front Ecol Evol. 7:338. https://doi.org/10.3389/fevo.2019.00338 <br><br> Irvine LM, Winsor MH, Follett TM, Mate BR, Palacios DM. 2020. An at-sea assessment of Argos location accuracy for three species of large whales, and the effect of deep-diving behavior on location error. Animal Biotelemetry. 8:20. https://doi.org/10.1186/s40317-020- [illegible] | https://doi.org/10.5441/001/1.47h576f2 | CC_BY |
| Movebank | 985143423 | Larus fuscus | Eric Stienen | Peter Desmet | Stienen EWM, Desmet P, Milotic T, Hernandez F, Deneudt K, Bouten W, Müller W, Matheve H, Lens L (2019) LBBG_ZEEBRUGGE - Lesser black-backed gulls (Larus fuscus, Laridae) breeding at the southern North Sea coast (Belgium and the Netherlands). Dataset. <a target="_blank" href="https://doi.org/10.5281/zenodo.3540799">https://doi.org/10.5281/zenodo.3540799</a> | https://doi.org/10.5281/zenodo.3540799 | CC_0 |
| Movebank | 986040562 | Larus argentatus | Eric Stienen | Peter Desmet | Stienen EWM, Desmet P, Milotic T, Hernandez F, Deneudt K, Matheve H, Bouten W (2019) HG_OOSTENDE - Herring gulls (Larus argentatus, Laridae) breeding at the southern North Sea coast (Belgium). Dataset. <a target="_blank" href="https://doi.org/10.5281/zenodo.3541811">https://doi.org/10.5281/zenodo.3541811</a> | https://doi.org/10.5281/zenodo.3541811 | CC_0 |
| Movebank | 1007794443 | Larus marinus | Stefan Garthe | Stefan Garthe | Borrmann RM, Phillips RA, Clay TA, Garthe S. 2019. Data from: High foraging site fidelity and spatial segregation among individual great black-backed gulls. Movebank Data Repository. <a href="https://www.doi.org/10.5441/001/1.ht5jf68h" target="_blank">https://www.doi.org/10.5441/001/1.ht5jf68h</a> <br><br> Borrmann RM, Phillips RA, Clay TA, Garthe S. 2019. High foraging site fidelity and spatial segregation among individual great black-backed gulls. J Avian Biol. 50(12). https://doi.org/10.1111/jav.02156 | https://doi.org/10.5441/001/1.ht5jf68h | CC_BY_NC |
| Movebank | 1030734949 | Cygnus columbianus | Rascha Nuijten | Rascha Nuijten | Nuijten RJM, Gerrits T, de Vries PP, Müskens GJDM, Nolet BA. 2020. Data from: Less is more: on-board lossy compression of accelerometer data increases biologging capacity. Movebank Data Repository. <a href="https://www.doi.org/10.5441/001/1.8ms7mm80" target="_blank">https://www.doi.org/10.5441/001/1.8ms7mm80</a> <br><br> Nuijten RJM, Gerrits T, Shamoun-Baranes J, Nolet BA. 2020. Less is more: on-board lossy compression of accelerometer data increases biologging capacity. J Anim Ecol. 89(1):237-247. https://www.doi.org/10.1111/1365-2656.13164 | https://doi.org/10.5441/001/1.8ms7mm80 | CC_0 |

| Source | Study ID | Species | Principal Investigator | Contact person | Citation | Recommended reference | License type |
|---|---|---|---|---|---|---|---|
| Movebank | 1049685237 | Anser albifrons | Andrea Kölzsch | Andrea Kölzsch | Kölzsch A, Müskens GJDM, Glazov P, Kruckenberg H, Wikelski M. 2020. Data from: Goose parents lead migration V. Movebank Data Repository. <a href="https://www.doi.org/10.5441/001/1.ms87s2m6" target="_blank">https://www.doi.org/10.5441/001/1.ms87s2m6</a><br><br>Kölzsch A, Flack A, Müskens GJDM, Kruckenberg H, Glazov P, Wikelski M. 2020. Goose parents lead migration V. J Avian Biol. 51(3). https://doi.org/10.1111/jav.02392 | https://doi.org/10.5441/001/1.ms87s2m6 | CC_0 |
| Movebank | 1071101052 | Larus argentatus | Robert Ronconi | Robert Ronconi | Ronconi RA, Shlepr KR. 2020. Data from: Study "Herring Gulls (Larus Argentatus); Ronconi; Kent Island, Canada". Movebank Data Repository. <a href="https://www.doi.org/10.5441/001/1.t5n4s456" target="_blank">https://www.doi.org/10.5441/001/1.t5n4s456</a> <br><br> Anderson CM, Gilchrist HG, Ronconi RA, Shlepr KR, Clark DE, Fifield DA, Robertson GJ, Mallory ML. 2020. Both short and long distance migrants use energy-minimizing strategies in North American herring gulls. Movement Ecology. 8:26. https://doi.org/10.1186/s40462-020-00207-9 | https://doi.org/10.5441/001/1.t5n4s456 | CC_0 |
| Movebank | 1071134107 | Larus argentatus | Robert Ronconi | Robert Ronconi | Ronconi RA, Shlepr KR. 2020. Data from: Study "Herring Gulls (Larus Argentatus); Ronconi; Brier Island, Canada". Movebank Data Repository. <a href="https://www.doi.org/10.5441/001/1.282vr7kd" target="_blank">https://www.doi.org/10.5441/001/1.282vr7kd</a> <br><br> Anderson CM, Gilchrist HG, Ronconi RA, Shlepr KR, Clark DE, Fifield DA, Robertson GJ, Mallory ML. 2020. Both short and long distance migrants use energy-minimizing strategies in North American herring gulls. Movement Ecology. 8:26. https://doi.org/10.1186/s40462-020-00207-9 | https://doi.org/10.5441/001/1.282vr7kd | CC_0 |
| Movebank | 1073231887 | Buteo jamaicensis | Pete Bloom | Pete Bloom | Bloom PH (2015) Northward summer migration of red-tailed hawks fledged from southern latitudes. Journal of Raptor Research 49(1): 1-17. https://doi.org/10.3356/jrr-14-54.1 <br><br> Bloom PH (2011) Vagrancy, natal dispersal and migrations of red-tailed and red-shouldered hawks banded in the Pacific Flyway. Ph.D. dissertation, University of Idaho, Moscow, ID USA. | https://doi.org/10.3356/jrr-14-54.1 | CC_BY |
| Movebank | 1077432441 | Pteropus poliocephalus | Wayne Boardman | David Roshier | Roshier D, Boardman WSJ. 2021. Data from: Spring foraging movements of an urban population of grey-headed flying foxes (Pteropus poliocephalus). Movebank Data Repository. <a href="https://www.doi.org/10.5441/001/1.5bd6pq55" target="_blank">https://www.doi.org/10.5441/001/1.5bd6pq55</a> <br><br> Boardman WSJ, Roshier D, Reardon T, Burbidge K, McKeown A, Westcott DA, Caraguel CGB, Prowse TAA. 2021. Spring foraging movements of an urban population of grey-headed flying foxes (Pteropus poliocephalus). J Urban Ecol. 7(1):juaa034. https://doi.org/10.1093/jue/juaa034 | https://doi.org/10.5441/001/1.5bd6pq55 | CC_0 |

| Source | Study ID | Species | Principal Investigator | Contact person | Citation | Recommended reference | License type |
|---|---|---|---|---|---|---|---|
| Movebank | 1080341217 | Larus argentatus | Dan Clark | Christine Anderson | Clark DE, Mackenzie SA, Koenen K, Whitney J, DeStefano S. 2020. Data from: Study "Herring Gulls (Larus Argentatus); Clark; Massachussets, United States". Movebank Data Repository. <a href="https://www.doi.org/10.5441/001/1.3th8b5q3" target="_blank">https://www.doi.org/10.5441/001/1.3th8b5q3</a> <br><br> Anderson CM, Gilchrist HG, Ronconi RA, Shlepr KR, Clark DE, Fifield DA, Robertson GJ, Mallory ML. 2020. Both short and long distance migrants use energy-minimizing strategies in North American herring gulls. Movement Ecology. 8:26. https://doi.org/10.1186/s40462-020-00207-9 | https://doi.org/10.5441/001/1.3th8b5q3 | CC_0 |
| Movebank | 1080341737 | Larus argentatus | Robert Ronconi | Robert Ronconi | Ronconi RA, Taylor PD. 2020. Data from: Study "Herring Gulls (Larus Argentatus); Ronconi; Sable Island, Canada". Movebank Data Repository. <a href="https://www.doi.org/10.5441/001/1.3264ss3v" target="_blank">https://www.doi.org/10.5441/001/1.3264ss3v</a> <br><br> Anderson CM, Gilchrist HG, Ronconi RA, Shlepr KR, Clark DE, Fifield DA, Robertson GJ, Mallory ML. 2020. Both short and long distance migrants use energy-minimizing strategies in North American herring gulls. Movement Ecology. 8:26. https://doi.org/10.1186/s40462-020-00207-9 | https://doi.org/10.5441/001/1.3264ss3v | CC_0 |
| Movebank | 1087068449 | Tockus deckeni | Martin Wikelski | Katherine Mertes | Mertes K, Jetz W, Wikelski M (2020) Data from: Hierarchical multi-grain models improve descriptions of species’ environmental associations, distribution, and abundance. Movebank Data Repository. <a href="https://www.doi.org/10.5441/001/1.cp97k9j1" target="_blank">https://www.doi.org/10.5441/001/1.cp97k9j1</a> <br><br> Schwartz K, Jarzyna M, Jetz W (2020) Hierarchical multi-grain models improve descriptions of species’ environmental associations, distribution, and abundance. Ecological Applications. https://doi.org/10.1002/eap.2117 | https://doi.org/10.5441/001/1.cp97k9j1 | CC_0 |
| Movebank | 1088836380 | Lynx rufus,Pekania pennanti | Scott LaPoint | Scott LaPoint | LaPoint et al. in progress | Movebank study 'Carnivore movements near Black Rock Forest New York'. Accessed on [date]. | CC_BY |
| Movebank | 1091848505 | Larus ,Larus hyperboreus,Larus glaucescens,Larus smithsonianus | Andrew Ramey | Andrew Ramey | </br>Ramey, A.M., Ahlstrom, C.A., van Toor, M.L., Woksepp, H., Chandler, J.C., Reed, J.A., Reeves, A.B., Waldenström, J., Franklin, A.B., Bonnedahl, J., and Douglas, D.C. 2020, Tracking data for three large-bodied gull species and hybrids (Larus spp.) (ver 1.0, June 2020): U.S. Geological Survey data release, <a href="https://doi.org/10.5066/P9FZ4OJW" target="_blank">doi:10.5066/P9FZ4OJW</a></br> | https://doi.org/10.1016/j.scitotenv.2020.144551 | CC_0 |
| Movebank | 1099562810 | Haematopus ostralegus | Geert Spanoghe | Tanja Milotic | Spanoghe G, Desmet P, Milotic T, Van Ryckegem G, Vanoverbeke J, Ens BJ, Bouten W (2020) O_WESTERSCHELDE - Eurasian oystercatchers (Haematopus ostralegus, Haematopodidae) breeding in East Flanders (Belgium). Dataset. <a target="_blank" href="https://doi.org/10.5281/zenodo.3734898">https://doi.org/10.5281/zenodo.3734898</a> | https://doi.org/10.3897/zookeys.1123.90623 | CC_0 |

| Source | Study ID | Species | Principal Investigator | Contact person | Citation | Recommended reference | License type |
|---|---|---|---|---|---|---|---|
| Movebank | 1120749252 | Nasua narica,Pecari tajacu,Ateles geoffroyi,Cebus capucinus,Potos flavus | Meg Crofoot | Meg Crofoot | Kays R, Hirsch B, Caillaud D, Mares R, Alavi S, Havmøller RW, Crofoot M. 2023. Data from: Multi-scale movement syndromes for comparative analyses of animal movement patterns. Movebank Data Repository. <a href="https://doi.org/10.5441/001/1.295" target="_blank">https://doi.org/10.5441/001/1.295</a> <br><br> These data are described in<br> Kays R, Hirsch B, Caillaud D, Mares R, Alavi S, Havmøller RW, Crofoot M. 2023. Multi-scale movement syndromes for comparative analyses of animal movement patterns. Move Ecol. 11:61. https://doi.org/10.1186/s40462-022-00365-y | https://doi.org/10.5441/001/1.295 | CC_BY |
| Movebank | 1123149708 | Pagophila eburnea | Morten Frederiksen | Morten Frederiksen | Frederiksen, M. (2019) GPS tracking of breeding ivory gulls in NE Greenland 2018-2019 | Movebank study 'Ivory gull N Greenland 2018/19'. Accessed on [date]. | CC_BY_NC |
| Movebank | 1207235729 | Canis latrans | Nucharin Songsasen, PhD, DVM | Jared Stabach | | Movebank study 'Coyote (Canis latrans) movement in the northern blue ridge mountains - Virginia'. Accessed on [date]. | CC_0 |
| Movebank | 1208105916 | Larus cachinnans | INTERREX | INTERREX | | Movebank study 'Caspian Gulls - Poland'. Accessed on [date]. | CC_0 |
| Movebank | 1229945587 | Grus grus | Petras Kurlavičius | Aleksej Perzhu | Unpublished & ongoing study by Lithuanian University of Educational Sciences (LEU) | Movebank study 'Common Crane 2020 (Lithuanian University of Educational Studies; LEU)'. Accessed on [date]. | CC_0 |
| Movebank | 1241071371 | Vulpes lagopus | Dominique Berteaux | Dominique Berteaux | Clermont J., Grenier-Potvin A., Duchesne É., Couchoux C., Dulude-de Broin F., Beardsell A., Bêty J., Berteaux D. 2021. The predator activity landscape predicts the anti-predator behavior and distribution of prey in a tundra community. Ecosphere 12(12):e03858. <a href="https://doi.org/10.1002/ecs2.3858"target="_blank">https://doi.org/10.1002/ecs2.3858</a><br><br>Clermont J., Woodward-Gagné S., Berteaux D. 2021. Digging into the behaviour of an active hunting predator: arctic fox prey caching events revealed by accelerometry. Mov Ecol 9:58. <a href="https://doi.org/10.1186/s40462-021-00295-1" target="_blank">https://doi.org/10.1186/s40462-021-00295-1</a><br><br>Poulin M-P, Clermont J, Berteaux D. 2021. Extensive daily movement rates measured in territorial arctic foxes. Ecol Evol. 11(6):2503-14. <a href="https://www.doi.org/10.1002/ece3.7165 " target="_blank">https://www.doi.org/10.1002/ece3.7165</a><br><br>Grenier-Potvin A, Clermont J, Gauthier G, Berteaux D. 2021. Prey and habitat distribution are not enough to explain predator habitat selection: addressing intraspecific interactions, behavioural state and time. Mov Ecol. 9:12. <a href="https://www.doi.org/10.1186/s40462-021-00250-0" target="_blank">https://www.doi.org/10.1186/s40462-021-00250-0</a><br><br> | https://doi.org/10.1002/ece3.7165 | CC_0 |
| Movebank | 1252230318 | Ciconia ciconia | Marcin Tobółka | Marcin Tobółka | Tobółka, M., Nowak, B., Białas, J.T., Jagiełło, Z., Zielińska, Z. 2020. The influence of parental experience and conditions during breeding and non-breeding period on reproductive success of long-lived birds, an example of the White Stork. | Movebank study 'White stork Ciconidae Western Poland'. Accessed on [date]. | CC_0 |

| Source | Study ID | Species | Principal Investigator | Contact person | Citation | Recommended reference | License type |
|---|---|---|---|---|---|---|---|
| Movebank | 1258895879 | Larus fuscus,Larus argentatus | Eric Stienen | Peter Desmet | Stienen EWM, Buijs R-J, de Visser J, Fijn R, Lilipaly S, Platteeuw M, Desmet P (2023) DELTATRACK - Herring gulls (Larus argentatus, Laridae) and lesser black-backed gulls (Larus fuscus, Laridae) breeding at Neeltje Jans (Netherlands). Dataset. <a target="_blank" href="https://doi.org/10.5281/zenodo.10209520">https://doi.org/10.5281/zenodo.10209520</a> | https://doi.org/10.5281/zenodo.10209520 | CC_0 |
| Movebank | 1259686571 | Larus fuscus,Larus argentatus | Eric Stienen | Peter Desmet | Stienen EWM, Müller W, Lens L, Milotic T, Desmet P (2021) LBBG_JUVENILE - Juvenile lesser black-backed gulls (Larus fuscus, Laridae) and herring gulls (Larus argentatus, Laridae) hatched in Zeebrugge (Belgium). Dataset. <a target="_blank" href="https://doi.org/10.5281/zenodo.5075868">https://doi.org/10.5281/zenodo.5075868</a> | https://doi.org/10.5281/zenodo.5075868 | CC_0 |
| Movebank | 1266784970 | Corvus corone | JIGUET Frédéric | Jiguet Frédéric | | Movebank study 'Corvus corone [ID_PROG 883]'. Accessed on [date]. | CC_BY_NC |
| Movebank | 1278021460 | Buteo buteo,Circus cyaneus,Circus pygargus,Asio flammeus,Circus aeruginosus | Geert Spanoghe | Peter Desmet | Spanoghe G, Janssens K, Klaassen R, Schaub T, Milotic T, Desmet P (2021) BOP_RODENT - Rodent specialized birds of prey (Circus, Asio, Buteo) in Flanders (Belgium). Dataset. <a target="_blank" href="https://doi.org/10.5281/zenodo.5735405">https://doi.org/10.5281/zenodo.5735405</a> | https://doi.org/10.5281/zenodo.5735405 | CC_0 |
| Movebank | 1326534946 | Rissa ,Rissa tridactyla | Kyle Elliott | Kyle Elliott | | Movebank study 'Black-legged Kittiwake Rissa tridactyla Middleton Island'. Accessed on [date]. | CC_BY |
| Movebank | 1375204238 | Pygoscelis adeliae | Yan Ropert-Coudert | Candice Michelot | Michelot et al (Accepted in Plos One) Adélie penguins foraging consistency and site fidelity are conditioned by breeding status and environmental conditions | https://doi.org/10.1371/journal.pone.0244298 | CC_BY_NC |
| Movebank | 1393954358 | Cathartes aura | Martin Wikelski | Kamran Safi | | Movebank study 'Cathartes aura MPIAB Cuba'. Accessed on [date]. | CC_BY_NC |
| Movebank | 1395952585 | Numenius arquata | Philipp Schwemmer | Philipp Schwemmer | Schwemmer P, Garthe S. 2021. Data from: Migrating curlews on schedule: departure and arrival patterns of a long-distance migrant depend on time and breeding location rather than on wind conditions. Movebank Data Repository. <a href="https://www.doi.org/10.5441/001/1.715k46g2" target="_blank">https://www.doi.org/10.5441/001/1.715k46g2</a> <br><br> Schwemmer P, Mercker M, Vanselow KH, Bocher P, Garthe S. 2021. Migrating curlews on schedule: departure and arrival patterns of a long-distance migrant depend on time and breeding location rather than on wind conditions. Movement Ecology. https://doi.org/10.1186/s40462-021-00252-y | https://doi.org/10.5441/001/1.715k46g2 | CC_0 |
| Movebank | 1404033416 | Anas rubripes | Chris Williams | Amanda Hoyt | | Movebank study 'NC Breeding ABDU'. Accessed on [date]. | CC_0 |
| Movebank | 1406291704 | Panthera onca | Ronaldo Morato | Ronaldo Morato | Morato, RG; Tortato, F. | Movebank study 'Jaguar Rescue'. Accessed on [date]. | CC_BY |
| Movebank | 1410035327 | Ciconia ciconia | Ben Carlson | Ben Carlson | Carlson BS, Rotics S, Nathan R, Wikelski M, Jetz W. 2021. Individual environmental niches in mobile organisms. Nat Commun. 12:4572. doi: <a href="10.1038/s41467-021-24826-x">10.1038/s41467-021-24826-x</a> | https://doi.org/10.5441/001/1.rj21g1p1 | CC_0 |

| Source | Study ID | Species | Principal Investigator | Contact person | Citation | Recommended reference | License type |
|---|---|---|---|---|---|---|---|
| Movebank | 1415844328 | Anser fabalis | Antti Piironen | Antti Piironen | Piironen A, Paasivaara A, Laaksonen T. 2021. Data from: Birds of three worlds: moult migration to high Arctic expands a boreal-temperate flyway to a third biome. Movebank Data Repository. <a href="https://www.doi.org/10.5441/001/1.22kk5126" target="_blank">https://www.doi.org/10.5441/001/1.22kk5126</a> <br><br> Piironen A, Paasivaara A, Laaksonen T. 2021. Birds of three worlds: moult migration to high Arctic expands a boreal-temperate flyway to a third biome. Movement Ecology. 9:47. https://www.doi.org/10.1186/s40462-021-00284-4 | https://doi.org/10.5441/001/1.22kk5126 | CC_BY |
| Movebank | 1426277950 | Tyto furcata | Matthew Johnson | Allison Huysman | Huysman AE, Castañeda XA, Johnson MD. 2021. Data from: Habitat selection by a predator of rodent pests is resilient to wildfire in a vineyard agroecosystem. Movebank Data Repository. <a href="https://www.doi.org/10.5441/001/1.82b5h1rk" target="_blank">https://www.doi.org/10.5441/001/1.82b5h1rk</a> <br><br> Huysman AE, Johnson MD. 2021. Habitat selection by a predator of rodent pests is resilient to wildfire in a vineyard agroecosystem. Ecol Evol. https://doi.org/10.1002/ece3.8416 <br><br> Castañeda XA, Huysman AE, Johnson MD. 2021. Barn Owls select uncultivated habitats for hunting in a winegrape growing region of California. Ornithol Appl. 123(1):duaa058. https://doi.org/10.1093/ornithapp/duaa058 | https://doi.org/10.5441/001/1.82b5h1rk | CC_BY |
| Movebank | 1429664821 | Connochaetes taurinus | Tom Morrison | Tom Morrison | Morrison TA, Link W, Newmark W, Foley C, Bolger DT. 2016. Tarangire revisited: population consequences of loss of migratory connectivity in a tropical ungulate. Biological Conservation. 197, 53-60. | https://doi.org/10.1016/j.biocon.2016.02.034 | CC_BY_NC |
| Movebank | 1445058753 | Larus fuscus | Marcin | Marcin | www.interrex-tracking.com | Movebank study 'Larus heuglini_1'. Accessed on [date]. | CC_0 |
| Movebank | 1448377103 | Mycteria americana | Mathieu Basille | Mathieu Basille | Basille M, Borkhataria RR, Bryan AL, Bucklin DN, Picardi S, Frederick PC. 2021. Data from: Study "Wood stork (Mycteria americana) Southeastern US 2004–2019". Movebank Data Repository. DOI: <a href="https://doi.org/10.5441/001/1.r0h6725k">10.5441/001/1.r0h6725k</a><br><br>Picardi S, Frederick PC, Borkhataria RR, Basille M. 2020. Partial migration in a subtropical wading bird in the Southeastern United States. Ecosphere. 11(2):e03054. DOI: <a href="https://doi.org/10.1002/ecs2.3054">10.1002/ecs2.3054</a><br><br>For the two particular individuals that moved to Mexico, see:<br>Picardi S, Borkhataria RR, Bryan AL, Jr, Frederick PC, Basille M. 2018. GPS telemetry reveals occasional dispersal of Wood Storks from the Southeastern US to Mexico. Caribbean Naturalist. Special Issue 2:23–29. | https://doi.org/10.5441/001/1.r0h6725k | CC_0 |
| Movebank | 1448409403 | Vanellus vanellus | Jelle Loonstra | Jelle Loonstra | | Movebank study 'Lapwing NFW Vanellus Vanellus'. Accessed on [date]. | CC_BY_NC |

| Source | Study ID | Species | Principal Investigator | Contact person | Citation | Recommended reference | License type |
|---|---|---|---|---|---|---|---|
| Movebank | 1480583194 | Hydroprogne caspia | Patrik Byholm | Patrik Byholm | Byholm P, Beal M, Lötberg U, Åkesson S. 2022. Data from: Paternal transmission of migration knowledge in a long-distance bird migrant. Movebank Data Repository. <a href="https://www.doi.org/10.5441/001/1.352qf1cv" target="_blank">https://www.doi.org/10.5441/001/1.352qf1cv</a> <br><br> Byholm P, Beal M, Isaksson N, Lötberg U, Åkesson S. 2022. Paternal transmission of migration knowledge in a long-distance bird migrant. Nat Commun. https://doi.org/10.1038/s41467-022-29300-w | https://doi.org/10.5441/001/1.352qf1cv | CC_BY |
| Movebank | 1531481854 | Phaethon aethereus,Sula dactylatra,Sula leucogaster,Fregata aquila,Onychoprion fuscatus | Diane Elizabeth Baum | Diane Elizabeth Baum | Ascension Island Government and MPIAB | Movebank study 'ICARUS Ascension Island Government'. Accessed on [date]. | CC_0 |
| Movebank | 1539583795 | Cuculus canorus | Dutch Montagu's Harrier Foundation | Dutch Montagu's Harrier Foundation | MPIAB and Dutch Montagu's Harrier Foundation | Movebank study 'ICARUS Cuckoo Holland'. Accessed on [date]. | CC_0 |
| Movebank | 1542155599 | Gallinago gallinago,Cuculus canorus | Nyambayar Batbayar | Nyambayar Batbayar | Wildlife science and conservation center of Mongolia and MPIAB | Movebank study 'ICARUS Mongolia cuckoos Nymba'. Accessed on [date]. | CC_0 |
| Movebank | 1542192442 | Streptopelia turtur | JIGUET Frédéric | Jiguet Frédéric | MNHN and MPIAB | Movebank study 'ICARUS Turtle doves'. Accessed on [date]. | CC_BY_NC |
| Movebank | 1542206798 | Cuculus canorus | JIGUET Frédéric | Jiguet Frédéric | MNHN and MPIAB | Movebank study 'ICARUS Cuckoo Fred Jiguet [ID_PROG 1189]'. Accessed on [date]. | CC_BY_NC |
| Movebank | 1543123177 | Onychoprion fuscatus | Christopher John Feare | Martin Wikelski | Seychelles Governmen, Chris Feare, Rachel Bristol, Christine Larose and MPIAB | Movebank study 'ICARUS Seychelles Sooty terns'. Accessed on [date]. | CC_0 |
| Movebank | 1562253659 | Ciconia ciconia | Wolfgang Fiedler | Wolfgang Fiedler | Data from study "LifeTrack White Stork Sarralbe" in www.movebank.org by Max Planck Institute of Animal Behavior (Radolfzell, Germany) and Comune de Sarralbe | Movebank study 'LifeTrack White Stork Sarralbe [ID_PROG 1093]'. Accessed on [date]. | CC_0 |
| Movebank | 1575973021 | Hydroprogne caspia | Susanne Akesson | Susanne Akesson | Åkesson S, Lötberg U, Rueda-Uribe C. 2022. Data from: Study "Tracking of Caspian Terns (Hydroprogne caspia) in the Swedish Baltic Sea 2017-2020". Movebank Data Repository. <a href="https://www.doi.org/10.5441/001/1.hg1v55ct" target="_blank">https://www.doi.org/10.5441/001/1.hg1v55ct</a><br><br> Rueda-Uribe C, Lötberg U, Ericsson M, Tesson SVM, Åkesson S. 2021. First tracking of declining Caspian Terns (Hydroprogne caspia) breeding in the Baltic Sea reveals high migratory dispersion and disjunct annual ranges as obstacles to effective conservation. J Avian Biol. 52(9). https://doi.org/10.1111/jav.02743 | https://doi.org/10.5441/001/1.hg1v55ct | CC_BY_NC |

| Source | Study ID | Species | Principal Investigator | Contact person | Citation | Recommended reference | License type |
|---|---|---|---|---|---|---|---|
| Movebank | 1605024900 | Chelydra serpentina,Branta canadensis,Anas platyrhynchos,Aix sponsa,Accipiter cooperii,Buteo jamaicensis,Buteo lineatus,Meleagris gallopavo,Bubo virginianus,Strix varia,Didelphis virginiana,Procyon lotor,Canis latrans | Stephen Blake | Stephen Blake | | Movebank study 'Forest Park Living Lab'. Accessed on [date]. | CC_0 |
| Movebank | 1605797471 | Haematopus ostralegus | Bruno J. Ens | Peter Desmet | Dijkstra B, Dillerop R, Oosterbeek K, Bouten W, Desmet P, van der Kolk H, Ens BJ (2021) O_ASSEN - Eurasian oystercatchers (Haematopus ostralegus, Haematopodidae) breeding in Assen (the Netherlands). Dataset. <a target="_blank" href="https://doi.org/10.5281/zenodo.5653310">https://doi.org/10.5281/zenodo.5653310</a> | https://doi.org/10.5281/zenodo.5653310 | CC_0 |
| Movebank | 1605798640 | Haematopus ostralegus | Bruno J. Ens | Peter Desmet | Dokter AM, Oosterbeek K, Baptist M, Desmet P, van der Kolk H, Bouten W, Ens BJ (2021) O_BALGZAND - Eurasian oystercatchers (Haematopus ostralegus, Haematopodidae) wintering on Balgzand (the Netherlands). Dataset. <a target="_blank" href="https://doi.org/10.5281/zenodo.5653441">https://doi.org/10.5281/zenodo.5653441</a> | https://doi.org/10.3897/zookeys.1123.90623 | CC_0 |
| Movebank | 1605799506 | Haematopus ostralegus | Bruno J. Ens | Peter Desmet | Oosterbeek K, Bom R, Shamoun-Baranes J, Desmet P, van der Kolk H, Bouten W, Ens BJ (2021) O_SCHIERMONNIKOOG - Eurasian oystercatchers (Haematopus ostralegus, Haematopodidae) breeding on Schiermonnikoog (the Netherlands). Dataset. <a target="_blank" href="https://doi.org/10.5281/zenodo.5653477">https://doi.org/10.5281/zenodo.5653477</a> | https://doi.org/10.3897/zookeys.1123.90623 | CC_0 |
| Movebank | 1605802367 | Haematopus ostralegus | Bruno J. Ens | Henk-Jan van der Kolk | van der Kolk H, Oosterbeek K, Jongejans E, Frauendorf M, Allen AM, Bouten W, Desmet P, de Kroon H, Ens BJ, van de Pol M (2021) O_VLIELAND - Eurasian oystercatchers (Haematopus ostralegus, Haematopodidae) breeding and wintering on Vlieland (the Netherlands). Dataset. <a target="_blank" href="https://doi.org/10.5281/zenodo.5653890">https://doi.org/10.5281/zenodo.5653890</a> | https://doi.org/10.1111/ibi.13035 | CC_0 |

| Source | Study ID | Species | Principal Investigator | Contact person | Citation | Recommended reference | License type |
|---|---|---|---|---|---|---|---|
| Movebank | 1605803389 | Haematopus ostralegus | Bruno J. Ens | Peter Desmet | Oosterbeek K, de Jong J, Desmet P, van der Kolk H, Bouten W, Ens BJ (2021) O_AMELAND - Eurasian oystercatchers (Haematopus ostralegus, Haematopodidae) breeding on Ameland (the Netherlands). Dataset. <a target="_blank" href="https://doi.org/10.5281/zenodo.5647596">https://doi.org/10.5281/zenodo.5647596</a> | https://doi.org/10.3897/zookeys.1123.90623 | CC_0 |
| Movebank | 1609400843 | Ichthyaetus melanocephalus | Eric Stienen | Peter Desmet | Stienen EWM, Desmet P, Milotic T, Spanoghe G, Janssens K (2022) MEDGULL_ANTWERPEN - Mediterranean gulls (Ichthyaetus melanocephalus, Laridae) breeding near Antwerp (Belgium). Dataset. <a target="_blank" href="https://doi.org/10.5281/zenodo.6599272">https://doi.org/10.5281/zenodo.6599272</a> | https://doi.org/10.5281/zenodo.6599272 | CC_0 |
| Movebank | 1623175929 | Rupicapra rupicapra | Martin Wikelski | Martin Wikelski | MPIAB and Miriam Wiesner | Movebank study 'MPIAB Wiesner Chamois Conservation '. Accessed on [date]. | CC_0 |
| Movebank | 1671751878 | Struthio camelus | Dr Willem Burger | Willem Burger | | Movebank study 'Tchad Redneck Ostrich'. Accessed on [date]. | CC_BY_NC |
| Movebank | 1701931040 | Pernis apivorus | Wolfgang Fiedler | Wolfgang Fiedler | | Movebank study 'Honey Buzzard Care Centre Release S Germany'. Accessed on [date]. | CC_0 |
| Movebank | 1720694224 | Canis latrans,Puma concolor | Julie Young | Peter Mahoney | | Movebank study 'USU: Coyote and Puma (Fishlake NF, UT)'. Accessed on [date]. | CC_0 |
| Movebank | 1724475305 | Larus michahellis | Marcin | Marcin | | Movebank study 'Yellow-legged Gulls'. Accessed on [date]. | CC_0 |
| Movebank | 1767692280 | Larus hyperboreus | Kyle Elliott | Allison Patterson | Baak JE, Patterson A, Gilchrist HG, Elliott KH. 2021. Data from: First evidence of diverging migration and overwintering strategies in glaucous gulls (Larus hyperboreus) from the Canadian Arctic. Movebank Data Repository. <a href="https://www.doi.org/10.5441/001/1.tj948m64" target="_blank">https://www.doi.org/10.5441/001/1.tj948m64</a> <br><br> Baak JE, Patterson A, Gilchrist HG, Elliott KH. 2021. First evidence of diverging migration and overwintering strategies in glaucous gulls (Larus hyperboreus) from the Canadian Arctic. Anim Migr. 8:98-109. https://doi.org/10.1515/ami-2020-0107 | https://doi.org/10.5441/001/1.tj948m64 | CC_BY |
| Movebank | 1832666571 | Caracal caracal | Laurel Serieys | Laurel Serieys | Serieys LEK, Bishop JM. 2024. Data from: Study "Caracal movement ecology study in Cape Town, South Africa". Movebank Data Repository. <a href="https://doi.org/10.5441/001/1.317" target="_blank">https://doi.org/10.5441/001/1.317</a> <br><br> These data are described in<br> Serieys LEK, Bishop JM, Rogan MS, Smith JA, Suraci JP, O'Riain MJ, Wilmers CC. 2023. Anthropogenic activities and age class mediate carnivore habitat selection in a human-dominated landscape. iScience. 26(7):107050. https://doi.org/10.1016/j.isci.2023.107050 | https://doi.org/10.5441/001/1.317 | CC_BY_NC |
| Movebank | 1841091905 | Numenius arquata | Geert Spanoghe | Peter Desmet | Spanoghe G, Janssens K, Nijs G, Milotic T, Desmet P (2021) CURLEW_VLAANDEREN - Eurasian curlews (Numenius arquata, Scolopacidae) breeding in Flanders (Belgium). Dataset. <a target="_blank" href="https://doi.org/10.5281/zenodo.5779130">https://doi.org/10.5281/zenodo.5779130</a> | https://doi.org/10.5281/zenodo.5779130 | CC_0 |

| Source | Study ID | Species | Principal Investigator | Contact person | Citation | Recommended reference | License type |
|---|---|---|---|---|---|---|---|
| Movebank | 1841261165 | Anas penelope | Jonas Waldenström | Marielle van Toor | van Toor ML, Kharitonov S, Švažas S, Dagys M, Kleyheeg E, Müskens G, Ottosson U, Žydelis R, Waldenström J. 2021. Migration distance affects how closely Eurasian wigeons follow spring phenology during migration. Mov Ecol. 9:61. doi: <a href="https://doi.org/10.1186/s40462-021-00296-0">10.1186/s40462-021-00296-0</a> | https://doi.org/10.5441/001/1.dv5mm289 | CC_BY |
| Movebank | 1852945982 | Pteropus melanotus | Christopher | Christopher | Todd CM, Westcott DA, Martin JM, Rose K, McKeown A, Hall J, Welbergen JA. 2022. Data from: Study "Movements of the Christmas Island flying fox, Australia". Movebank Data Repository. <a href="https://www.doi.org/10.5441/001/1.mn019k4d" target="_blank">https://www.doi.org/10.5441/001/1.mn019k4d</a> <br><br> These data are described in<br> Todd CM, Westcott DA, Martin JM, Rose K, McKeown A, Hall J, Welbergen JA. 2022. Body-size dependent foraging strategies in the Christmas Island flying-fox: implications for seed and pollen dispersal within a threatened island ecosystem. Mov Ecol. 10:19. https://doi.org/10.1186/s40462-022-00315-8 | https://doi.org/10.5441/001/1.mn019k4d | CC_0 |
| Movebank | 1879445455 | Chelonia mydas,Varanus salvator,Dendrocygna javanica,Anas platyrhynchos,Accipiter trivirgatus,Milvus migrans,Gyps himalayensis,Fulica atra,Numenius arquata,Tyto alba,Homo sapiens,Canis lupus,Bubo sumatranus,Nisaetus alboniger | Ratiwan Sitdhibutr | Martin Wikelski | | Movebank study 'Poultry network Thailand 2022'. Accessed on [date]. | CC_BY |
| Movebank | 1904388965 | Branta leucopsis,Anser albifrons,Anser anser | Bart Nolet | Nelleke Buitendijk | Buitendijk NH, de Jager M, Kruckenberg H, Kölzsch A, Moonen S, Müskens GJDM, Nolet BA. 2023. Data from: More grazing, more damage? Assessed damage to grassland relates non-linearly to goose grazing pressure. Movebank Data Repository. <a href="https://www.doi.org/10.5441/001/1.fk899541" target="_blank">https://www.doi.org/10.5441/001/1.fk899541</a> <br><br> These data are described in<br> Buitendijk N, de Jager M, Hornman M, Kruckenberg H, K ölzsch A, Moonen S, Nolet B. 2022. More grazing, more damage? Assessed yield loss on agricultural grassland relates non-linearly to goose grazing pressure. J Appl Ecol. 59(12): 2878-2889. https://doi.org/10.1111/1365-2664.14306 | https://doi.org/10.5441/001/1.fk899541 | CC_BY |

| Source | Study ID | Species | Principal Investigator | Contact person | Citation | Recommended reference | License type |
|---|---|---|---|---|---|---|---|
| Movebank | 1907973121 | Tapirus terrestris,calibration | Patricia Medici | Michael Noonan | Medici EP. 2023. Data from: Study "Lowland tapirs, Tapirus terrestris, in Southern Brazil". Movebank Data Repository. <a href="https://www.doi.org/10.5441/001/1.03ck4s52" target="_blank">https://www.doi.org/10.5441/001/1.03ck4s52</a> <br><br> These data are described in<br> Fleming CH, Deznabi I, Alavi S, Crofoot MC, Hirsch BT, Medici EP, Noonan MJ, Kays R, Fagan WF, Sheldon D, et al. 2022. Population-level inference for home-range areas. Methods Ecol Evol. 13(5):1027-1041. https://doi.org/10.1111/2041-210X.13815 <br><br> Medici EP, Mezzini S, Fleming CH, Calabrese JM, Noonan MJ. 2022. Movement ecology of vulnerable lowland tapirs between areas of varying human disturbance. Move Ecol. 10:14. https://doi.org/10.1186/s40462-022-00313-w | https://doi.org/10.5441/001/1.03ck4s52 | CC_BY_NC |
| Movebank | 1909487338 | Anas platyrhynchos | Martin Wikelski | Martin Wikelski | MPIAB and FAO | Movebank study 'Poultry network China 2022'. Accessed on [date]. | CC_0 |
| Movebank | 1941203363 | Aquila rapax,Trigonoceps occipitalis,Terathopius ecaudatus | Andre Botha | Martin Wikelski | Vultures for Africa and MPIAB | Movebank study 'South Africa vultures VfA MPIAB'. Accessed on [date]. | CC_0 |
| Movebank | 1958514115 | Aquila chrysaetos | Anders Tøttrup | Signe Agermose Mathiasen Andersen | | Movebank study 'The Danish Golden Eagle Project'. Accessed on [date]. | CC_BY_NC |
| Movebank | 2105214573 | Branta canadensis | Manon Sorais | Manon Sorais | Sorais M., Patenaude-Monette M., Sharp C., Askren R., LaRocque A., Leblon B., and Giroux J.-F. | Movebank study 'North-East American Canada goose migration 2015-2021'. Accessed on [date]. | CC_0 |
| Movebank | 2190520177 | Larus michahellis | Marcin | Marcin | www.interrex-tracking.com | Movebank study 'Larus michahellis X Larus cachinnans hybrids'. Accessed on [date]. | CC_0 |
| Movebank | 2272852276 | Larus fuscus | Marcin | Marcin | www.interrex-tracking.com | Movebank study 'Gulls - Minsk'. Accessed on [date]. | CC_BY_NC |
| Movebank | 2277774459 | Pelecanus occidentalis | Paul Leberg | Brock Geary | | Movebank study 'Brown Pelican Leberg Gulf of Mexico 2019'. Accessed on [date]. | CC_BY_NC |
| Movebank | 2290252202 | Branta leucopsis,Anser albifrons,Anser fabalis,Anser brachyrhynchus | Bart Nolet | Andrea Kölzsch | Kölzsch A, Lameris TK, Müskens GJDM, Schreven KHT, Buitendijk NH, Kruckenberg H, Moonen S, Heinicke T, Cao L, Madsen J, Wikelski M, Nolet BA. 2022. Data from: Wild goose chase: geese flee high and far, and with aftereffects from New Year’s fireworks. Movebank Data Repository. <a href="https://www.doi.org/10.5441/001/1.g51fs0jv" target="_blank">https://www.doi.org/10.5441/001/1.g51fs0jv</a> <br><br> These data are described in<br> Kölzsch A, Lameris TK, Müskens GJDM, Schreven KHT, Buitendijk NH, Kruckenberg H, Moonen S, Heinicke T, Cao L, Madsen J, et al. 2022. Wild goose chase: geese flee high and far, and with aftereffects from New Year’s fireworks. Conserv Lett. e12927. [illegible] | https://doi.org/10.5441/001/1.g51fs0jv | CC_BY |

| Source | Study ID | Species | Principal Investigator | Contact person | Citation | Recommended reference | License type |
|---|---|---|---|---|---|---|---|
| Movebank | 2298738353 | Larus fuscus | Eric Stienen | Peter Desmet | Stienen EWM, Müller W, Lens L, Milotic T, Desmet P (2023) LBBG_ADULT - Lesser black-backed gulls (Larus fuscus, Laridae) breeding in Belgium. Dataset. <a target="_blank" href="https://doi.org/10.5281/zenodo.10055493">https://doi.org/10.5281/zenodo.10055493</a> | https://doi.org/10.5281/zenodo.10055493 | CC_0 |
| Movebank | 2313947453 | Platalea leucorodia | Geert Spanoghe | Peter Desmet | Spanoghe G, Janssens K, Milotic T, Desmet P (2023) SPOONBILL_VLAANDEREN - Eurasian spoonbills (Platalea leucorodia, Threskiornithidae) in Flanders (Belgium). Dataset. <a target="_blank" href="https://doi.org/10.5281/zenodo.10055132">https://doi.org/10.5281/zenodo.10055132</a> | https://doi.org/10.5281/zenodo.10055132 | CC_0 |
| Movebank | 2379775078 | Hypsignathus monstrosus | Elodie Schloesing | Elodie Schloesing | Schloesing E, Caron A, Chambon R, Courbin N, Labadie M, Nina R, Mouti Mdadinga F, Ngoubili W, Sandial D, N'Kaya-Tobi , Bourgarel M, De Nys HM, Cappelle J. 2024. Data from: Foraging and mating behaviors of Hypsignathus monstrosus at the bat-human interface in central African rainforest. Movebank Data Repository. <a href="https://doi.org/10.5441/001/1.278" target="_blank">https://doi.org/10.5441/001/1.278</a> <br><br> These data are described in<br> Schloesing E, Caron A, Chambon R, Courbin N, Labadie M, Nina R, Mouiti Mbadinga F, Ngoubili W, Sandiala D, N'Kaya Tobi, Bourgarel M, De Nys HM, Cappelle J. 2023. Foraging and mating behaviors of Hypsignathus monstrosus at the bat-human interface in central African rainforest. Ecol Evol. 13(7):e10240. https://doi.org/10.1002/ece3.10240 | https://doi.org/10.5441/001/1.278 | CC_0 |
| Movebank | 2398637362 | Falco naumanni | Javier Bustamante | Javier Bustamante | | Movebank study '(EBD) Lesser Kestrel (Falco naumanni) Senegal, MERCURIO-SUMHAL'. Accessed on [date]. | CC_BY_NC |
| Movebank | 2425475445 | Gyps bengalensis,Gyps himalayensis | A B M Sarowar ALAM | A B M Sarowar ALAM | | Movebank study 'Vulture Conservation Bangladesh'. Accessed on [date]. | CC_0 |
| Movebank | 2526306516 | Mergus merganser | Anthony Wetherhill | Anthony Wetherhill | | Movebank study 'Satellite tracking of Goosander in Scotland'. Accessed on [date]. | CC_BY_NC |
| Movebank | 2608802883 | Rangifer tarandus | Leif Egil Loe | Samantha P. H. Dwinnell | Loe, L. E., B. B. Hansen, A. Stien, S. D. Albon, R. Bischof, A. Carlsson, R. J. Irvine, M. Meland, I. M. Rivrud, E. Ropstad, V. Veiberg, and A. Mysterud. 2016. Behavioral buffering of extreme weather events in a high- Arctic herbivore. Ecosphere 7(6):e01374. 10.1002/ecs2.1374 | https://doi.org/10.1002/ecs2.1374 | CC_BY_NC |
| Movebank | 2630711281 | Morus bassanus | William A. Montevecchi | Kyle d'Entremont | d'Entremont KJN, Davoren GK, Montevecchi WA. 2023. Data from: Study "Northern Gannet Breeding Season GPS Data from Cape St. Mary's, NL, Canada: 2019 to 2022". Movebank Data Repository. <a href="https://www.doi.org/10.5441/001/1.5km7v2s3" target="_blank">https://www.doi.org/10.5441/001/1.5km7v2s3</a> <br><br> These data are described in<br> d'Entremont KJN, Davoren GK, Walsh CJ, Wilhelm SI, Montevecchi WA. 2022. Intra- and inter-annual shifts in foraging tactics by parental northern gannets Morus bassanus indicate changing prey fields. Mar Ecol Prog Ser. 698:155-170. https://www.doi.org/10.3354/meps14164 | https://doi.org/10.5441/001/1.5km7v2s3 | CC_BY_NC |

| Source | Study ID | Species | Principal Investigator | Contact person | Citation | Recommended reference | License type |
|---|---|---|---|---|---|---|---|
| Movebank | 2636372210 | Lynx rufus,Canis latrans | Laura Prugh | Laura Prugh | Prugh LR. 2023. Data from: Study "GPS tracking of bobcats and coyotes in northern Washington". Movebank Data Repository. <a href="https://www.doi.org/10.5441/001/1.gm93267b" target="_blank">https://www.doi.org/10.5441/001/1.gm93267b</a> <br><br> These data are described in<br> Prugh LR, Cunningham CX, Windell RM, Kertson BN, Ganz TR, Walker SL, Wirsing AJ. 2023. Fear of large carnivores amplifies human-caused mortality for mesopredators. Science. 380(6646):754-758. https://doi.org/10.1126/science.adf2472 <br><br> Bassing SB, DeVivo M, Ganz TR, Kertson BN, Prugh LR, Roussin T, Satterfield L, Windell RM, Wirsing AJ, Gardner B. 2023. Are we telling the same story? Comparing inferences made from camera trap and telemetry data for wildlife monitoring. Ecol Appl. 33(1):e2745. https://doi.org/10.1002/eap.2745 | https://doi.org/10.5441/001/1.gm93267b | CC_BY |
| Movebank | 2658117564 | Morus bassanus | Jana W E Jeglinski | Jana W E Jeglinski | Jeglinski JWE, Matthiopoulos J, Votier SC, Lane JV. 2024. Data from: Strong breeding colony fidelity in northern gannets following high pathogenicity avian influenza HPAIV outbreak. Movebank Data Repository. <a href="https://doi.org/10.5441/001/1.608" target="_blank">https://doi.org/10.5441/001/1.608</a> <br><br> These data are described in<br>Grémillet D, Ponchon A, Provost P, Gamble A, Abed-Zahar M, Bernard A, Courbin N, Delavaud G, Deniau A, Fort J, et al. 2023. Strong breeding colony fidelity in northern gannets following high pathogenicity avian influenza HPAIV outbreak. Biol Conserv. 286:110269. https://doi.org/10.1016/j.biocon.2023.110269<br><br> Jeglinski JWE, Lane JV, Votier SC, Furness RW, Hamer KC, McCafferty DJ, Nager RG, Sheddan M, Wanless S, Matthiopoulos J. 2024. HPAIV outbreak triggers short-term colony connectivity in a seabird metapopulation. Sci Rep. 14:3126. https://doi.org/10.1038/s41598-024-53550-x | https://doi.org/10.5441/001/1.608 | CC_BY |
| Movebank | 2658220054 | Morus bassanus | Jana W E Jeglinski | Jana W E Jeglinski | Jeglinski JWE, Lane JV, Votier SC, Furness RW, Hamer KC, McCafferty DJ, Nager R, Sheddan M, Wanless S, Matthiopoulos J. 2024. Data from: HPAIV outbreak triggers short-term colony connectivity in a seabird metapopulation. Movebank Data Repository. <a href="https://doi.org/10.5441/001/1.607" target="_blank">https://doi.org/10.5441/001/1.607</a> <br><br> These data are described in<br> Jeglinski JWE, Lane JV, Votier SC, Furness RW, Hamer KC, McCafferty DJ, Nager RG, Sheddan M, Wanless S, Matthiopoulos J. 2024. HPAIV outbreak triggers short-term colony connectivity in a seabird metapopulation. Sci Rep. 14:3126. https://doi.org/10.1038/s41598-024-53550-x | https://doi.org/10.5441/001/1.607 | CC_BY_NC |
| Movebank | 2830157729 | Nycticorax nycticorax | Mike Ward | Sarah Slayton | | Movebank study 'Chicago BCNH Project'. Accessed on [date]. | CC_0 |
| Movebank | 2915700754 | Bubulcus ibis | Benjamin Vollot | Benjamin Vollot | | Movebank study 'Bubulcus ibis_Vollot_ENVT_65_ID_PROG1035'. Accessed on [date]. | CC_0 |

| Source | Study ID | Species | Principal Investigator | Contact person | Citation | Recommended reference | License type |
|---|---|---|---|---|---|---|---|
| Movebank | 2944153255 | Larus smithsonianus | Mark Mallory | Sarah Gutowsky | Mallory ML, Craik S, Allard KA, Gutowsky S. 2023. Data from: Study "American Herring Gulls - GPS - Lobster Bay, Southwest Nova Scotia, Canada". Movebank Data Repository. <a href="https://www.doi.org/10.5441/001/1.292" target="_blank">https://www.doi.org/10.5441/001/1.292</a> <br><br> These data are described in<br> Gutowsky S, Baak JE, Craik S, Mallory M, Knutson N, d'Entremont A, Allard K. 2023. Seasonal and circadian patterns of herring gull (Larus smithsoniansus) movements reveal temporal shifts in industry and coastal island interaction. Ecol Solut Evid. 4(3):e12274. https://doi.org/10.1002/2688-8319.12274 | https://doi.org/10.5441/001/1.292 | CC_0 |
| Movebank | 2961927604 | Falco naumanni | Javier Bustamante | Javier Bustamante | Lesser kestrel movements, EBD-CSIC, projects MERCURIO-SUMHAL | Movebank study '(EBD) Lesser Kestrel (Falco naumanni) Spain, MERCURIO-SUMHAL'. Accessed on [date]. | CC_BY_NC |
| Movebank | 2970193504 | Falco tinnunculus | Javier Bustamante | Javier Bustamante | Common kestrel (Falco tinnunculus) movements, EBD-CSIC, projects MERCURIO & SUMHAL | Movebank study '(EBD) Common Kestrel (Falco tinnunculus) Spain, MERCURIO-SUMHAL'. Accessed on [date]. | CC_BY_NC |
| Movebank | 2976678046 | Odocoileus virginianus | Blaise Ashley Newman | Blaise Ashley Newman | Newman BA, Dyal JR, Miller KV, Cherry MJ, D'Angelo GJ. 2023. Data from: Influence of visual perception on movement decisions by an ungulate prey species. Movebank Data Repository. <a href="https://www.doi.org/10.5441/001/1.293" target="_blank">https://www.doi.org/10.5441/001/1.293</a> <br><br> These data are described in<br> Newman BA, Dyal JR, Miller KV, Cherry MJ, D'Angelo GJ. 2023. Influence of visual perception on movement decisions by an ungulate prey species. Biol Open. 12(10):bio059932. https://doi.org/10.1242/bio.059932 <br><br> Dyal JR, Miller KV, Cherry MJ, D'Angelo GJ. 2022. White-tailed deer movement in response to helicopter surveys. Wildlife Soc Bull. 46(5):e1383. https://doi.org/10.1002/wsb.1383 | https://doi.org/10.5441/001/1.293 | CC_BY |
| Movebank | 2984378217 | Anthropoides virgo | Yuriy | Ivan Pokrovskiy | Andryushchenko Y, Pokrovsky I, Bernd V, Fiedler W, Wikelski M. 2024. Data from: Study "1000 Cranes. Ukraine.". Movebank Data Repository. <a href="https://doi.org/10.5441/001/1.590" target="_blank">https://doi.org/10.5441/001/1.590</a> <br><br> These data are described in<br> Yanco SW, Oliver RY, Iannarilli F, Carlson BS, Heine G, Mueller U, Richter N, Vorneweg B, Andryushchenko Y, Batbayar N, et al. 2024. Migratory birds modulate niche tradeoffs in rhythm with seasons and life history. Proc Natl Acad Sci USA. https://doi.org/10.1073/pnas.2316827121 | https://doi.org/10.5441/001/1.590 | CC_BY |
| Movebank | 3094468724 | Neophron percnopterus | Ron Efrat | Ron Efrat | Efrat R, Hatzofe O, Mueller T, Sapir N, Berger-Tal O. 2023. Data from: Early life and acquired experiences interact in shaping migratory and flight behaviors. Movebank Data Repository. <a href="https://www.doi.org/10.5441/001/1.298" target="_blank">https://www.doi.org/10.5441/001/1.298</a> <br><br> These data are described in<br> Efrat R, Hatzofe O, Mueller T, Sapir N, Berger-Tal O. 2023. Early life and acquired experiences interact in shaping migratory and flight behaviors. Curr Biol. https://doi.org/10.1016/j.cub.2023.11.012 | https://doi.org/10.5441/001/1.298 | CC_BY |

| Source | Study ID | Species | Principal Investigator | Contact person | Citation | Recommended reference | License type |
|---|---|---|---|---|---|---|---|
| Movebank | 3131029654 | Milvus milvus | Jan Skrabal | Jan Skrabal | Škrábal J, Literák I, Raab R. 2023. Study "Milvus_milvus_Soaring_over_Adriatic_sea". Movebank Data Repository. <a href="https://doi.org/10.5441/001/1.300" target="_blank">https://doi.org/10.5441/001/1.300</a> <br><br> These data are described in<br> Škrábal J, Krejčí Š, Raab R, Sebastián-González E, Literák I. 2023. Soaring over open waters: horizontal winds provide lift to soaring migrants in weak thermal conditions. Movement Ecol. 11:76. https://doi.org/10.1186/s40462-023-00438-6 | https://doi.org/10.5441/001/1.300 | CC_BY_NC |
| Movebank | 3133580132 | Lepus europaeus | Chloe tavernier | Chloe tavernier | | Movebank study 'Lepus.europanus-Tavernier-SolarPark'. Accessed on [date]. | CC_BY_NC |
| Movebank | 3179890710 | Vulpes vulpes | Tom Porteus | Tom Porteus | Porteus TA, Short MJ, Hoodless AN, Reynolds JC. 2024. Data from: Study "Red Fox (Vulpes vulpes) in UK wet grasslands". Movebank Data Repository. <a href="https://doi.org/10.5441/001/1.304" target="_blank">https://doi.org/10.5441/001/1.304</a> <br><br> These data are described in<br> Porteus TA, Short MJ, Hoodless AN, Reynolds JC. 2024. Movement ecology and minimum density estimates of red foxes in wet grassland habitats used by breeding wading birds. Eur J Wildlife Res. 70:8. https://doi.org/10.1007/s10344-023-01759-y | https://doi.org/10.5441/001/1.304 | CC_0 |
| Movebank | 3186161092 | Anatidae ,Dendrocygna bicolor | Manuel Olivier Grosselet | Manuel Olivier Grosselet | | Movebank study 'Dendrocygna bicolor movement from Mexico City Manuel Grosselet'. Accessed on [date]. | CC_0 |
| Movebank | 3218698288 | Rynchops niger | Manuel Olivier Grosselet | Manuel Olivier Grosselet | | Movebank study 'rynchops niger'. Accessed on [date]. | CC_0 |
| Movebank | 3367931169 | Nucifraga caryocatactes | Eike Lena Neuschulz | Eike Lena Neuschulz | Graf V, Sorensen MC, Mueller T, Neuschulz EL. 2024. Data from: Study "Movement patterns of seed dispersing spotted nutcrackers (Nucifraga caryocatactes)". Movebank Data Repository. <a href="https://doi.org/10.5441/001/1.324" target="_blank">https://doi.org/10.5441/001/1.324</a> <br><br> These data are described in<br> Sorensen MC, Mueller T, Donoso I, Graf V, Merges D, Vanoni M, Fiedler W, Neuschulz EL. Scatter-hoarding birds disperse seeds to sites unfavorable for plant regeneration. Mov Ecol. 10:38. https://doi.org/10.1186/s40462-022-00338-1 <br><br> Graf V, Mueller T, Grüebler MU, Kormann UG, Albrecht J, Hertel AG, Sorensen MC, Tschumi M, Neuschulz EL. In press, 2024. Individual behavior shapes patterns of bird-mediated seed dispersal. Funct Ecol. | https://doi.org/10.5441/001/1.324 | CC_BY |
| Movebank | 3413045568 | Streptopelia turtur | JIGUET Frédéric | Andrea Kölzsch | HABITRACK: Habitat tracking for the conservation of huntable bird species | Movebank study 'Habitrack European Turtle Dove'. Accessed on [date]. | CC_BY |
| Movebank | 3578422072 | Tyto furcata | Hermann Wagner | Hermann Wagner | | Movebank study 'Galapagos barn owl project'. Accessed on [date]. | CC_0 |
| Movebank | 3791354435 | Panthera leo | Kevin MacFarlane | Kevin MacFarlane | MacFarlane, Kevin, 2014, The Ecology and Management of Kalahari lions in a Conflict Area in Central Botswana, PhD, Australian National University | https://doi.org/10.25911/5d78d54c1a50c | CC_0 |
| Movebank | 3809257699 | Panthera leo | Kevin MacFarlane | Robert Heinsohn | MacFarlane K. 2014. The ecology and management of Kalahari lions in a conflict area in central Botswana [dissertation]. [Canberra (Australia)]: Australian National University. https://doi.org/10.25911/5d78d54c1a50c | https://doi.org/10.25911/5d78d54c1a50c | CC_BY |

| Source | Study ID | Species | Principal Investigator | Contact person | Citation | Recommended reference | License type |
|---|---|---|---|---|---|---|---|
| Movebank | 3902656356 | Nycticorax nycticorax | Mike Ward | Sarah Slayton | | Movebank study 'Chicago BCNH Project (Druid)'. Accessed on [date]. | CC_0 |
| Movebank | 3945977313 | Aphriza virgata | Shawn Crimmins | Sam Simon | | Movebank study 'Surfbird_UAF_BLM_InteriorAlaska'. Accessed on [date]. | CC_0 |
| Movebank | 4447816442 | Rissa tridactyla | Jacob Davies | Jacob Davies | | Movebank study 'BTO - Whinnyfold 2021 - Kittiwake'. Accessed on [date]. | CC_0 |
| Movebank | 4645303014 | Vultur gryphus | Hannah Williams | Hannah Williams | | Movebank study 'Juvenile Condor Cordoba Tatu Carreta Reserve'. Accessed on [date]. | CC_0 |
| Movebank | 4725153649 | Choloepus hoffmanni | Amelia Symeou | Amelia Symeou | | Movebank study 'Choloepus hoffmanni, The Sloth Conservation Foundation, Puerto Viejo de Talamanca Costa Rica'. Accessed on [date]. | CC_0 |
| Movebank | 4732684199 | Bradypus variegatus | Rebecca Cliffe | Amelia Symeou | | Movebank study 'Bradypus variegatus, The Sloth Conservation Foundation, Puerto Viejo de Talamanca Costa Rica'. Accessed on [date]. | CC_0 |
| Movebank | 4739876185 | Rallus ,Rallus elegans | Richard Temple Jr. | Richard Temple Jr. | | Movebank study 'KIRA SW LA'. Accessed on [date]. | CC_0 |
| Movebank | 4901146318 | Connochaetes taurinus | Grant Hopcraft | Tom Morrison | Hopcraft, J.G.C., Morrison, T.A., and Stabach, J.A. (2024). Data from: Movement of resident wildebeest across the Masai Mara Ecosystem, Kenya. Movebank Data Repository. | Movebank study 'White-bearded wildebeest (Connochaetes taurinus) - Greater Mara Ecosystem (2017-2021)'. Accessed on [date]. | CC_BY_NC |
| Movebank | 5175345606 | Lynx rufus,Urocyon cinereoargenteus | Laurel Serieys | Laurel Serieys | Serieys LEK, Matsushima SS, Wilmers CC. 2024. Data from: Study "Bobcat habitat connectivity study in central California". Movebank Data Repository. <a href="https://doi.org/https://doi.org/10.5441/001/1.323" target="_blank">https://doi.org/https://doi.org/10.5441/001/1.323</a> <br><br> These data are described in<br> Serieys LEK, Rogan MS, Matsushima SS, Wilmers CC. 2021. Road-crossings, vegetative cover, land use and poisons interact to influence corridor effectiveness. Biol Conserv. 253:108930. https://doi.org/10.1016/j.biocon.2020.108930. | https://doi.org/10.5441/001/1.323 | CC_0 |
| Tucker at al 2023 | | Alces alces | Bram Van Moorter | | | | |
| Tucker at al 2023 | | Alces alces | Matthew Kauffman | | | | |
| Tucker at al 2023 | | Antilocapra americana | Chris Geremia | | | | |
| Tucker at al 2023 | | Antilocapra americana | Julie Young | | | | |
| Tucker at al 2023 | | Antilocapra americana | Matthew Kauffman | | | | |
| Tucker at al 2023 | | Canis aureus | Hubert Potocnik | | | | |

| Source | Study ID | Species | Principal Investigator | Contact person | Citation | Recommended reference | License type |
|---|---|---|---|---|---|---|---|
| Tucker at al 2023 | | Canis latrans | David Drake | | | | |
| Tucker at al 2023 | | Canis lupus | Daniel Stahler | | | | |
| Tucker at al 2023 | | Canis lupus | Jerrold Belant | | | | |
| Tucker at al 2023 | | Canis lupus | Mark Hebblewhite | | | | |
| Tucker at al 2023 | | Capreolus capreolus | Francesca Cagnacci | | | | |
| Tucker at al 2023 | | Cervus canadensis | Daniel Stahler | | | | |
| Tucker at al 2023 | | Cervus canadensis | Julie Young | | | | |
| Tucker at al 2023 | | Cervus canadensis | Kim Poole | | | | |
| Tucker at al 2023 | | Cervus elaphus | Erling Meisingset | | | | |
| Tucker at al 2023 | | Cervus elaphus | Johannes Signer | | | | |
| Tucker at al 2023 | | Cervus elaphus | Marco Heurich | | | | |
| Tucker at al 2023 | | Cervus elaphus | Peter Sunde | | | | |
| Tucker at al 2023 | | Chrysocyon brachyurus | Rogerio Cunha de Paula | | | | |
| Tucker at al 2023 | | Connochaetes taurinus | Grant Hopcraft | | | | |
| Tucker at al 2023 | | Equus hemionus | Petra Kaczensky / Buuveibaatar Bayarbaatar | | | | |
| Tucker at al 2023 | | Loxodonta africana | Morgan Hauptfleisch | | | | |
| Tucker at al 2023 | | Loxodonta africana | Robert Pringle | | | | |
| Tucker at al 2023 | | Odocoileus hemionus | Chris Geremia | | | | |

| Source | Study ID | Species | Principal Investigator | Contact person | Citation | Recommended reference | License type |
|---|---|---|---|---|---|---|---|
| Tucker at al 2023 | | Odocoileus hemionus | Julie Young | | | | |
| Tucker at al 2023 | | Odocoileus hemionus | Matthew Kauffman | | | | |
| Tucker at al 2023 | | Odocoileus virginianus | Guillaume Bastille-Rousseau | | | | |
| Tucker at al 2023 | | Odocoileus virginianus | Jerrold Belant | | | | |
| Tucker at al 2023 | | Odocoileus virginianus | Vickie DeNicola | | | | |
| Tucker at al 2023 | | Ovis canadensis | Chris Geremia | | | | |
| Tucker at al 2023 | | Ovis canadensis | Kevin Monteith | | | | |
| Tucker at al 2023 | | Ovis canadensis californiana | Julie Young | | | | |
| Tucker at al 2023 | | Ovis canadensis canadensis | Julie Young | | | | |
| Tucker at al 2023 | | Ovis canadensis nelsoni | Julie Young | | | | |
| Tucker at al 2023 | | Panthera leo | Jacob Goheen | | | | |
| Tucker at al 2023 | | Procapra gutturosa | Nandintsetseg Dejid | | | | |
| Tucker at al 2023 | | Puma concolor | Christopher Wilmers | | | | |
| Tucker at al 2023 | | Puma concolor | Julie Young | | | | |
| Tucker at al 2023 | | Rangifer tarandus | Allicia Kelly | | | | |
| Tucker at al 2023 | | Rangifer tarandus | Bram Van Moorter | | | | |
| Tucker at al 2023 | | Tragelaphus angasii | Robert Pringle | | | | |
| Tucker at al 2023 | | Tragelaphus strepsiceros | Robert Pringle | | | | |

| Source | Study ID | Species | Principal Investigator | Contact person | Citation | Recommended reference | License type |
|---|---|---|---|---|---|---|---|
| Tucker at al 2023 | | Tragelaphus sylvaticus | Robert Pringle | | | | |
| Tucker at al 2023 | | Ursus arctos | Jerrold Belant | | | | |
| Tucker at al 2023 | | Ursus arctos | Jonas Kindberg | | | | |
| Tucker at al 2023 | | Ursus arctos | Nuria Selva | | | | |

| Source | Study ID | Species | Principal Investigator | Contact person | Citation | License type |
|---|---|---|---|---|---|---|
| Movebank | 82684 | Bycanistes bucinator | Wolfgang Fiedler | Wolfgang Fiedler | Lenz J, Fiedler W, Caprano T, Friedrichs W, Gaese BH, Wikelski M, Böhning-Gaese K. 2011. Seed-dispersal distributions by trumpeter hornbills in fragmented landscapes. Proceedings of the Royal Society B Biological Sciences. https://doi.org/10.1098/rspb.2010.2383 | CC_BY |
| Movebank | 446579 | Anas platyrhynchos | Martin Wikelski | Wolfgang Fiedler | Korner P, Sauter A, Fiedler W, Jenni L. 2016. Variable allocation of activity to daylight and night in the mallard. Animal Behaviour. 115: 69–79. https://doi.org/10.1016/j.anbehav.2016.02.026 | CC_BY |
| Movebank | 481458 | Cathartes aura,Coragyps atratus | David Barber | David Barber | Bildstein KL, Barber D, Bechard MJ, Graña Grilli M, Therrien J. 2021. Data from: Study "Vultures Acopian Center USA GPS" (2003-2021). Movebank Data Repository. <a href="https://www.doi.org/10.5441/001/1.f3qt46r2" target="_blank">https://www.doi.org/10.5441/001/1.f3qt46r2</a> <br><br> Mallon JM, Bildstein KL, Fagan WF. 2021. Inclement weather forces stopovers and prevents migratory progress for obligate soaring migrants. Mov Ecol. 9:39. https://doi.org/10.1186/s40462-021-00274-6 <br><br> Graña Grilli M, Lambertucci SA, Therrien J-F, Bildstein KL. 2017. Wing size but not wing shape is related to migratory behavior in a soaring bird. J Avian Biol. 48(5):669-678. https://doi.org/10.1111/jav.01220 <br><br> Dodge S, Bohrer G, Bildstein K, Davidson SC, Weinzierl R, Mechard MJ, Barber D, Kays R, Brandes D, Han J, et al. 2014. Environmental drivers of variability in the movement ecology of turkey vultures (Cathartes aura) in North and South America. Philos T Roy Soc B. 369(1643):20130195. https://doi.org/10.1098/rstb.2013.0195 <br><br> Portions of these data are published with DOIs 10.5441/001/1.37r2b884 (study "Vultures Acopian Center USA 2003-2016") and 10.5441/001/1.46ft1k05 (study "Turkey vultures in North and South America (data from Dodge et al. 2014)") | CC_BY |

| | | | | | | |
|---|---|---|---|---|---|---|
| Movebank | 1764627 | Syncerus caffer | Paul Cross | Paul Cross | Cross PC, Bowers JA, Hay CT, Wolhuter J, Buss P, Hofmeyr M, du Toit JT, Getz WM. 2016. Data from: Nonparameteric kernel methods for constructing home ranges and utilization distributions. Movebank Data Repository. <a href="https://www.doi.org/10.5441/001/1.j900f88t" target="_blank">https://www.doi.org/10.5441/001/1.j900f88t</a> <br><br> Calabrese JM, Fleming CH, Gurarie E. 2016. ctmm: an r package for analyzing animal relocation data as a continuous-time stochastic process. Methods Ecol Evol. 7(9):1124-1132. https://doi.org/10.1111/2041-210X.12559 <br><br> Cross PC, Heisey DM, Bowers JA, Hay CT, Wolhuter J, Buss P, Hofmeyr M, Michel AL, Bengis RG, Bird TLF, et al. 2009. Disease, predation and demography: assessing the impacts of bovine tuberculosis on African buffalo by monitoring at individual and population levels. J Appl Ecol. 46(2):467-475. https://doi.org/10.1111/j.1365-2664.2008.01589.x <br><br> Getz WM, Fortmann-Roe S, Cross PC, Lyons AJ, Ryan SJ, Wilmers CC. 2007. LoCoH: Nonparameteric kernel methods for constructing home ranges and utilization distributions. PLoS ONE. 2(2):e207. https://doi.org/10.1371/journal.pone.0000207 | CC_0 |
| Movebank | 1818825 | Loxodonta cyclotis | Stephen Blake | Stephen Blake | | CC_BY |
| Movebank | 1902221 | Pecari tajacu | Roland Kays | Roland Kays | No papers published | CC_BY |

| | | | | | | |
|---|---|---|---|---|---|---|
| Movebank | 2919708 | Gyps africanus,Torgos tracheliotus | Ran Nathan | Orr Spiegel | Spiegel O, Getz WM, Nathan R. 2014. Data from: Factors influencing foraging search efficiency: Why do scarce lappet-faced vultures outperform ubiquitous white-backed vultures? (V2). Movebank Data Repository. <a href="https://www.doi.org/10.5441/001/1.mf903197" target="_blank">https://www.doi.org/10.5441/001/1.mf903197</a> <br><br> Spiegel O, Getz WM, Nathan R. 2013. Factors influencing foraging search efficiency: why do scarce lappet-faced vultures outperform ubiquitous white-backed vultures? Am Nat. 181(5):E102-115. https://www.doi.org/10.1086/670009 | CC_0 |
| Movebank | 2927282 | Anas platyrhynchos,Anas penelope,Aythya ferina | Wolfgang Fiedler | Wolfgang Fiedler | Data by Max Planck Institute for Animal Behavior, Vogelwarte Radolfzell, Germany. | CC_BY |
| Movebank | 2930072 | Steatornis caripensis | Martin Wikelski | Martin Wikelski | Holland RA, Wikelski M, Kuemmeth F, Bosque C. 2012. Data from: The secret life of oilbirds: new insights into the movement ecology of a unique avian frugivore. Movebank Data Repository. <a href="https://www.doi.org/10.5441/001/1.35fs26kq" target="_blank">https://www.doi.org/10.5441/001/1.35fs26kq</a> <br><br> Holland RA, Wikelski M, Kuemmeth F, Bosque C. 2009. The secret life of oilbirds: new insights into the movement ecology of a unique avian frugivore. PLoS ONE. 4(12):e8264. https://doi.org/10.1371/journal.pone.0008264 | CC_0 |

| | | | | | | |
|---|---|---|---|---|---|---|
| Movebank | 2943485 | Aquila chrysaetos,Haliaeetus leucocephalus,Pandion haliaetus,Falco peregrinus,Asio flammeus | Roland Kays | Roland Kays | Nye P, Hewitt G, Swenson T, Kays R. 2018. Data from: New York State bald eagle report 2010. Movebank Data Repository. <a href="https://www.doi.org/10.5441/001/1.s65q50j0" target="_blank">https://www.doi.org/10.5441/001/1.s65q50j0</a> <br><br> Li Z, Han J, Ding B, Kays R. 2012. Mining periodic behaviors of object movements for animal and biological sustainability studies. Dat Min Knowl Disc. 24(2):355-386. https://doi.org/10.1007/s10618-011-0227-9 <br><br> Nye P. 2010. New York State bald eagle report 2010. Albany (NY): New York State Department of Environmental Conservation. 43 p. <br><br> Martell MS, Henny CJ, Nye PE, Solensky MJ. 2001. Fall migration routes, timing, and wintering sites of North American ospreys as determined by satellite telemetry. Condor. 103(4):715-724. https://doi.org/10.1650/0010-5422(2001)103[0715:FMRTAW]2.0.CO;2 <br><br> Rodriguez F, Martell M, Nye P, Bildstein KL. 2001. Osprey migration through Cuba. In: Bildstein K, Klem D Jr, editors. Hawkwatching in the Americas. North Wales (PA): Hawk Migration Association of North America. p. 107-117. <br><br> Bautz L, Nye P. 1987. Spring movement of an adult bald eagle from southeastern New York to central Ontario. The Eyas. 10(1):32-33. | CC_0 |
| Movebank | 2980294 | Anas strepera | Andrea Gehrold | Wolfgang Fiedler | Gehrold A, Bauer H-G, Fiedler W, Wikelski M. 2014. Great flexibility in autumn movement patterns of European Gadwalls (Anas strepera). Journal of Avian Biology. 45: 131–139. https://doi.org/10.1111/j.1600-048X.2013.00248.x<br><br>A subset of these data have been published as<br> Gehrold A, Wikelski M. 2013. Data from: Great flexibility in autumn movement patterns of European Gadwalls (Anas strepera). Movebank Data Repository. <a href="https://doi.org/10.5441/001/1.26dg08hv" target="_blank">https://doi.org/10.5441/001/1.26dg08hv</a> | CC_BY |

| | | | | | | |
|---|---|---|---|---|---|---|
| Movebank | 2988309 | Phalacrocorax carbo | Wolfgang Fiedler | Wolfgang Fiedler | | CC_BY |
| Movebank | 2988333 | Larus fuscus | Martin Wikelski | Martin Wikelski | Wikelski M, Arriero E, Gagliardo A, Holland R, Huttunen MJ, Juvaste R, Mueller I, Tertitski G, Thorup K, Wild M, Alanko M, Bairlein F, Cherenkov A, Cameron A, Flatz R, Hannila J, Hüppop O, Kangasniemi M, Kranstauber B, Penttinen M, Safi K, Semashko V, Schmid H, Wistbacka R. 2015. Data from: True navigation in migrating gulls requires intact olfactory nerves. Movebank Data Repository. <a href="https://doi.org/10.5441/001/1.q986rc29" target="_blank">https://doi.org/10.5441/001/1.q986rc29</a> <br><br> Wikelski M, Arriero E, Gagliardo A, Holand R, Huttunen MJ, Juvaste R, Mueller I, Tertitski G, Thorup K, Wild M, et al. 2015. True navigation in migrating gulls requires intact olfactory nerves. Sci Rep. 5:17061. https://doi.org/10.1038/srep17061 | CC_0 |
| Movebank | 3109235 | Anas platyrhynchos | Jonas Waldenström | Jonas Waldenström | Bengtsson, D., Safi, K., Avril, A., Fiedler, W., Wikelski, M., Gunnarsson, G., Elmberg, J., Tolf, C., Olsen, B. & Waldenström, J. 2016. Does influenza A virus infection affect movement behaviour during stopover in its wild reservoir host? Royal Society Open Science 3: 150633. [http://dx.doi.org/10.1098/rsos.150633]. <br ><br /> Bengtsson, D., Avril, A., Gunnarsson, G., Elmberg, J., Söderquist, P., Norevik, G., Tolf, C., Safi, K., Fiedler, W., Wikelski, M., Olsen, B. & Waldenström, J. 2014. Movements, home-range size and habitat selection of Mallards during fall migration. PLOS ONE 9: 00764 [doi: 10.1371/journal.pone.0100764]<br /><br /> | CC_BY_NC |
| Movebank | 3780829 | Bubo bubo | Martin Wikelski | Wolfgang Fiedler | Reinhard Vohwinkel et al. in prep. | CC_BY |

| | | | | | | |
|---|---|---|---|---|---|---|
| Movebank | 4124813 | Grus nigricollis | Sherub Sherub | Sherub Sherub | Sherub S, Wikelski M. 2024. Data from: Study "Black-necked crane Bhutan (UWICE-MPIAB)". Movebank Data Repository. <a href="https://doi.org/10.5441/001/1.601" target="_blank">https://doi.org/10.5441/001/1.601</a> <br><br> These data are described in<br> Yanco SW, Oliver RY, Iannarilli F, Carlson BS, Heine G, Mueller U, Richter N, Vorneweg B, Andryushchenko Y, Batbayar N, et al. 2024. Migratory birds modulate niche tradeoffs in rhythm with seasons and life history. Proc Natl Acad Sci USA. https://doi.org/10.1073/pnas.2316827121 | CC_BY |
| Movebank | 6770990 | Fregata magnificens | Martin Wikelski | Martin Wikelski | Wikelski et al., birds and hurricanes, in prep. | CC_BY |
| Movebank | 6925808 | Martes pennanti | Scott LaPoint | Scott LaPoint | Paper: LaPoint, S, Gallery P, Wikelski M, Kays R (2013) Animal behavior, cost-based corridor models, and real corridors. Landscape Ecology, v 28 i 8, p 1615–1630. doi:10.1007/s10980-013-9910-0 LaPoint S, Gallery P, Wikelski M, Kays R. 2013. Data from: Animal behavior, cost-based corridor models, and real corridors. Movebank Data Repository. https://doi.org/10.5441/001/1.2tp2j43g | CC_BY_NC |
| Movebank | 7002955 | Ciconia ciconia | Ran Nathan | Shay Rotics | Rotics et al. in prep | CC_BY |

| | | | | | | |
|---|---|---|---|---|---|---|
| Movebank | 7023252 | Papio anubis | Meg Crofoot | Meg Crofoot | Crofoot MC, Kays RW, Wikelski M. 2021. Data from: Study "Collective movement in wild baboons". Movebank Data Repository. <a href="https://www.doi.org/10.5441/001/1.3q2131q5" target="_blank">https://www.doi.org/10.5441/001/1.3q2131q5</a> <br><br> Harel R, Loftus JC, Crofoot MC. 2021. Locomotor compromises maintain group cohesion in baboon troops on the move. P Roy Soc B. 288(1955):20210839. https://doi.org/10.1098/rspb.2021.0839 <br><br> Farine DR, Strandburg-Peshkin A, Couzin ID, Berger-Wolf TY, Crofoot MC. 2017. Individual variation in local interaction rules can explain emergent patterns of spatial organization in wild baboons. P Roy Soc B. 284(1853):20162243. https://doi.org/10.1098/rspb.2016.2243 <br><br> Strandburg-Peshkin A, Farine DR, Crofoot MC, Couzin ID. 2017. Habitat and social factors shape individual decisions and emergent group structure during baboon collective movement. eLife. 6:e19505. https://doi.org/10.7554/eLife.19505 <br><br> Farine DR, Strandburg-Peshkin A, Berger-Wolf T, Ziebart B, Brugere I, Li J, Crofoot MG. 2016. Both nearest neighbours and long-term affiliates predict individual locations during collective movement in wild baboons. Sci Rep. 6:27704. https://doi.org/10.1038/srep27704 <br><br> Strandburg-Peshkin A, Farine DR, Couzin ID, Crofoot MC. 2015. Shared decision-making drives collective movement in wild baboons. Science. 348–6241:1358–1361. https://doi.org/10.1126/science.aaa5099 | CC_0 |
| Movebank | 7431347 | Ciconia ciconia | Movebank Administrator | Wolfgang Fiedler | Berthold P, Kaatz C, Kaatz M, Querner U, van den Bossche W, Chernetsov N, Fiedler W, Wikelski M. 2022. Data from: Study "MPIAB Argos white stork tracking (1991-2017)". Movebank Data Repository. <a href="https://www.doi.org/10.5441/001/1.k29d81dh" target="_blank">https://www.doi.org/10.5441/001/1.k29d81dh</a> | CC_BY |

| | | | | | | |
|---|---|---|---|---|---|---|
| Movebank | 8019591 | Bison bison | Stephen Blake | Stephen Blake | | CC_BY |
| Movebank | 8849813 | Ardea alba | Roland Kays | Roland Kays | Brzorad, J. N., M. C. Allen, S. Jennings, E. Condeso, S. Elbin, R. Kays, D. Lumpkin, S. Schweitzer, N. Tsipoura, and A. D. Maccarone. 2022. Seasonal Patterns in Daily Flight Distance and Space Use by Great Egrets (Ardea alba). Waterbirds 44:343–355. | CC_0 |
| Movebank | 8927992 | Anas platyrhynchos,Fulica atra | Martin Wikelski | Martin Wikelski | Wikelski et al., unpublished | CC_BY |
| Movebank | 9493881 | Ciconia ciconia | Martin Wikelski | Wolfgang Fiedler | Data from study "LifeTrack White Stork Uzbekistan" in www.movebank.org by Max Planck institute of Animal Behavior (Radolfzell, Germany) and cooperation partners. | CC_BY |

| | | | | | | |
|---|---|---|---|---|---|---|
| Movebank | 9651291 | Neophron percnopterus | Evan Buechley | Evan Buechley | Buechley ER, Şekercioğlu CH. 2019. Data from: Satellite tracking a wide-ranging endangered vulture species to target conservation actions in the Middle East and East Africa. Movebank Data Repository. <a href="https://www.doi.org/10.5441/001/1.385gk270" target="_blank">https://www.doi.org/10.5441/001/1.385gk270</a> <br><br> Phipps WL, López-López P, Buechley ER, Oppel S, Álvarez E, Arkumarev V, Bekmansurov R, Berger-Tal O, Bermejo A, Bounas A, Alanís IC, de la Puente J, Dobrev V, Duriez O, Efrat R, Fréchet G, García J, Galán M, García-Ripollés C, Gil A, Iglesias-Lebrija JJ, Jambas J, Karyakin IV, Kobierzycki E, Kret E, Loercher F, Monteiro A, Etxebarria JM, Nikolov SC, Pereira J, Peške L, Ponchon C, Realinho E, Saravia V, Şekercioğlu ÇH, Skartsi T, Tavares J, Teodósio J, Urios V, Vallverdú N. 2019. Spatial and temporal variability in migration of a soaring raptor across three continents. Front Ecol Evol. 7:323. https://doi.org/10.3389/fevo.2019.00323 <br><br> Buechley ER, McGrady MJ, Çoban E, Şekercioğlu ÇH. 2018. Satellite tracking a wide-ranging endangered vulture species to target conservation actions in the Middle East and East Africa. Biodivers Conserv. 27(9):2293-2310. https://doi.org/10.1007/s10531-018-1538-6 <br><br> Buechley ER, Oppel S, Beatty WS, Nikolov SC, Dobrev V, Arkumarev V, Saravia V, Bougain C, Bounas A, Kret E, Skartsi T, Aktay L, Ahababyan K, Frehner E, Şekercioğlu ÇH. 2018. Identifying critical migratory bottlenecks and high-use areas for an endangered migratory soaring bird across three continents. J Avian Biol. 49(7):e01629. https://doi.org/10.1111/jav.01629 | CC_BY |
| Movebank | 10006517 | Uria lomvia | Grant Gilchrist | Kyle Elliott | | CC_0 |
| Movebank | 10157679 | Ciconia ciconia | Azafzaf.H | Wolfgang Fiedler | Azafzaf, H., Feltrup-Azafzaf, C., Flack, A., Wikelski, M. & Fiedler, W. (year of access): GPS Tracking data from study "LifeTrack White Stork Tunisia" in www.movebank.org.. | CC_BY |

| | | | | | | |
|---|---|---|---|---|---|---|
| Movebank | 10204361 | Pandion haliaetus | Barb Jensen | Barb Jensen | | CC_0 |
| Movebank | 10236270 | Ciconia ciconia | Wolfgang Fiedler | Wolfgang Fiedler | Parts of this dataset are also available in the study "MPIO white stork lifetime tracking data (2013-2014)" and published as<br> Flack A, Fiedler W, Blas J, Pokrovski I, Mitropolsky B, Kaatz M, Aghababyan K, Khachatryan A, Fakriadis I, Makrigianni E, Jerzak L, Shamin M, Shamina C, Azafzaf H, Feltrup-Azafzaf C, Mokotjomela TM, Wikelski M. 2015. Data from: Costs of migratory decisions: a comparison across eight white stork populations. Movebank Data Repository. <a href="https://www.doi.org/10.5441/001/1.78152p3q" target="_blank">https://www.doi.org/10.5441/001/1.78152p3q</a> <br><br> Flack A, Fiedler W, Blas J, Pokrovski I, Kaatz M, Mitropolsky M, Aghababyan K, Fakriadis Y, Makrigianni E, Jerzak L, et al. 2016. Costs of migratory decisions: a comparison across eight white stork populations. Science Advances. 2(1): e1500931. https://doi.org/10.1126/sciadv.1500931 <br><br> Kays R, Davidson SC, Berger M, Bohrer G, Fiedler W, Flack A, Hirt J, Hahn C, Gauggel D, Russell B, et al. 2021. The Movebank system for studying global animal movement and demography. Methods Ecol Evol. https://doi.org/10.1111/2041-210X.13767 | CC_BY |
| Movebank | 10449318 | Ciconia ciconia,Homo sapiens | Martin Wikelski | Wolfgang Fiedler | Data from study "LifeTrack White Stork Loburg" in www.movebank.org by Max Planck Institute of Animal Behavior (Radolfzell, Germany), Hebrew University of Jerusalem (Israel), University of Potsdam (Germany) and Storchenhof Loburg (Germany) | CC_BY |

| | | | | | | |
|---|---|---|---|---|---|---|
| Movebank | 10449535 | Ciconia ciconia | Martin Wikelski | vasilis elias | Parts of this dataset are also available in the study "MPIO white stork lifetime tracking data (2013-2014)" and published as<br> Flack A, Fiedler W, Blas J, Pokrovski I, Mitropolsky B, Kaatz M, Aghababyan K, Khachatryan A, Fakriadis I, Makrigianni E, Jerzak L, Shamin M, Shamina C, Azafzaf H, Feltrup-Azafzaf C, Mokotjomela TM, Wikelski M. 2015. Data from: Costs of migratory decisions: a comparison across eight white stork populations. Movebank Data Repository. <a href="https://www.doi.org/10.5441/001/1.78152p3q" target="_blank">https://www.doi.org/10.5441/001/1.78152p3q</a> <br><br> Flack A, Fiedler W, Blas J, Pokrovski I, Kaatz M, Mitropolsky M, Aghababyan K, Fakriadis Y, Makrigianni E, Jerzak L, et al. 2016. Costs of migratory decisions: a comparison across eight white stork populations. Science Advances. 2(1): e1500931. https://doi.org/10.1126/sciadv.1500931<br><br>Kays R, Davidson SC, Berger M, Bohrer G, Fiedler W, Flack A, Hirt J, Hahn C, Gauggel D, Russell B, et al. 2021. The Movebank system for studying global animal movement and demography. Methods Ecol Evol. https://doi.org/10.1111/2041-210X.13767 | CC_BY |
| Movebank | 10449698 | Ciconia ciconia | Ran Nathan | Shay Rotics | Rotics et al. unpublished data | CC_BY |

| | | | | | | |
|---|---|---|---|---|---|---|
| Movebank | 10596067 | Ciconia ciconia | Martin Wikelski | Andrea Flack | Parts of this dataset are also available in the study "MPIO white stork lifetime tracking data (2013-2014)" and published as<br> Flack A, Fiedler W, Blas J, Pokrovski I, Mitropolsky B, Kaatz M, Aghababyan K, Khachatryan A, Fakriadis I, Makrigianni E, Jerzak L, Shamin M, Shamina C, Azafzaf H, Feltrup-Azafzaf C, Mokotjomela TM, Wikelski M. 2015. Data from: Costs of migratory decisions: a comparison across eight white stork populations. Movebank Data Repository. <a href="https://www.doi.org/10.5441/001/1.78152p3q" target="_blank">https://www.doi.org/10.5441/001/1.78152p3q</a> <br><br> Flack A, Fiedler W, Blas J, Pokrovski I, Kaatz M, Mitropolsky M, Aghababyan K, Fakriadis Y, Makrigianni E, Jerzak L, et al. 2016. Costs of migratory decisions: a comparison across eight white stork populations. Science Advances. 2(1): e1500931. https://doi.org/10.1126/sciadv.1500931<br><br>Kays R, Davidson SC, Berger M, Bohrer G, Fiedler W, Flack A, Hirt J, Hahn C, Gauggel D, Russell B, et al. 2021. The Movebank system for studying global animal movement and demography. Methods Ecol Evol. https://doi.org/10.1111/2041-210X.13767 | CC_BY |
| Movebank | 10722328 | Grus grus | Ramūnas Žydelis | Ramūnas Žydelis | Žydelis R, Desholm M, Månsson J, Nilsson L, Skov H. 2024. Data from: Study "GPS telemetry of Common Cranes, Sweden". Movebank Data Repository. <a href="https://doi.org/10.5441/001/1.597" target="_blank">https://doi.org/10.5441/001/1.597</a> <br><br> These data are described in<br> Yanco SW, Oliver RY, Iannarilli F, Carlson BS, Heine G, Mueller U, Richter N, Vorneweg B, Andryushchenko Y, Batbayar N, et al. 2024. Migratory birds modulate niche tradeoffs in rhythm with seasons and life history. Proc Natl Acad Sci USA. https://doi.org/10.1073/pnas.2316827121 | CC_BY |

| | | | | | | |
|---|---|---|---|---|---|---|
| Movebank | 10763606 | Ciconia ciconia | Martin Wikelski | Wolfgang Fiedler | Flack A, Fiedler W, Blas J, Pokrovsky I, Kaatz M, Mitropolsky M, Aghababyan K, Fakriadis I, Makrigianni E, Jerzak L, et al. 2016. Costs of migratory decisions: A comparison across eight white stork populations. Sci Adv. 2(1):e1500931. https://doi.org/10.1126/sciadv.1500931. | CC_BY |
| Movebank | 11223924 | Anser fabalis | Tomas Aarvak | Tomas Aarvak | | CC_BY |
| Movebank | 11948467 | Didelphis virginiana,Procyon lotor | Roland Kays | Roland Kays | | CC_0 |
| Movebank | 13978569 | Phalcoboenus australis | David Barber | David Barber | | CC_BY |
| Movebank | 14288429 | Equus hemionus | Petra Kaczensky | Petra Kaczensky | Kaczensky, P., Walzer, C. 2009. Gobi khulan GPS-ARGOS 2002-2008 tracking dataset. | CC_BY_NC |
| Movebank | 14671003 | Necrosyrtes monachus | David Barber | David Barber | | CC_BY |
| Movebank | 15661938 | Physeter macrocephalus | Daniel Palacios | Barb Lagerquist | Irvine LM, Follett TM, Winsor MH, Mate BR, Palacios DM. 2020. Data from: Study "Sperm whales Gulf of Mexico 2011-2013 - Argos data". Movebank Data Repository. doi: <a href="https://www.doi.org/10.5441/001/1.tj291471">10.5441/001/1.tj291471</a> <br><br> Irvine LM, Winsor MH, Follett TM, Mate BR, Palacios DM. 2020. An at-sea assessment of Argos location accuracy for three species of large whales, and the effect of deep-diving behavior on location error. Animal Biotelemetry. 8:20. doi: <a href="https://doi.org/10.1186/s40317-020-00207-x">10.1186/s40317-020-00207-x</a> | CC_BY |

| | | | | | | |
|---|---|---|---|---|---|---|
| Movebank | 16880941 | Cathartes aura | David Barber | David Barber | A more complete version of this dataset is stored in study "Vultures Acopian Center USA GPS".<br><br>Bildstein K, Barber D, Bechard MJ. 2014. Data from: Environmental drivers of variability in the movement ecology of turkey vultures (Cathartes aura) in North and South America. Movebank Data Repository. <a href="https://doi.org/10.5441/001/1.46ft1k05" target="_blank">https://doi.org/10.5441/001/1.46ft1k05</a> <br><br> Dodge S, Bohrer G, Bildstein K, Davidson SC, Weinzierl R, Mechard MJ, Barber D, Kays R, Brandes D, Han J. 2014. Environmental drivers of variability in the movement ecology of turkey vultures (Cathartes aura) in North and South America. Philos T Roy Soc B. 369(1643):20130195. https://doi.org/10.1098/rstb.2013.0195 | CC_0 |
| Movebank | 17469219 | Ardeidae ,Ardea herodias,Egretta thula,Ardea alba | John Brzorad | John Brzorad | | CC_BY_NC |
| Movebank | 19186107 | Fregata magnificens | Patrick Jodice | Patrick Jodice | Jodice, P.G.R., K. Meyer, S. Zaluski, and L. Soanes | CC_BY |

| | | | | | | |
|---|---|---|---|---|---|---|
| Movebank | 19411459 | Panthera onca | Ronaldo Morato | Ronaldo Morato | Morato RG, Kantek DLZ, Miyazaki S, Deluque T, de Paula RC. 2021. Data from: Jaguar movement database: a GPS-based movement dataset of an apex predator in the Neotropics. Movebank Data Repository. <a href="https://www.doi.org/10.5441/001/1.3c4fv0m4" target="_blank">https://www.doi.org/10.5441/001/1.3c4fv0m4</a> <br><br> These data are described in the following written publications:<br><br>Morato RG Thompson JJ, Paviolo A, De la Torre JA, Lima F, McBride Jr RT, De Paula RC, Cullen Jr L, Silveira L, Kantek DLZ, et al. 2018. Jaguar movement database: a GPS-based movement dataset of an apex predator in the Neotropics. Ecology. 99(7):1691. https://doi.org/10.1002/ecy.2379 <br><br> Morato RG, Connette GM, Stabach JA, De Paula RC, Ferraz KMPM, Kantek DLZ, Miyazaki SS, Pereira TDC, Silva LC, Paviolo A, et al. 2018. Resource selection in an apex predator and variation in response to local landscape characteristics. Biol Conserv. 228:233–40. https://doi.org/10.1016/j.biocon.2018.10.022 <br><br> Morato RG, Stabach JA, Fleming CH, Calabrese JM, De Paula RC, Ferraz KMPM, Kantek DLZ, Miyazaki SS, Pereira TDC, Araujo GR, et al. 2016. Space use and movement of a neotropical top predator: the endangered jaguar. PLoS ONE. 11(12):e0168176. https://doi.org/10.1371/journal.pone.0168176 <br><br> In addition, a version of these data are included as part of the following published datasets: <br><br> Morato RG, Thompson JJ, Paviolo A, de la Torre JA, Lima F, McBride Jr RT, Paula RC, Cullen Jr L, Silveira L, Kantek DLZ, et al. 2019. Data from: Jaguar Movement Database: a GPS-based movement dataset of an apex predator in the Neotropics. Dryad. https://doi.org/10.5061/dryad.2dh0223 <br><br> miltinhoastronauta. 2018. LEEClab/jaguar_movement: Jaguar Movement Database v1.0 released! (v1.0). Zenodo. [illegible] | CC_0 |

| | | | | | | |
|---|---|---|---|---|---|---|
| Movebank | 20202974 | Gyps himalayensis | Martin Wikelski | Sherub Sherub | Portions of these data are published as<br> Sherub S and Wikelski M. 2021. Data from: Longer days enable higher diurnal activity for migratory birds [Himalayan griffons]. Movebank Data Repository. <a href="https://www.doi.org/10.5441/001/1.4n2501f5" target="_blank">https://www.doi.org/10.5441/001/1.4n2501f5</a> <br><br> Sherub S, Wikelski M, Fiedler W, Davidson SC. 2016. Data from: Behavioural adaptations to flight into thin air. Movebank Data Repository. https://doi.org/10.5441/001/1.143v2p2k (study "High-altitude flights of Himalayan vultures (data from Sherub et al. 2016)") <br><br> Papers<br> Pokrovsky I, Kölzsch A, Sherub S, Fiedler W, Glazov P, Kulikova O, Wikelski M, Flack A. 2021. Longer days enable higher diurnal activity for migratory birds. J Anim Ecol. https://doi.org/10.1111/1365-2656.13484 <br><br> Sherub S, Bohrer G, Wikelski M, Weinzierl R. 2016. Behavioural adaptations to flight into thin air. Biology Letters. 12(10):20160432. https://doi.org/10.1098/rsbl.2016.0432 | CC_BY |
| Movebank | 20873986 | Haliaeetus leucocephalus | Roland Kays | Roland Kays | Roland Kays and Ted Simons | CC_0 |

| | | | | | | |
|---|---|---|---|---|---|---|
| Movebank | 21231406 | Ciconia ciconia | Martin Wikelski | Wolfgang Fiedler | Fiedler W, Flack A, Schäfle W, Keeves B, Quetting M, Eid B, Schmid H, Wikelski M. 2024. Data from: Study "LifeTrack White Stork SW Germany" (2013-2023). Movebank Data Repository. <a href="https://doi.org/10.5441/001/1.ck04mn78_2" target="_blank">https://doi.org/10.5441/001/1.ck04mn78_2</a> <br><br> These data are described in<br> Aikens EO, Nourani E, Fiedler W, Wikelski M, Flack A. 2024. Learning shapes the development of migratory behavior. Proc Natl Acad Sci USA. 121(12):e2306389121. https://doi.org/10.1073/pnas.2306389121<br><br>Kays R, Davidson SC, Berger M, Bohrer G, Fiedler W, Flack A, Hirt J, Hahn C, Gauggel D, Russell B, et al. 2022. The Movebank system for studying global animal movement and demography. Methods Ecol Evol. 13(2):419-431. https://doi.org/10.1111/2041-210X.13767 <br><br> Cheng Y, Fiedler W, Wikelski M, Flack A. 2019. "Closer-to-home" strategy benefits juvenile survival in a long-distance migratory bird. Ecology and Evolution. 9(16): 8945–8952. https://doi.org/10.1002/ece3.5395 <br><br> Flack A, Fiedler W, Blas J, Pokrovski I, Kaatz M, Mitropolsky M, Aghababyan K, Fakriadis Y, Makrigianni E, Jerzak L, et al. 2016. Costs of migratory decisions: a comparison across eight white stork populations. Science Advances. 2(1): e1500931. https://doi.org/10.1126/sciadv.1500931 <br><br> Weinzierl R, Bohrer G, Kranstauber B, Fiedler W, Wikelski M, Flack A. 2016. Wind estimation based on thermal soaring of birds. Ecology and Evolution. 6(24): 8706–8718. https://doi.org/10.1002/ece3.2585 <br><br> Parts of this dataset are also available in studies "Fall migration of white storks in 2014" (published as DOI 10.5441/001/1.bj96m274) and "MPIO white stork lifetime tracking data (2013-2014)" (published as DOI 10.5441/001/1.78152p3q). | CC_BY |

| | | | | | | |
|---|---|---|---|---|---|---|
| Movebank | 24442409 | Ciconia ciconia | Wolfgang Fiedler | Wolfgang Fiedler | Fiedler W, Leppelsack E, Leppelsack H, Stahl T, Wieding O, Wikelski M. 2024. Data from: Study "LifeTrack White Stork Bavaria" (2014-2023). Movebank Data Repository. <a href="https://doi.org/10.5441/001/1.v1cs4nn0_2" target="_blank">https://doi.org/10.5441/001/1.v1cs4nn0_2 </a> <br><br> Aikens EO, Nourani E, Fiedler W, Wikelski M, Flack A. 2024. Learning shapes the development of migratory behavior. Proc Natl Acad Sci USA. 121(12):e2306389121. https://doi.org/10.1073/pnas.2306389121<br><br> Brønnvik H, Nourani E, Fiedler W, Flack A. 2024. Experience reduces reliance on availability of conspecifics for route selection by a collectively migrating soaring bird. Curr Biol. 34(9):2030-2037.e3. https://doi.org/10.1016/j.cub.2024.03.052<br><br> Kays R, Davidson SC, Berger M, Bohrer G, Fiedler W, Flack A, Hirt J, Hahn C, Gauggel D, Russell B, et al. 2022. The Movebank system for studying global animal movement and demography. Methods Ecol Evol. 13(2):419-431. https://doi.org/10.1111/2041-210X.13767<br><br> Cheng Y, Fiedler W, Wikelski M, Flack A. 2019. "Closer-to-home" strategy benefits juvenile survival in a long-distance migratory bird. Ecology and Evolution. 9(16): 8945–8952. https://doi.org/10.1002/ece3.5395 | CC_BY |
| Movebank | 28691134 | Buteo platypterus | Laurie Goodrich | David Barber | contact PI | CC_BY |

| | | | | | | |
|---|---|---|---|---|---|---|
| Movebank | 29799425 | Branta leucopsis | Klaus-Michael Exo | Michael Exo | van der Jeugd H, Oosterbeek K, Ens BJ, Shamoun-Baranes J, Exo K. 2014. Data from: Forecasting spring from afar? Timing of migration and predictability of phenology along different migration routes of an avian herbivore [Barents Sea data]. Movebank Data Repository. <a href="https://www.doi.org/10.5441/001/1.ps244r11" target="_blank">https://www.doi.org/10.5441/001/1.ps244r11</a> <br><br> Shariati-Najafabadi M et al. 2016. Environmental parameters linked to the last migratory stage of barnacle geese en route to their breeding grounds. Animal Behavior. 118:81-95. https://doi.org/10.1016/j.anbehav.2016.05.018<br><br> Shariati-Najafabadi M et al. 2015. Satellite-versus temperature-derived green wave indices for predicting the timing of spring migration of avian herbivores. Ecological Indicators 58:322–331. https://doi.org/10.1016/j.ecolind.2015.06.005<br><br> Kölzsch A et al. 2015. Forecasting spring from afar? Timing of migration and predictability of phenology along different migration routes of an avian herbivore. J Anim Ecol. 84(1):272-283. https://doi.org/10.1111/1365-2656.12281<br><br> Shariatinajafabadi M et al. 2014. Migratory herbivorous waterfowl track satellite-derived green wave index. PLoS ONE 9(9):e108331. https://doi.org/10.1371/journal.pone.0108331<br><br> Ens BJ et al. 2008. Tracking of individual birds—report on WP 3230 (bird tracking sensor characterization) and WP 4130 (sensor adaptation and calibration for bird tracking system) of the FlySafe basic activities project. SOVON-onderzoeksrapport 2008/10, SOVON Vogelonderzoek Nederland, Beek-Ubbergen 78 p. | CC_0 |
| Movebank | 33622846 | Uria lomvia | Grant Gilchrist | Kyle Elliott | | CC_0 |

| | | | | | | |
|---|---|---|---|---|---|---|
| Movebank | 33643212 | Gyps fulvus,Gyps africanus,Torgos tracheliotus | Ran Nathan | Movement Ecology Lab at the Hebrew University of Jerusalem, Israel | Spiegel OM, Harel R, Centeno-Cuadros A, Hatzofe O, Getz WM, Nathan R. 2015. Data from: Moving beyond curve-fitting: using complementary data to assess alternative explanations for long movements of three vulture species. Movebank Data Repository. <a href="https://www.doi.org/10.5441/001/1.8c56f72s" target="_blank">https://www.doi.org/10.5441/001/1.8c56f72s</a> <br><br> Spiegel OM, Harel R, Centeno-Cuadros A, Hatzofe O, Getz WM, Nathan R. 2015. Moving beyond curve-fitting: using complementary data to assess alternative explanations for long movements of three vulture species. Am Nat. 185(2). https://www.doi.org/10.1086/679314 | CC_0 |
| Movebank | 37350671 | Buceros bicornis,Rhyticeros undulatus | Aparajita Datta and Rohit Naniwadekar | Rohit | Naniwadekar R, Rathore A, Shukla U, Chaplod S, Datta A. 2019. Data from: How far do Asian forest hornbills disperse seeds? Movebank Data Repository. <a href="https://www.doi.org/10.5441/001/1.14sm8k1d" target="_blank">https://www.doi.org/10.5441/001/1.14sm8k1d</a> <br><br> Naniwadekar R, Rathore A, Shukla U, Chaplod S, Datta A. 2019. How far do Asian forest hornbills disperse seeds? Acta Oecol. 101:103482. https://doi.org/10.1016/j.actao.2019.103482 | CC_0 |
| Movebank | 40906102 | Lemur catta | Teague O'Mara | Teague O'Mara | O'Mara. 2012. Development of feeding in ring-tailed lemurs. PhD Thesis. Arizona State University. | CC_0 |
| Movebank | 42451582 | Numenius americanus | Jay Carlisle | Jay Carlisle | | CC_BY_NC |
| Movebank | 47450376 | Grus antigone | Robert van Zalinge | Robert van Zalinge | | CC_BY |

| | | | | | | |
|---|---|---|---|---|---|---|
| Movebank | 53460105 | Papio anubis | Meg Crofoot | Meg Crofoot | Crofoot MC, Kays RW, Wikelski M. 2015. Data from: Shared decision-making drives collective movement in wild baboons. Movebank Data Repository. <a href="https://www.doi.org/10.5441/001/1.kn0816jn" target="_blank">https://www.doi.org/10.5441/001/1.kn0816jn</a> <br><br> Strandburg-Peshkin A, Farine DR, Couzin ID, Crofoot MC. 2015. Shared decision-making drives collective movement in wild baboons. Science. 348–6241: 1358–1361. https://doi.org/10.1126/science.aaa5099 <br><br> A larger dataset containing the data in this study is available as https://doi.org/10.5441/001/1.3q2131q5 and in the study "Collective movement in wild baboons". | CC_BY |
| Movebank | 67281010 | Mycteria americana | Roland Kays | Roland Kays | Schweitzer S, Bryan AL Jr, Brzorad J, Kays R. 2023. Data from: Study "NC Wood Stork Tracking". Movebank Data Repository. <a href="https://doi.org/10.5441/001/1.303" target="_blank">https://doi.org/10.5441/001/1.303</a> | CC_BY |
| Movebank | 69724677 | Branta bernicla,Branta leucopsis | Stefan Garthe | Stefan Garthe | | CC_0 |

| | | | | | | |
|---|---|---|---|---|---|---|
| Movebank | 74496970 | Ciconia ciconia | Andrea Flack | Andrea Flack | Flack A, Fiedler W, Blas J, Pokrovski I, Mitropolsky B, Kaatz M, Aghababyan K, Khachatryan A, Fakriadis I, Makrigianni E, et al. 2015. Data from: Costs of migratory decisions: a comparison across eight white stork populations. Movebank Data Repository. <a href="https://www.doi.org/10.5441/001/1.78152p3q" target="_blank">https://www.doi.org/10.5441/001/1.78152p3q</a> <br><br> Flack A, Fiedler W, Blas J, Pokrovski I, Kaatz M, Mitropolsky M, Aghababyan K, Fakriadis Y, Makrigianni E, Jerzak L, et al. 2016. Costs of migratory decisions: a comparison across eight white stork populations. Science Advances. 2(1): e1500931. https://doi.org/10.1126/sciadv.1500931<br><br>Kays R, Crofoot MC, Jetz W, Wikelski M. 2015. Terrestrial animal tracking as an eye on life and planet. Science. 348(6240):aaa2478. https://doi.org/10.1126/science.aaa2478 | CC_BY |

| | | | | | | |
|---|---|---|---|---|---|---|
| Movebank | 76367850 | Ciconia ciconia,Homo sapiens | Wolfgang Fiedler | Wolfgang Fiedler | Fiedler W, Hilsendegen C, Reis C, Lehmann J, Hilsendegen P, Schmid H, Wikelski M. 2024. Data from: Study "LifeTrack White Stork Rheinland-Pfalz" (2015-2023). Movebank Data Repository. <a href="https://doi.org/10.5441/001/1.4192t2j4_2" target="_blank">https://doi.org/10.5441/001/1.4192t2j4_2</a> <br><br>Aikens EO, Nourani E, Fiedler W, Wikelski M, Flack A. 2024. Learning shapes the development of migratory behavior. Proc Natl Acad Sci USA. 121(12):e2306389121. https://doi.org/10.1073/pnas.2306389121<br><br>Brønnvik H, Nourani E, Fiedler W, Flack A. 2024. Experience reduces reliance on availability of conspecifics for route selection by a collectively migrating soaring bird. Curr Biol. 34(9):2030-2037.e3. https://doi.org/10.1016/j.cub.2024.03.052<br><br>Kays R, Davidson SC, Berger M, Bohrer G, Fiedler W, Flack A, Hirt J, Hahn C, Gauggel D, Russell B, et al. 2022. The Movebank system for studying global animal movement and demography. Methods Ecol Evol. 13(2):419-431. https://doi.org/10.1111/2041-210X.13767<br><br>Cheng Y, Fiedler W, Wikelski M, Flack A. 2019. "Closer-to-home" strategy benefits juvenile survival in a long-distance migratory bird. Ecology and Evolution. 9(16): 8945–8952. https://doi.org/10.1002/ece3.5395 | CC_BY |
| Movebank | 79206236 | Ciconia ciconia | Martin Wikelski | Wolfgang Fiedler | Grasso R, Gagliardo A, Zafarana M, Müller I, Schmid H, Fiedler W, Wikelski M. 2021. Data from: Study "LifeTrack White Stork Sicily". Movebank Data Repository. <a href="https://www.doi.org/10.5441/001/1.4v8q16qf" target="_blank">https://www.doi.org/10.5441/001/1.4v8q16qf</a> <br><br> Kays R, Davidson SC, Berger M, Bohrer G, Fiedler W, Flack A, Hirt J, Hahn C, Gauggel D, Russell B, et al. 2021. The Movebank system for studying global animal movement and demography. Methods Ecol Evol. https://doi.org/10.1111/2041-210X.13767 | CC_BY |

| | | | | | | |
|---|---|---|---|---|---|---|
| Movebank | 92261778 | Cygnus cygnus | Dmitrijs Boiko | Wolfgang Fiedler | Dimitris Boiko, Wolfgang Fiedler & Martin Wikelski (2017): Project "Life Track Whooper Swan Latvia" at www.movebank.org | CC_BY |
| Movebank | 133992043 | Anser albifrons | Andrea Kölzsch | Andrea Kölzsch | Kölzsch A, Kruckenberg H, Glazov P, Müskens GJDM, Wikelski M. 2016. Data from: Towards a new understanding of migration timing: slower spring than autumn migration in geese reflects different decision rules for stopover use and departure. Movebank Data Repository. <a href="https://www.doi.org/10.5441/001/1.31c2v92f" target="_blank">https://www.doi.org/10.5441/001/1.31c2v92f</a> <br><br> Kölzsch A, Müskens GJDM, Kruckenberg H, Glazov P, Weinzierl R, Nolet BA, Wikelski M. 2016. Towards a new understanding of migration timing: slower spring than autumn migration in geese reflects different decision rules for stopover use and departure. Oikos. 125(10):1496–1507. https://doi.org/10.1111/oik.03121 | CC_BY |
| Movebank | 146932094 | Grus grus | Mindaugas Dagys & Ramūnas Žydelis | Ramūnas Žydelis | Dagys M, Žydelis R. 2024. Data from: Study "Common Crane Lithuania GPS, 2015-2016". Movebank Data Repository. <a href="https://doi.org/10.5441/001/1.604" target="_blank">https://doi.org/10.5441/001/1.604</a> <br><br> These data are described in<br> Yanco SW, Oliver RY, Iannarilli F, Carlson BS, Heine G, Mueller U, Richter N, Vorneweg B, Andryushchenko Y, Batbayar N, et al. 2024. Migratory birds modulate niche tradeoffs in rhythm with seasons and life history. Proc Natl Acad Sci USA. https://doi.org/10.1073/pnas.2316827121 | CC_BY |
| Movebank | 150597370 | Geronticus eremita | Didone Frigerio | Konrad Lorenz Research Station | Puehringer-Sturmayr V, Krejci J, Schuster R, et al. Space use and site fidelity in the endangered Northern Bald Ibis Geronticus eremita: Effects of age, season, and sex. Bird Conservation International. 2023;33:e10. doi:10.1017/S0959270922000053 | CC_BY_NC |
| Movebank | 154820583 | Gyps fulvus | Goran Susic | Wolfgang Fiedler | Life Track Griffon Vulture Croatia Study | CC_BY |
| Movebank | 172255794 | Steatornis caripensis | David A. Rodríguez. | Bernd Vorneweg | | CC_0 |

| | | | | | | |
|---|---|---|---|---|---|---|
| Movebank | 172972156 | Anthropoides virgo | Nyambayar Batbayar | Martin Wikelski | Batbayar N, Galtbalt B, Natsagdorj T, Sukhbaatar T, Wikelski M. 2024. Data from: Study "LifeTrack Mongolia Demoiselle cranes" Movebank Data Repository. <a href="https://doi.org/10.5441/001/1.599" target="_blank">https://doi.org/10.5441/001/1.599</a> <br><br> These data are described in<br> Yanco SW, Oliver RY, Iannarilli F, Carlson BS, Heine G, Mueller U, Richter N, Vorneweg B, Andryushchenko Y, Batbayar N, et al. 2024. Migratory birds modulate niche tradeoffs in rhythm with seasons and life history. Proc Natl Acad Sci USA. https://doi.org/10.1073/pnas.2316827121<br><br> Galtbalt B, Batbayar N, Sukhbaatar T, Vorneweg B, Heine G, Müller U, Wikelski M, Klaassen M. 2022. Differences in on-ground and aloft conditions explain seasonally different migration paths in demoiselle crane. Movement Ecology. https://doi.org/10.1186/s40462-022-00302-z <br><br> The processed data from Galtbalt et al. (2022) are published in Dryad at https://doi.org/10.5061/dryad.cnp5hqc1r | CC_BY_NC |

| | | | | | | |
|---|---|---|---|---|---|---|
| Movebank | 173641633 | Ciconia ciconia | Wolfgang Fiedler | Wolfgang Fiedler | Fiedler W, Niederer W, Schönenberger A, Flack A, Wikelski M. 2024. Data from: Study "LifeTrack White Stork Vorarlberg" (2016-2023). Movebank Data Repository. <a href="https://doi.org/10.5441/001/1.71r7pp6q_2" target="_blank">https://doi.org/10.5441/001/1.71r7pp6q_2</a> <br><br> These data are described in<br> Aikens EO, Nourani E, Fiedler W, Wikelski M, Flack A. 2024. Learning shapes the development of migratory behavior. Proc Natl Acad Sci USA. 121(12):e2306389121. https://doi.org/10.1073/pnas.2306389121<br><br>Brønnvik H, Nourani E, Fiedler W, Flack A. 2024. Experience reduces reliance on availability of conspecifics for route selection by a collectively migrating soaring bird. Curr Biol. 34(9):2030-2037.e3. https://doi.org/10.1016/j.cub.2024.03.052<br><br> Cheng Y, Fiedler W, Wikelski M, Flack A. 2019. "Closer-to-home" strategy benefits juvenile survival in a long-distance migratory bird. Ecology and Evolution. 9(16): 8945–8952. https://doi.org/10.1002/ece3.5395 | CC_BY |
| Movebank | 175720577 | Ciconia ciconia | Wolfgang Fiedler | Wolfgang Fiedler | Maxhuni Q, Gashi A, Hoxha L, Wolf G, Fiedler W. 2021. Data from: Study "LifeTrack White Stork Kosova". Movebank Data Repository. <a href="https://www.doi.org/10.5441/001/1.s367rd3k" target="_blank">https://www.doi.org/10.5441/001/1.s367rd3k</a> <br><br> Kays R, Davidson SC, Berger M, Bohrer G, Fiedler W, Flack A, Hirt J, Hahn C, Gauggel D, Russell B, et al. 2021. The Movebank system for studying global animal movement and demography. Methods Ecol Evol. https://doi.org/10.1111/2041-210X.13767 | CC_BY |

| | | | | | | |
|---|---|---|---|---|---|---|
| Movebank | 178979729 | Canis lupus | Dave Latham, Stan Boutin | Maria Cecilia Latham | Latham ADM, Boutin S. 2019. Data from: Wolf ecology and caribou-primary prey-wolf spatial relationships in low productivity peatland complexes in northeastern Alberta. Movebank Data Repository. <a href="https://www.doi.org/10.5441/001/1.7vr1k987" target="_blank">https://www.doi.org/10.5441/001/1.7vr1k987</a> <br><br> Latham AD. 2009. Wolf ecology and caribou-primary prey-wolf spatial relationships in low productivity peatland complexes in northeastern Alberta [dissertation]. [Alberta (CA)]: ProQuest Dissertations Publishing, University of Alberta. NR55419. 197 p. http://search.proquest.com/docview/305051214 <br><br> Latham AD, Latham MC, Boyce MS, Boutin S. 2011. Movement responses by wolves to industrial linear features and their effect on woodland caribou in northeastern Alberta. Ecol Appl. 21(8):2854–2865. https://doi.org/10.1890/11-0666.1 | CC_BY_NC |
| Movebank | 180290122 | Thalassoica antarctica | Descamps | Descamps | Descamps S, Tarroux A, Cherel Y, Delord K, Godø OR, Kato A, Krafft BA, Lorentsen S, Ropert-Coudert Y, Skaret G, Varpe Ø. 2016. Data from: At-sea distribution and prey selection of Antarctic petrels and commercial krill fisheries. Movebank Data Repository. <a href="https://www.doi.org/10.5441/001/1.q4gn4q56" target="_blank">https://www.doi.org/10.5441/001/1.q4gn4q56</a><br><br>These data are described in<br>Descamps S, Tarroux A, Cherel Y, Delord K, Godø OR, Kato A, Krafft BA, Lorentsen S-H, Ropert-Coudert Y, Skaret G, et al. 2016. At-sea distribution and prey selection of Antarctic petrels and commercial krill fisheries. PLoS ONE. 11(8):e0156968. https://doi.org/10.1371/journal.pone.0156968 | CC_BY |

| | | | | | | |
|---|---|---|---|---|---|---|
| Movebank | 182459847 | Cygnus cygnus | Martin Wikelski | Wolfgang Fiedler | Boiko D, Wikelski M, Fiedler W. 2019. Data from: Moulting sites of Latvian whooper swan Cygnus cygnus cygnets fitted with GPS-GSM transmitters. Movebank Data Repository. <a href="https://www.doi.org/10.5441/001/1.f89984gn" target="_blank">https://www.doi.org/10.5441/001/1.f89984gn</a> <br><br>Boiko D, Wikelski M. 2019. Moulting sites of Latvian whooper swan Cygnus cygnus cygnets fitted with GPS-GSM transmitters. Wildfowl. 5: 229-241. https://wildfowl.wwt.org.uk/index.php/wildfowl/article/view/2714<br><br> Boiko D, Wikelski M. 2018. Abstract: Moulting sites of Latvian Whooper Swan Cygnus cygnus cygnets tagged with transmitters in summer 2016. p. 40: http://conference.emu.ee/userfiles/swan2018/Luigekonverents148X210web.pdf | CC_BY |
| Movebank | 182746263 | Gyps himalayensis | Martin Wikelski | Sherub Sherub | Sherub S, Wikelski M, Fiedler W, Davidson SC. 2016. Data from: Behavioural adaptations to flight into thin air. Movebank Data Repository. <a href="https://www.doi.org/10.5441/001/1.143v2p2k" target="_blank">https://www.doi.org/10.5441/001/1.143v2p2k</a> <br><br> Sherub S, Bohrer G, Wikelski M, Weinzierl R. 2016. Behavioural adaptations to flight into thin air. Biology Letters. 12(10):20160432. https://doi.org/10.1098/rsbl.2016.0432 | CC_0 |
| Movebank | 183209639 | Larus fuscus | Stefan Garthe | Stefan Garthe | Garthe S, Schwemmer P, Paiva VH, Corman A-M, Fock HO, Voigt CC, Adler S (2016) Terrestrial and marine foraging strategies of an opportunistic seabird species breeding in the Wadden Sea. PLoS ONE 11(8): e0159630. doi:10.1371/journal.pone.0159630 | CC_BY_NC |
| Movebank | 185780950 | Geronticus eremita | Didone Frigerio | Konrad Lorenz Research Station | Puehringer-Sturmayr V, Krejci J, Schuster R, et al. Space use and site fidelity in the endangered Northern Bald Ibis Geronticus eremita: Effects of age, season, and sex. Bird Conservation International. 2023;33:e10. doi:10.1017/S0959270922000053 | CC_BY_NC |

| | | | | | | |
|---|---|---|---|---|---|---|
| Movebank | 186178781 | Pernis apivorus,Gyps fulvus | Daniel Schmidt-Rothmund | Wolfgang Fiedler | | CC_BY |
| Movebank | 193545363 | Canis latrans,Puma concolor | Julie Young, PhD | Peter Mahoney | Mahoney PJ, Ebinger M, Jaeger M, Shivik JA, Young JK. 2017. Data from: Uncovering behavioural states from animal activity and site fidelity patterns. Movebank Data Repository. <a href="https://www.doi.org/10.5441/001/1.7d8301h2" target="_blank">https://www.doi.org/10.5441/001/1.7d8301h2</a> <br><br>These data are described in<br>Mahoney PJ, Young JK. 2017. Uncovering behavioural states from animal activity and site fidelity patterns. Methods Ecol Evol. 8(2):174-183. https://doi.org/10.1111/2041-210X.12658 | CC_0 |
| Movebank | 208413731 | Connochaetes taurinus | Randall Boone | Jared Stabach | Stabach JA, Hughey L, Reid RS, Worden JS, Leimgruber P, Boone RB. 2020. Data from: Study "White-bearded wildebeest in Kenya". Movebank Data Repository. <a href="https://www.doi.org/10.5441/001/1.h0t27719" target="_blank">https://www.doi.org/10.5441/001/1.h0t27719</a> <br><br> Stabach JA, Hughey LF, Crego RD, Fleming CH, Hopcraft JGC, Leimgruber P, Morrison TA, Ogutu JO, Reid RS, Worden JS, Boone RB. 2022. Increasing anthropogenic disturbance restricts wildebeest movement across East African grazing systems. Front Ecol Evol. https://doi.org/10.3389/fevo.2022.846171 <br><br> Walton T, Boone R. 2017. Analyzing wildebeest behavior and effects of humans on activity in southwest Kenya. Poster [stored in this study; see Files] <br><br> Stabach JA. 2015. Movement, resource selection, and the physiological stress response of white-bearded wildebeest [dissertation]. [Fort Collins (CO, USA)]: Colorado State University. https://doi.org/10217/167207 <br><br> Boone RB, Reid RS, Lilieholm RJ, Worden JS, Ogutu JO. 2009. Wildebeest forage acquisition in fragmented landscapes under variable climates. National Science Foundation (NSF) DEB Grant 0919383. | CC_BY |

| | | | | | | |
|---|---|---|---|---|---|---|
| Movebank | 209824313 | Canis lupus | Mark Hebblewhite | Mark Hebblewhite | Hebblewhite M, Merrill EH (2008) Modelling wildlife-human relationships for social species with mixed-effects resource selection models. Journal of Applied Ecology 45:834-844. doi:10.1111/j.1365-2664.2008.01466.x <br> <br> Hebblewhite M, Merrill EH (2007) Multiscale wolf predation risk for elk: does migration reduce risk? Oecologia 152:377-387. doi:10.1007/s00442-007-0661-y | CC_BY |
| Movebank | 212096177 | Ciconia ciconia | Wolfgang Fiedler | Wolfgang Fiedler | Fiedler W, Flack A, Schmid A, Reinhard U, Wikelski M. 2024. Data from: Study "LifeTrack White Stork Oberschwaben" (2014-2023). Movebank Data Repository. <a href="https://doi.org/10.5441/001/1.c42j3js7_2" target="_blank">https://doi.org/10.5441/001/1.c42j3js7_2</a> <br><br> These data are described in<br> Aikens EO, Nourani E, Fiedler W, Wikelski M, Flack A. 2024. Learning shapes the development of migratory behavior. Proc Natl Acad Sci USA. 121(12):e2306389121. https://doi.org/10.1073/pnas.2306389121<br><br> Brønnvik H, Nourani E, Fiedler W, Flack A. 2024. Experience reduces reliance on availability of conspecifics for route selection by a collectively migrating soaring bird. Curr Biol. 34(9):2030-2037.e3. https://doi.org/10.1016/j.cub.2024.03.052<br><br> Kays R, Davidson SC, Berger M, Bohrer G, Fiedler W, Flack A, Hirt J, Hahn C, Gauggel D, Russell B, et al. 2021. The Movebank system for studying global animal movement and demography. Methods Ecol Evol. https://doi.org/10.1111/2041-210X.13767<br><br> Cheng Y, Fiedler W, Wikelski M, Flack A. 2019. "Closer-to-home" strategy benefits juvenile survival in a long-distance migratory bird. Ecology and Evolution. 9(16): 8945–8952. https://doi.org/10.1002/ece3.5395 | CC_BY |

| | | | | | | |
|---|---|---|---|---|---|---|
| Movebank | 216040785 | Rangifer tarandus | Dale R. Seip | Sarah Davidson | Seip DR, Price E. 2019. Data from: Science update for the South Peace Northern Caribou (Rangifer tarandus caribou pop. 15) in British Columbia. Movebank Data Repository. <a href="https://www.doi.org/10.5441/001/1.p5bn656k" target="_blank">https://www.doi.org/10.5441/001/1.p5bn656k</a> <br><br> Seip D, Jones E. 2017. Population status of Central Mountain caribou herds in British Columbia and response to recovery management actions. Ministry of Environment and Climate Change Strategy. 18 p. https://www2.gov.bc.ca/assets/gov/environment/plants-animals-and-ecosystems/wildlife-wildlife-habitat/regional-wildlife/northeast-region/caribou/central_mountains_caribou_population_report_2017.pdf <br><br> Johnson CJ, Ehlers LPW, Seip DR. 2015. Witnessing extinction: cumulative impacts across landscapes and the future loss of an evolutionarily significant unit of woodland caribou in Canada. Biol Conserv. 186:176-186. https://doi.org/10.1016/j.biocon.2015.03.012 <br><br> BC Ministry of Environment. 2014. Science update for the South Peace Northern caribou (Rangifer tarandus caribou pop. 15) in British Columbia. Victoria, BC. 43 p. https://www2.gov.bc.ca/assets/gov/environment/plants-animals-and-ecosystems/wildlife-wildlife-habitat/caribou/science_update_final_from_web_jan_2014.pdf <br><br> Jones ES, Gillingham MP, Seip DR, Heard DC. 2007. Comparison of seasonal habitat selection between threatened woodland caribou ecotypes in central British Columbia. Rangifer. 27(4):111-128. https://doi.org/10.7557/2.27.4.325 | CC_BY |

| | | | | | | |
|---|---|---|---|---|---|---|
| Movebank | 217784323 | Cathartes aura,Coragyps atratus | David Barber | David Barber | A more complete version of this dataset is stored in study "Vultures Acopian Center USA GPS".<br><br> Bildstein KL, Barber D, Bechard MJ, Graña Grilli M. 2016. Data from: Wing size but not wing shape is related to migratory behavior in a soaring bird. Movebank Data Repository. <a href="https://www.doi.org/10.5441/001/1.37r2b884" target="_blank">https://www.doi.org/10.5441/001/1.37r2b884</a> <br><br> Graña Grilli M, Lambertucci SA, Therrien J-F, Bildstein KL. 2017. Wing size but not wing shape is related to migratory behavior in a soaring bird. J Avian Biol. 48(5):669-678. https://doi.org/10.1111/jav.01220 <br><br> Dodge S, Bohrer G, Bildstein K, Davidson SC, Weinzierl R, Mechard MJ, Barber D, Kays R, Brandes D, Han J, et al. 2014. Environmental drivers of variability in the movement ecology of turkey vultures (Cathartes aura) in North and South America. Philos T Roy Soc B. 369(1643):20130195. https://doi.org/10.1098/rstb.2013.0195 | CC_BY |
| Movebank | 220078181 | Thalassarche chrysostoma | David Thompson | Rachael Orben | Thompson DR, Torres LG, Sagar PM, Kroeger CE, Orben RA. 2017. Data from: Classification of animal movement behavior through residence in space and time. Movebank Data Repository. <a href="https://www.doi.org/10.5441/001/1.694p666h" target="_blank">https://www.doi.org/10.5441/001/1.694p666h</a><br><br> Torres LG, Orben RA, Tolkova I, Thompson DR. 2017. Classification of animal movement behavior through residence in space and time. PLoS ONE. 12(1):e0168513. https://doi.org/10.1371/journal.pone.0168513 | CC_BY_NC |
| Movebank | 227843945 | Physeter macrocephalus | Daniel Palacios | Barb Lagerquist | Irvine L, Palacios DM, Urbán, Mate B. 2017. Sperm whale dive behavior characteristics derived from intermediate-duration archival tag data. Ecology and Evolution. 7:7822–7837. doi: <a href="https://doi.org/10.1002/ece3.3322">10.1002/ece3.3322</a> | CC_BY |

| | | | | | | |
|---|---|---|---|---|---|---|
| Movebank | 233141114 | Grus vipio | Nyambayar Batbayar | Nyambayar Batbayar | Batbayar N, Galtbalt B, Natsagdorj T, Sukhbaatar T, Wikelski M. 2024. Data from: Study "White-naped crane Mongolia WSCC" Movebank Data Repository. <a href="https://doi.org/10.5441/001/1.600" target="_blank">https://doi.org/10.5441/001/1.600</a> <br><br> These data are described in<br> Yanco SW, Oliver RY, Iannarilli F, Carlson BS, Heine G, Mueller U, Richter N, Vorneweg B, Andryushchenko Y, Batbayar N, et al. 2024. Migratory birds modulate niche tradeoffs in rhythm with seasons and life history. Proc Natl Acad Sci USA. https://doi.org/10.1073/pnas.2316827121 | CC_BY_NC |
| Movebank | 236953686 | Anas platyrhynchos | Wolfgang Fiedler | Wolfgang Fiedler | Duck data Max Planck Institute for Animal Behavior | CC_0 |
| Movebank | 249737372 | Uria lomvia | Kyle Elliott | Allison Patterson | | CC_0 |
| Movebank | 268904527 | Wallabia bicolor | Manuela Fischer | Manuela Fischer | Fischer M, Di Stefano J, Gras P, Kramer-Schadt S, Sutherland DR, Coulson G, Stillfried M (2020) Data from: Circadian rhythms enable efficient resource selection in a human-modified landscape. Movebank Data Repository. <a href="https://www.doi.org/10.5441/001/1.6ss053tn" target="_blank">https://www.doi.org/10.5441/001/1.6ss053tn</a> <br><br> Fischer M, Di Stefano J, Gras P, Kramer-Schadt S, Sutherland DR, Coulson G, Stillfried M (2019) Circadian rhythms enable efficient resource selection in a human-modified landscape. Ecology and Evolution 9(13): 7509–7527. https://doi.org/10.1002/ece3.5283 | CC_0 |
| Movebank | 270079388 | Uria lomvia | Grant Gilchrist | Allison Patterson | | CC_0 |
| Movebank | 277815715 | Ichthyaetus audouinii | Faggio | G Faggio | Recorbet, B., Faggio, G. & Boutten, W. (2017. Etude par géolocalisation de l’utilisation de l’espace par le goéland d’Audouin reproducteur des ZPS du Golfe d’Ajaccio. CEN-Corse | CC_BY_NC |
| Movebank | 295134472 | Equus quagga | Chamaille-Jammes | Chamaille-Jammes | | CC_BY_NC |
| Movebank | 312267867 | Larus marinus,Larus fuscus,Larus argentatus | Grigori Tertitski | Grigori Tertitski | | CC_0 |

| | | | | | | |
|---|---|---|---|---|---|---|
| Movebank | 326568799 | Gyps fulvus | Wolfgang Fiedler | Wolfgang Fiedler | Dieter Haas, Wolfgang Fiedler u.a. | CC_BY |
| Movebank | 358865092 | Uria lomvia | Kyle Elliott | Allison Patterson | | CC_0 |
| Movebank | 384868221 | Haliaeetus albicilla | Pawel; Mirski | Pawel; Mirski | Mirski P., unpubl. data. | CC_BY_NC |
| Movebank | 384882516 | Aquila pomarina | Pawel; Mirski | Pawel; Mirski | Mirski P., unpubl. | CC_BY_NC |
| Movebank | 439735878 | Hypsignathus monstrosus | Sarah Olson | Sarah Olson | Olson SH, Bounga G, Ondzie A, Bushmaker T, Seifert SN, Kuisma E, et al. (2019) Lek-associated movement of a putative Ebolavirus reservoir, the hammer-headed fruit bat (Hypsignathus monstrosus), in northern Republic of Congo. PLoS ONE 14(10): e0223139. https://doi.org/10.1371/journal.pone.0223139 | CC_BY |
| Movebank | 467107447 | Anser albifrons | Helmut Kruckenberg | Andrea Kölzsch | Kruckenberg H, Müskens GJDM, Ebbinge BS. 2018. Data from: A periodic Markov model to formalise animal migration on a network [white-fronted goose data]. Movebank Data Repository. <a href="https://www.doi.org/10.5441/001/1.kk38017f" target="_blank">https://www.doi.org/10.5441/001/1.kk38017f</a> <br><br> Kölzsch A, Kleyheeg E, Kruckenberg H, Kaatz M, Blasius B. 2018. A periodic Markov model to formalise animal migration on a network. R Soc Open Sci. 5(6):180438. https://doi.org/10.1098/rsos.180438 <br><br> Kölzsch A, Müskens GJDM, Kruckenberg H, Glazov P, Weinzierl R, Nolet BA, Wikelski M. 2016. Towards a new understanding of migration timing: slower spring than autumn migration in geese reflects different decision rules for stopover use and departure. Oikos. 125(10):1496-1507. https://doi.org/10.1111/oik.03121 <br><br> Van Wijk RE, Kölzsch A, Kruckenberg H, Ebbinge BS, Müskens GJDM, Nolet BA. 2012. Individually tracked geese follow peaks of temperature acceleration during spring migration. Oikos. 121(5):655-664. https://doi.org/10.1111/j.1600-0706.2011.20083.x | CC_BY |

| | | | | | | |
|---|---|---|---|---|---|---|
| Movebank | 468460067 | Nasua narica,Pecari tajacu,Ateles geoffroyi,Cebus capucinus,Potos flavus | Margaret Crofoot | Rasmus Havmoeller | Kays R, Hirsch B, Caillaud D, Mares R, Alavi S, Havmøller RW, Crofoot M. 2023. Data from: Multi-scale movement syndromes for comparative analyses of animal movement patterns. Movebank Data Repository. <a href="https://doi.org/10.5441/001/1.295" target="_blank">https://doi.org/10.5441/001/1.295</a> <br><br> These data are described in<br> Kays R, Hirsch B, Caillaud D, Mares R, Alavi S, Havmøller RW, Crofoot M. 2023. Multi-scale movement syndromes for comparative analyses of animal movement patterns. Move Ecol. 11:61. https://doi.org/10.1186/s40462-022-00365-y | CC_BY |
| Movebank | 492444603 | Canis lupus | Stan Boutin | Amanda Droghini | Bohm, H., A. Droghini, E. W. Neilson, and S. Boutin. 2018. Northern Alberta Grey Wolf, Study ID 492444603. Available on Movebank (movebank.org). Accessed on [access date]. | CC_0 |
| Movebank | 496130871 | Anthropoides virgo | Elena Ilyashenko | Ivan Pokrovskiy | Ilyashenko EI, Pokrovsky I, Ilyashenko VY, Mudrik EA, Korepov M, Mnatsekanov RA, Politov D, Fiedler W, Wikelski M. 2024. Data from: Study "1000 Cranes. Russia. Taman. Azov.". Movebank Data Repository. <a href="https://doi.org/10.5441/001/1.594" target="_blank">https://doi.org/10.5441/001/1.594</a> <br><br> These data are described in<br> Yanco SW, Oliver RY, Iannarilli F, Carlson BS, Heine G, Mueller U, Richter N, Vorneweg B, Andryushchenko Y, Batbayar N, et al. 2024. Migratory birds modulate niche tradeoffs in rhythm with seasons and life history. Proc Natl Acad Sci USA. https://doi.org/10.1073/pnas.2316827121 | CC_BY |

| | | | | | | |
|---|---|---|---|---|---|---|
| Movebank | 496136836 | Anthropoides virgo | Elena Ilyashenko | Ivan Pokrovskiy | Ilyashenko EI, Pokrovsky I, Ilyashenko VY, Mudrik EA, Korepov M, Politov D, Gugueva E, Fiedler W, Wikelski M. 2024. Data from: Study "1000 Cranes. Russia. Volga-Ural Interfluve.". Movebank Data Repository. <a href="https://doi.org/10.5441/001/1.595" target="_blank">https://doi.org/10.5441/001/1.595</a> <br><br> These data are described in<br> Yanco SW, Oliver RY, Iannarilli F, Carlson BS, Heine G, Mueller U, Richter N, Vorneweg B, Andryushchenko Y, Batbayar N, et al. 2024. Migratory birds modulate niche tradeoffs in rhythm with seasons and life history. Proc Natl Acad Sci USA. https://doi.org/10.1073/pnas.2316827121 | CC_BY |
| Movebank | 497087673 | Anthropoides virgo | Elena Ilyashenko | Ivan Pokrovskiy | Ilyashenko EI, Pokrovsky I, Mudrik EA, Politov D, Postelnykh K, Ilyashenko VY, Fiedler W, Wikelski M. 2024. Data from: Study "1000 Cranes. Russia. Altai.". Movebank Data Repository. <a href="https://doi.org/10.5441/001/1.596" target="_blank">https://doi.org/10.5441/001/1.596</a> <br><br> These data are described in<br> Yanco SW, Oliver RY, Iannarilli F, Carlson BS, Heine G, Mueller U, Richter N, Vorneweg B, Andryushchenko Y, Batbayar N, et al. 2024. Migratory birds modulate niche tradeoffs in rhythm with seasons and life history. Proc Natl Acad Sci USA. https://doi.org/10.1073/pnas.2316827121 | CC_BY |
| Movebank | 497091077 | Grus vipio,Anthropoides virgo | Elena Ilyashenko | Ivan Pokrovskiy | Ilyashenko EI, Pokrovsky I, Goroshko OA, Mudrik EA, Politov D, Ilyashenko VY, Fiedler W, Wikelski M. 2024. Data from: Study "1000 Cranes. Russia. Transbaikalia.". Movebank Data Repository. <a href="https://doi.org/10.5441/001/1.592" target="_blank">https://doi.org/10.5441/001/1.592</a> <br><br> These data are described in<br> Yanco SW, Oliver RY, Iannarilli F, Carlson BS, Heine G, Mueller U, Richter N, Vorneweg B, Andryushchenko Y, Batbayar N, et al. 2024. Migratory birds modulate niche tradeoffs in rhythm with seasons and life history. Proc Natl Acad Sci USA. https://doi.org/10.1073/pnas.2316827121 | CC_BY |

| | | | | | | |
|---|---|---|---|---|---|---|
| Movebank | 497138661 | Anthropoides virgo | Elena Ilyashenko | Ivan Pokrovskiy | Ilyashenko EI, Pokrovsky I, Gavrilov AE, Zaripova S, Fiedler W, Wikelski M. 2024. Data from: Study "1000 Cranes. Southern Kazakhstan.". Movebank Data Repository. <a href="https://doi.org/10.5441/001/1.591" target="_blank">https://doi.org/10.5441/001/1.591</a> <br><br> These data are described in<br> Yanco SW, Oliver RY, Iannarilli F, Carlson BS, Heine G, Mueller U, Richter N, Vorneweg B, Andryushchenko Y, Batbayar N, et al. 2024. Migratory birds modulate niche tradeoffs in rhythm with seasons and life history. Proc Natl Acad Sci USA. https://doi.org/10.1073/pnas.2316827121 | CC_BY |
| Movebank | 505156776 | Anser anser | Harald Grabenhofer | Harald Grabenhofer | Grabenhofer, H. et al, unpublished Data | CC_BY_NC |
| Movebank | 542653863 | Grus grus | Elena Ilyashenko | Ivan Pokrovskiy | Ilyashenko EI, Pokrovsky I, Ilyashenko VY, Korepov M, Fiedler W, Wikelski M. 2024. Data from: Study "1000 Cranes. Russia. Common Crane.". Movebank Data Repository. <a href="https://doi.org/10.5441/001/1.593" target="_blank">https://doi.org/10.5441/001/1.593</a> <br><br> These data are described in<br> Yanco SW, Oliver RY, Iannarilli F, Carlson BS, Heine G, Mueller U, Richter N, Vorneweg B, Andryushchenko Y, Batbayar N, et al. 2024. Migratory birds modulate niche tradeoffs in rhythm with seasons and life history. Proc Natl Acad Sci USA. https://doi.org/10.1073/pnas.2316827121 | CC_BY |
| Movebank | 560041066 | Ciconia ciconia | Ran Nathan | Shay Rotics | Rotics S, Kaatz M, Turjeman S, Zurell D, Wikelski M, Sapir N, Eggers U, Fiedler W, Jeltsch F, Nathan R. 2018. Data from: Early arrival at breeding grounds: causes, costs and a trade-off with overwintering latitude. Movebank Data Repository. <a href="https://www.doi.org/10.5441/001/1.v8d24552" target="_blank">https://www.doi.org/10.5441/001/1.v8d24552</a> <br><br> Rotics S, Kaatz M, Turjeman S, Zurell D, Wikelski M, Sapir N, Eggers U, Fiedler W, Jeltsch F, Nathan R. 2018. Early arrival at breeding grounds: causes, costs and a trade-off with overwintering latitude. J Anim Ecol. 87(6):1627-1638. https://doi.org/10.1111/1365-2656.12898 | CC_BY |

| | | | | | | |
|---|---|---|---|---|---|---|
| Movebank | 581719332 | Pelecanus occidentalis | Jordan Karubian | Jordan Karubian | | CC_BY_NC |
| Movebank | 604806671 | Circus aeruginosus | Anny Anselin | Tanja Milotic | Anselin A, Desmet P, Milotic T, Janssens K, T'Jollyn F, De Bruyn L, Bouten W (2019) MH_WATERLAND - Western marsh harriers (Circus aeruginosus, Accipitridae) breeding near the Belgium-Netherlands border. Dataset. <a target="_blank" href="https://doi.org/10.5281/zenodo.3532940">https://doi.org/10.5281/zenodo.3532940</a> | CC_0 |
| Movebank | 631036041 | Anthropoides virgo | Nyambayar Batbayar | Nyambayar Batbayar | Batbayar N, Galtbalt B, Natsagdorj T, Sukhbaatar T, Wikelski M. 2024. Data from: Study "1000 Cranes. Mongolia." Movebank Data Repository. <a href="https://doi.org/10.5441/001/1.598" target="_blank">https://doi.org/10.5441/001/1.598</a> <br><br> These data are described in<br> Yanco SW, Oliver RY, Iannarilli F, Carlson BS, Heine G, Mueller U, Richter N, Vorneweg B, Andryushchenko Y, Batbayar N, et al. 2024. Migratory birds modulate niche tradeoffs in rhythm with seasons and life history. Proc Natl Acad Sci USA. https://doi.org/10.1073/pnas.2316827121 | CC_BY_NC |
| Movebank | 657674643 | Anser albifrons | Andrea Kölzsch | Andrea Kölzsch | Kölzsch A, Müskens GJDM, Moonen S, Kruckenberg H, Glazov P, Wikelski M. 2019. Data from: Flyway connectivity and exchange primarily driven by moult migration in geese [North Sea population]. Movebank Data Repository. <a href="https://www.doi.org/10.5441/001/1.ct72m82n" target="_blank">https://www.doi.org/10.5441/001/1.ct72m82n</a> <br><br> Kölzsch A, Müskens GJDM, Szinai P, Moonen S, Glazov P, Kruckenberg H, Wikelski M, Nolet BA. 2019. Flyway connectivity and exchange primarily driven by moult migration in geese. Move Ecol. 7:3. https://doi.org/10.1186/s40462-019-0148-6 | CC_BY |

| | | | | | | |
|---|---|---|---|---|---|---|
| Movebank | 657965212 | Anser albifrons | Andrea Kölzsch | Andrea Kölzsch | Müskens GJDM, Szinai P, Sapi T, Kölzsch A, Wikelski M, Nolet BA. 2019. Data from: Flyway connectivity and exchange primarily driven by moult migration in geese [Pannonic population]. Movebank Data Repository. <a href="https://www.doi.org/10.5441/001/1.46b0mq21" target="_blank">https://www.doi.org/10.5441/001/1.46b0mq21</a> <br><br> Kölzsch A, Müskens GJDM, Szinai P, Moonen S, Glazov P, Kruckenberg H, Wikelski M, Nolet BA. 2019. Flyway connectivity and exchange primarily driven by moult migration in geese. Move Ecol. 7:3. https://doi.org/10.1186/s40462-019-0148-6 | CC_BY |
| Movebank | 672882373 | Milvus milvus | Patrick Scherler | Patrick Scherler | | CC_BY_NC |
| Movebank | 673728219 | Ovis dalli | James P Lawler | James P Lawler | Lawler JP, Griffith B, Johnson D, Burch J. 2005. The effects of military jet overflights on Dall's Sheep in Interior Alaska. Report to the Department of the Air Force. National Park Service, Alaska Region Natural Resource Technical Report NPS/AR/NRTR-2005-51, 181 p. | CC_BY |
| Movebank | 681056756 | Pteropus lylei | Julien Cappelle | Julien Cappelle | Choden K, Ravon S, Epstein JH, Hoem T, Furey N, Gely M, Jolivot A, Hul V, Neung C, Tran A, Cappelle J (2020) Data from: Pteropus lylei primarily forages in residential areas in Kandal, Cambodia. Movebank Data Repository. <a href="https://www.doi.org/10.5441/001/1.j25661td" target="_blank">https://www.doi.org/10.5441/001/1.j25661td</a> <br><br> Choden K, Ravon S, Epstein JH, Hoem T, Furey N, Gely M, Jolivot A, Hul V, Neung C, Tran A, Cappelle J (2019) Pteropus lylei primarily forages in residential areas in Kandal, Cambodia. Ecology and Evolution 9(7): 4181–4191. https://doi.org/10.1002/ece3.5046 | CC_0 |

| | | | | | | |
|---|---|---|---|---|---|---|
| Movebank | 736029750 | Loxodonta africana | Abi Tamim Vanak | Abi Tamim Vanak | Slotow R, Thaker M, Vanak AT (2019) Data from: Fine-scale tracking of ambient temperature and movement reveals shuttling behavior of elephants to water. Movebank Data Repository. <a href="https://www.doi.org/10.5441/001/1.403h24q5" target="_blank">https://www.doi.org/10.5441/001/1.403h24q5</a> <br><br> Thaker M, Gupte PR, Prins HHT, Slotow R, Vanak AT. 2019. Fine-scale tracking of ambient temperature and movement reveals shuttling behavior of elephants to water. Front Ecol Evol. 7:4. https://doi.org/10.3389/fevo.2019.00004 | CC_BY_NC |

| | | | | | | |
|---|---|---|---|---|---|---|
| Movebank | 897981076 | Cervus elaphus | Mark Hebblewhite | Mark Hebblewhite | Data for 2001-2020 are published as<br>Hebblewhite M, Merrill EH, Martin H, Berg JE, Bohm H, Eggeman SL. 2020. Data from: Study "Ya Ha Tinda elk project, Banff National Park, 2001-2020 (females)". Movebank Data Repository. <a href="https://www.doi.org/10.5441/001/1.5g4h5t6c" target="_blank">https://www.doi.org/10.5441/001/1.5g4h5t6c</a> <br><br> Tucker MA, Böhning-Gaese K, Fagan WF, Fryxell JM, Van Moorter B, Alberts SC, Ali AH, Allen AM, Attias N, Avgar T, et al. 2018. Moving in the Anthropocene: global reductions in terrestrial mammalian movements. Science 359(6374):466–469. https://www.doi.org/10.1126/science.aam9712 <br><br> Eggeman S, Hebblewhite M, Bohm H, Whittington J, Merrill E. 2016. Behavioral flexibility in migratory behavior in a long-lived large herbivore. J Anim Ecol 85(3):785-797. https://www.doi.org/10.1111/1365-2656.12495 <br><br> Hebblewhite M, Merrill EH, Morgantini LE, White CA, Allen JR, Bruns E, Thurston L, Hurd TE. 2006. Is the migratory behaviour of montane elk herds in peril? The case of Alberta’s Ya Ha Tinda elk herd. Wildlife Soc B 34(5):1280-1294. https://www.doi.org/10.2193/0091-7648(2006)34[1280:ITMBOM]2.0.CO;2 <br><br> Hebblewhite M, Merrill EH. 2009. Trade-offs between predation risk and forage differ between migrant strategies in a migratory ungulate. Ecology 90(12):3445-3454. https://www.doi.org/10.1890/08-2090.1 <br><br> Hebblewhite M, Merrill EH. 2008. Modelling wildlife-human relationships for social species with mixed-effects resource selection models. J Appl Ecol 45(3):834-844. https://www.doi.org/10.1111/j.1365-2664.2008.01466.x <br><br> Hebblewhite M, Merrill EH, McDermid G. 2008. A multi-scale test of the forage maturation hypothesis in a partially migratory ungulate population. Ecol Monogr 78(2):141-166. https://www.doi.org/10.1890/06-1708.1 <br><br> Hebblewhite M, Merrill EH. 2007. Multiscale wolf | CC_0 |
| Movebank | 909521569 | Pernis apivorus | Patrik Byholm | Elham Nourani | Nourani E, Vansteelant WMG, Byholm P, Safi K. 2020 Dynamics of the energy seascape can explain intra-specific variations in sea-crossing behaviour of soaring birds. Biol. Lett. 16: 20190797. http://dx.doi.org/10.1098/rsbl.2019.0797 | CC_BY_NC |

| | | | | | | |
|---|---|---|---|---|---|---|
| Movebank | 914913822 | Pelecanus onocrotalus | Ran Nathan | Ron Efrat | Efrat R, Hatzofe O, Nathan R. 2019. Data from: Landscape-dependent time versus energy optimisations in pelicans migrating through a large ecological barrier. Movebank Data Repository. <a href="https://www.doi.org/10.5441/001/1.hs79pk45" target="_blank">https://www.doi.org/10.5441/001/1.hs79pk45</a> <br><br> Efrat R, Hatzofe O, Nathan R. 2019. Landscape-dependent time versus energy optimisations in pelicans migrating through a large ecological barrier. Funct Ecol. 33(11):2161-2171. https://doi.org/10.1111/1365-2435.13426 | CC_0 |
| Movebank | 922263102 | Circus aeruginosus | Almut Schlaich & Tonio Schaub | Tanja Milotic | Koks B, Schlaich A, Schaub T, Klaassen R, Anselin A, Desmet P, Milotic T, Janssens K, Bouten W (2019) H_GRONINGEN - Western marsh harriers (Circus aeruginosus, Accipitridae) breeding in Groningen (the Netherlands). Dataset. <a target="_blank" href="https://doi.org/10.5281/zenodo.3552507">https://doi.org/10.5281/zenodo.3552507</a> | CC_0 |
| Movebank | 924418031 | Gallinago gallinago,Larus canus | Marcin | Marcin | www.interrex-tracking.com | CC_0 |

| | | | | | | |
|---|---|---|---|---|---|---|
| Movebank | 933711994 | Cervus elaphus | Mark S. Boyce | Sarah Davidson | Boyce MS and Ciuti S (2020) Data from: Human selection of elk behavioural traits in a landscape of fear. Movebank Data Repository. <a href="https://www.doi.org/10.5441/001/1.j484vk24" target="_blank">https://www.doi.org/10.5441/001/1.j484vk24</a> <br><br> Paton DG, Ciuti S, Quinn M, Boyce MS (2017) Hunting exacerbates the response to human disturbance in large herbivores while migrating through a road network. Ecosphere 8(6): e01841. https://doi.org/10.1002/ecs2.1841 <br><br> Prokopenko CM, Boyce MS, Avgar T (2017) Characterizing wildlife behavioural responses to roads using integrated step selection analysis. Journal of Applied Ecology 54: 470-479. https://doi.org/10.1111/1365-2664.12768 <br><br> Prokopenko CM, Boyce MS, Avgar T (2017) Extent-dependent habitat selection in a migratory large herbivore: road avoidance across scales. Landscape Ecology 32: 313-325. https://doi.org/10.1007/s10980-016-0451-1 <br><br> Roberts DR, Bahn V, Ciuti S, Boyce MS, Elith J, Guillera-Arroita G, Hauenstein S, Lahoz-Monfort JJ, Schöder B, Thuiller W, Warton DI, Wintle BA, Hartig F, Dormann CF (2017) Cross-validation strategies for data with temporal, spatial, hierarchical, or phylogenetic structure. Ecography 40: 913-929. https://doi.org/10.1111/ecog.02881 <br><br> Thurfjell H, Ciuti S, Boyce MS (2017) Learning from the mistakes of others: How female elk (Cervus elaphus) adjust behaviour with age to avoid hunters. PLoS ONE 12(6): e0178082. https://doi.org/10.1371/journal.pone.0178082 <br><br> Benz RA, Boyce MS, Thurfjell H, Paton DG, Musiani M, Dormann CF, et al. (2016) Dispersal ecology informs design of large-scale wildlife corridors. PLoS ONE 11(9): e0162989. https://doi.org/10.1371/journal.pone.0162989 <br><br> Ensing EP, Ciuti S, de Wijs FALM, Lentferink DH, ten Hoedt A, Boyce MS, Hut RA (2014) GPS based [illegible] | CC_BY |
| Movebank | 938783961 | Buteo buteo,Circus aeruginosus | Geert Spanoghe | Tanja Milotic | Spanoghe G, Desmet P, Milotic T, Janssens K, De Regge N, Vanoverbeke J, Bouten W (2019) MH_ANTWERPEN - Western marsh harriers (Circus aeruginosus, Accipitridae) breeding near Antwerp (Belgium). Dataset. <a target="_blank" href="https://doi.org/10.5281/zenodo.3550093">https://doi.org/10.5281/zenodo.3550093</a> | CC_0 |

| | | | | | | |
|---|---|---|---|---|---|---|
| Movebank | 943824007 | Balaenoptera physalus,Balaenoptera musculus | Daniel Palacios | Barb Lagerquist | Irvine LM, Palacios DM, Lagerquist BA, Mate BR, Follett TM. 2019. Data from: Scales of blue and fin whale feeding behavior off California, USA, with implications for prey patchiness. Movebank Data Repository. <a href="https://www.doi.org/10.5441/001/1.47h576f2" target="_blank">https://www.doi.org/10.5441/001/1.47h576f2</a> <br><br> Irvine LM, Palacios DM, Lagerquist BA, Mate BR. 2019. Scales of blue and fin whale feeding behavior off California, USA, with implications for prey patchiness. Front Ecol Evol. 7:338. https://doi.org/10.3389/fevo.2019.00338 <br><br> Irvine LM, Winsor MH, Follett TM, Mate BR, Palacios DM. 2020. An at-sea assessment of Argos location accuracy for three species of large whales, and the effect of deep-diving behavior on location error. Animal Biotelemetry. 8:20. https://doi.org/10.1186/s40317-020-00207-x | CC_BY |
| Movebank | 985143423 | Larus fuscus | Eric Stienen | Peter Desmet | Stienen EWM, Desmet P, Milotic T, Hernandez F, Deneudt K, Bouten W, Müller W, Matheve H, Lens L (2019) LBBG_ZEEBRUGGE - Lesser black-backed gulls (Larus fuscus, Laridae) breeding at the southern North Sea coast (Belgium and the Netherlands). Dataset. <a target="_blank" href="https://doi.org/10.5281/zenodo.3540799">https://doi.org/10.5281/zenodo.3540799</a> | CC_0 |
| Movebank | 986040562 | Larus argentatus | Eric Stienen | Peter Desmet | Stienen EWM, Desmet P, Milotic T, Hernandez F, Deneudt K, Matheve H, Bouten W (2019) HG_OOSTENDE - Herring gulls (Larus argentatus, Laridae) breeding at the southern North Sea coast (Belgium). Dataset. <a target="_blank" href="https://doi.org/10.5281/zenodo.3541811">https://doi.org/10.5281/zenodo.3541811</a> | CC_0 |

| | | | | | | |
|---|---|---|---|---|---|---|
| Movebank | 1007794443 | Larus marinus | Stefan Garthe | Stefan Garthe | Borrmann RM, Phillips RA, Clay TA, Garthe S. 2019. Data from: High foraging site fidelity and spatial segregation among individual great black-backed gulls. Movebank Data Repository. <a href="https://www.doi.org/10.5441/001/1.ht5jf68h" target="_blank">https://www.doi.org/10.5441/001/1.ht5jf68h</a> <br><br> Borrmann RM, Phillips RA, Clay TA, Garthe S. 2019. High foraging site fidelity and spatial segregation among individual great black-backed gulls. J Avian Biol. 50(12). https://doi.org/10.1111/jav.02156 | CC_BY_NC |
| Movebank | 1030734949 | Cygnus columbianus | Rascha Nuijten | Rascha Nuijten | Nuijten RJM, Gerrits T, de Vries PP, Müskens GJDM, Nolet BA. 2020. Data from: Less is more: on-board lossy compression of accelerometer data increases biologging capacity. Movebank Data Repository. <a href="https://www.doi.org/10.5441/001/1.8ms7mm80" target="_blank">https://www.doi.org/10.5441/001/1.8ms7mm80</a> <br><br> Nuijten RJM, Gerrits T, Shamoun-Baranes J, Nolet BA. 2020. Less is more: on-board lossy compression of accelerometer data increases biologging capacity. J Anim Ecol. 89(1):237-247. https://www.doi.org/10.1111/1365-2656.13164 | CC_0 |
| Movebank | 1049685237 | Anser albifrons | Andrea Kölzsch | Andrea Kölzsch | Kölzsch A, Müskens GJDM, Glazov P, Kruckenberg H, Wikelski M. 2020. Data from: Goose parents lead migration V. Movebank Data Repository. <a href="https://www.doi.org/10.5441/001/1.ms87s2m6" target="_blank">https://www.doi.org/10.5441/001/1.ms87s2m6</a><br><br>Kölzsch A, Flack A, Müskens GJDM, Kruckenberg H, Glazov P, Wikelski M. 2020. Goose parents lead migration V. J Avian Biol. 51(3). https://doi.org/10.1111/jav.02392 | CC_0 |

| | | | | | | |
|---|---|---|---|---|---|---|
| Movebank | 1071101052 | Larus argentatus | Robert Ronconi | Robert Ronconi | Ronconi RA, Shlepr KR. 2020. Data from: Study "Herring Gulls (Larus Argentatus); Ronconi; Kent Island, Canada". Movebank Data Repository. <a href="https://www.doi.org/10.5441/001/1.t5n4s456" target="_blank">https://www.doi.org/10.5441/001/1.t5n4s456</a> <br><br> Anderson CM, Gilchrist HG, Ronconi RA, Shlepr KR, Clark DE, Fifield DA, Robertson GJ, Mallory ML. 2020. Both short and long distance migrants use energy-minimizing strategies in North American herring gulls. Movement Ecology. 8:26. https://doi.org/10.1186/s40462-020-00207-9 | CC_0 |
| Movebank | 1071134107 | Larus argentatus | Robert Ronconi | Robert Ronconi | Ronconi RA, Shlepr KR. 2020. Data from: Study "Herring Gulls (Larus Argentatus); Ronconi; Brier Island, Canada". Movebank Data Repository. <a href="https://www.doi.org/10.5441/001/1.282vr7kd" target="_blank">https://www.doi.org/10.5441/001/1.282vr7kd</a> <br><br> Anderson CM, Gilchrist HG, Ronconi RA, Shlepr KR, Clark DE, Fifield DA, Robertson GJ, Mallory ML. 2020. Both short and long distance migrants use energy-minimizing strategies in North American herring gulls. Movement Ecology. 8:26. https://doi.org/10.1186/s40462-020-00207-9 | CC_0 |
| Movebank | 1073231887 | Buteo jamaicensis | Pete Bloom | Pete Bloom | Bloom PH (2015) Northward summer migration of red-tailed hawks fledged from southern latitudes. Journal of Raptor Research 49(1): 1-17. https://doi.org/10.3356/jrr-14-54.1 <br><br> Bloom PH (2011) Vagrancy, natal dispersal and migrations of red-tailed and red-shouldered hawks banded in the Pacific Flyway. Ph.D. dissertation, University of Idaho, Moscow, ID USA. | CC_BY |

| | | | | | | |
|---|---|---|---|---|---|---|
| Movebank | 1077432441 | Pteropus poliocephalus | Wayne Boardman | David Roshier | Roshier D, Boardman WSJ. 2021. Data from: Spring foraging movements of an urban population of grey-headed flying foxes (Pteropus poliocephalus). Movebank Data Repository. <a href="https://www.doi.org/10.5441/001/1.5bd6pq55" target="_blank">https://www.doi.org/10.5441/001/1.5bd6pq55</a> <br><br> Boardman WSJ, Roshier D, Reardon T, Burbidge K, McKeown A, Westcott DA, Caraguel CGB, Prowse TAA. 2021. Spring foraging movements of an urban population of grey-headed flying foxes (Pteropus poliocephalus). J Urban Ecol. 7(1):juaa034. https://doi.org/10.1093/jue/juaa034 | CC_0 |
| Movebank | 1080341217 | Larus argentatus | Dan Clark | Christine Anderson | Clark DE, Mackenzie SA, Koenen K, Whitney J, DeStefano S. 2020. Data from: Study "Herring Gulls (Larus Argentatus); Clark; Massachussets, United States". Movebank Data Repository. <a href="https://www.doi.org/10.5441/001/1.3th8b5q3" target="_blank">https://www.doi.org/10.5441/001/1.3th8b5q3</a> <br><br> Anderson CM, Gilchrist HG, Ronconi RA, Shlepr KR, Clark DE, Fifield DA, Robertson GJ, Mallory ML. 2020. Both short and long distance migrants use energy-minimizing strategies in North American herring gulls. Movement Ecology. 8:26. https://doi.org/10.1186/s40462-020-00207-9 | CC_0 |
| Movebank | 1080341737 | Larus argentatus | Robert Ronconi | Robert Ronconi | Ronconi RA, Taylor PD. 2020. Data from: Study "Herring Gulls (Larus Argentatus); Ronconi; Sable Island, Canada". Movebank Data Repository. <a href="https://www.doi.org/10.5441/001/1.3264ss3v" target="_blank">https://www.doi.org/10.5441/001/1.3264ss3v</a> <br><br> Anderson CM, Gilchrist HG, Ronconi RA, Shlepr KR, Clark DE, Fifield DA, Robertson GJ, Mallory ML. 2020. Both short and long distance migrants use energy-minimizing strategies in North American herring gulls. Movement Ecology. 8:26. https://doi.org/10.1186/s40462-020-00207-9 | CC_0 |

| | | | | | | |
|---|---|---|---|---|---|---|
| Movebank | 1087068449 | Tockus deckeni | Martin Wikelski | Katherine Mertes | Mertes K, Jetz W, Wikelski M (2020) Data from: Hierarchical multi-grain models improve descriptions of species’ environmental associations, distribution, and abundance. Movebank Data Repository. <a href="https://www.doi.org/10.5441/001/1.cp97k9j1" target="_blank">https://www.doi.org/10.5441/001/1.cp97k9j1</a> <br><br> Schwartz K, Jarzyna M, Jetz W (2020) Hierarchical multi-grain models improve descriptions of species’ environmental associations, distribution, and abundance. Ecological Applications. https://doi.org/10.1002/eap.2117 | CC_0 |
| Movebank | 1088836380 | Lynx rufus,Pekania pennanti | Scott LaPoint | Scott LaPoint | LaPoint et al. in progress | CC_BY |
| Movebank | 1091848505 | Larus ,Larus hyperboreus,Larus glaucescens,Larus smithsonianus | Andrew Ramey | Andrew Ramey | </br>Ramey, A.M., Ahlstrom, C.A., van Toor, M.L., Woksepp, H., Chandler, J.C., Reed, J.A., Reeves, A.B., Waldenström, J., Franklin, A.B., Bonnedahl, J., and Douglas, D.C. 2020, Tracking data for three large-bodied gull species and hybrids (Larus spp.) (ver 1.0, June 2020): U.S. Geological Survey data release, <a href="https://doi.org/10.5066/P9FZ4OJW" target="_blank">doi:10.5066/P9FZ4OJW</a></br> | CC_0 |
| Movebank | 1099562810 | Haematopus ostralegus | Geert Spanoghe | Tanja Milotic | Spanoghe G, Desmet P, Milotic T, Van Ryckegem G, Vanoverbeke J, Ens BJ, Bouten W (2020) O_WESTERSCHELDE - Eurasian oystercatchers (Haematopus ostralegus, Haematopodidae) breeding in East Flanders (Belgium). Dataset. <a target="_blank" href="https://doi.org/10.5281/zenodo.3734898">https://doi.org/10.5281/zenodo.3734898</a> | CC_0 |

| | | | | | | |
|---|---|---|---|---|---|---|
| Movebank | 1120749252 | Nasua narica,Pecari tajacu,Ateles geoffroyi,Cebus capucinus,Potos flavus | Meg Crofoot | Meg Crofoot | Kays R, Hirsch B, Caillaud D, Mares R, Alavi S, Havmøller RW, Crofoot M. 2023. Data from: Multi-scale movement syndromes for comparative analyses of animal movement patterns. Movebank Data Repository. <a href="https://doi.org/10.5441/001/1.295" target="_blank">https://doi.org/10.5441/001/1.295</a> <br><br> These data are described in<br> Kays R, Hirsch B, Caillaud D, Mares R, Alavi S, Havmøller RW, Crofoot M. 2023. Multi-scale movement syndromes for comparative analyses of animal movement patterns. Move Ecol. 11:61. https://doi.org/10.1186/s40462-022-00365-y | CC_BY |
| Movebank | 1123149708 | Pagophila eburnea | Morten Frederiksen | Morten Frederiksen | Frederiksen, M. (2019) GPS tracking of breeding ivory gulls in NE Greenland 2018-2019 | CC_BY_NC |
| Movebank | 1207235729 | Canis latrans | Nucharin Songsasen, PhD, DVM | Jared Stabach | | CC_0 |
| Movebank | 1208105916 | Larus cachinnans | INTERREX | INTERREX | | CC_0 |
| Movebank | 1229945587 | Grus grus | Petras Kurlavičius | Aleksej Perzhu | Unpublished & ongoing study by Lithuanian University of Educational Sciences (LEU) | CC_0 |

| | | | | | | |
|---|---|---|---|---|---|---|
| Movebank | 1241071371 | Vulpes lagopus | Dominique Berteaux | Dominique Berteaux | Clermont J., Grenier-Potvin A., Duchesne É., Couchoux C., Dulude-de Broin F., Beardsell A., Bêty J., Berteaux D. 2021. The predator activity landscape predicts the anti-predator behavior and distribution of prey in a tundra community. Ecosphere 12(12):e03858. <a href="https://doi.org/10.1002/ecs2.3858"target="_blank">https://doi.org/10.1002/ecs2.3858</a><br><br>Clermont J., Woodward-Gagné S., Berteaux D. 2021. Digging into the behaviour of an active hunting predator: arctic fox prey caching events revealed by accelerometry. Mov Ecol 9:58. <a href="https://doi.org/10.1186/s40462-021-00295-1" target="_blank"> https://doi.org/10.1186/s40462-021-00295-1</a><br><br>Poulin M-P, Clermont J, Berteaux D. 2021. Extensive daily movement rates measured in territorial arctic foxes. Ecol Evol. 11(6):2503-14. <a href="https://www.doi.org/10.1002/ece3.7165 " target="_blank">https://www.doi.org/10.1002/ece3.7165</a><br><br>Grenier-Potvin A, Clermont J, Gauthier G, Berteaux D. 2021. Prey and habitat distribution are not enough to explain predator habitat selection: addressing intraspecific interactions, behavioural state and time. Mov Ecol. 9:12. <a href="https://www.doi.org/10.1186/s40462-021-00250-0" target="_blank">https://www.doi.org/10.1186/s40462-021-00250-0</a><br><br> | CC_0 |
| Movebank | 1252230318 | Ciconia ciconia | Marcin Tobółka | Marcin Tobółka | Tobółka, M., Nowak, B., Białas, J.T., Jagiełło, Z., Zielińska, Z. 2020. The influence of parental experience and conditions during breeding and non-breeding period on reproductive success of long-lived birds, an example of the White Stork. | CC_0 |

| | | | | | | |
|---|---|---|---|---|---|---|
| Movebank | 1258895879 | Larus fuscus,Larus argentatus | Eric Stienen | Peter Desmet | Stienen EWM, Buijs R-J, de Visser J, Fijn R, Lilipaly S, Platteeuw M, Desmet P (2023) DELTATRACK - Herring gulls (Larus argentatus, Laridae) and lesser black-backed gulls (Larus fuscus, Laridae) breeding at Neeltje Jans (Netherlands). Dataset. <a target="_blank" href="https://doi.org/10.5281/zenodo.10209520">https://doi.org/10.5281/zenodo.10209520</a> | CC_0 |
| Movebank | 1259686571 | Larus fuscus,Larus argentatus | Eric Stienen | Peter Desmet | Stienen EWM, Müller W, Lens L, Milotic T, Desmet P (2021) LBBG_JUVENILE - Juvenile lesser black-backed gulls (Larus fuscus, Laridae) and herring gulls (Larus argentatus, Laridae) hatched in Zeebrugge (Belgium). Dataset. <a target="_blank" href="https://doi.org/10.5281/zenodo.5075868">https://doi.org/10.5281/zenodo.5075868</a> | CC_0 |
| Movebank | 1266784970 | Corvus corone | JIGUET Frédéric | Jiguet Frédéric | | CC_BY_NC |
| Movebank | 1278021460 | Buteo buteo,Circus cyaneus,Circus pygargus,Asio flammeus,Circus aeruginosus | Geert Spanoghe | Peter Desmet | Spanoghe G, Janssens K, Klaassen R, Schaub T, Milotic T, Desmet P (2021) BOP_RODENT - Rodent specialized birds of prey (Circus, Asio, Buteo) in Flanders (Belgium). Dataset. <a target="_blank" href="https://doi.org/10.5281/zenodo.5735405">https://doi.org/10.5281/zenodo.5735405</a> | CC_0 |
| Movebank | 1326534946 | Rissa ,Rissa tridactyla | Kyle Elliott | Kyle Elliott | | CC_BY |
| Movebank | 1375204238 | Pygoscelis adeliae | Yan Ropert-Coudert | Candice Michelot | Michelot et al (Accepted in Plos One) Adélie penguins foraging consistency and site fidelity are conditioned by breeding status and environmental conditions | CC_BY_NC |
| Movebank | 1393954358 | Cathartes aura | Martin Wikelski | Kamran Safi | | CC_BY_NC |

| | | | | | | |
|---|---|---|---|---|---|---|
| Movebank | 1395952585 | Numenius arquata | Philipp Schwemmer | Philipp Schwemmer | Schwemmer P, Garthe S. 2021. Data from: Migrating curlews on schedule: departure and arrival patterns of a long-distance migrant depend on time and breeding location rather than on wind conditions. Movebank Data Repository. <a href="https://www.doi.org/10.5441/001/1.715k46g2" target="_blank">https://www.doi.org/10.5441/001/1.715k46g2</a> <br><br> Schwemmer P, Mercker M, Vanselow KH, Bocher P, Garthe S. 2021. Migrating curlews on schedule: departure and arrival patterns of a long-distance migrant depend on time and breeding location rather than on wind conditions. Movement Ecology. https://doi.org/10.1186/s40462-021-00252-y | CC_0 |
| Movebank | 1404033416 | Anas rubripes | Chris Williams | Amanda Hoyt | | CC_0 |
| Movebank | 1406291704 | Panthera onca | Ronaldo Morato | Ronaldo Morato | Morato, RG; Tortato, F. | CC_BY |
| Movebank | 1410035327 | Ciconia ciconia | Ben Carlson | Ben Carlson | Carlson BS, Rotics S, Nathan R, Wikelski M, Jetz W. 2021. Individual environmental niches in mobile organisms. Nat Commun. 12:4572. doi: <a href="10.1038/s41467-021-24826-x">10.1038/s41467-021-24826-x</a> | CC_0 |
| Movebank | 1415844328 | Anser fabalis | Antti Piironen | Antti Piironen | Piironen A, Paasivaara A, Laaksonen T. 2021. Data from: Birds of three worlds: moult migration to high Arctic expands a boreal-temperate flyway to a third biome. Movebank Data Repository. <a href="https://www.doi.org/10.5441/001/1.22kk5126" target="_blank">https://www.doi.org/10.5441/001/1.22kk5126</a> <br><br> Piironen A, Paasivaara A, Laaksonen T. 2021. Birds of three worlds: moult migration to high Arctic expands a boreal-temperate flyway to a third biome. Movement Ecology. 9:47. https://www.doi.org/10.1186/s40462-021-00284-4 | CC_BY |

| | | | | | | |
|---|---|---|---|---|---|---|
| Movebank | 1426277950 | Tyto furcata | Matthew Johnson | Allison Huysman | Huysman AE, Castañeda XA, Johnson MD. 2021. Data from: Habitat selection by a predator of rodent pests is resilient to wildfire in a vineyard agroecosystem. Movebank Data Repository. <a href="https://www.doi.org/10.5441/001/1.82b5h1rk" target="_blank">https://www.doi.org/10.5441/001/1.82b5h1rk</a> <br><br> Huysman AE, Johnson MD. 2021. Habitat selection by a predator of rodent pests is resilient to wildfire in a vineyard agroecosystem. Ecol Evol. https://doi.org/10.1002/ece3.8416 <br><br> Castañeda XA, Huysman AE, Johnson MD. 2021. Barn Owls select uncultivated habitats for hunting in a winegrape growing region of California. Ornithol Appl. 123(1):duaa058. https://doi.org/10.1093/ornithapp/duaa058 | CC_BY |
| Movebank | 1429664821 | Connochaetes taurinus | Tom Morrison | Tom Morrison | Morrison TA, Link W, Newmark W, Foley C, Bolger DT. 2016. Tarangire revisited: population consequences of loss of migratory connectivity in a tropical ungulate. Biological Conservation. 197, 53-60. | CC_BY_NC |
| Movebank | 1445058753 | Larus fuscus | Marcin | Marcin | www.interrex-tracking.com | CC_0 |
| Movebank | 1448377103 | Mycteria americana | Mathieu Basille | Mathieu Basille | Basille M, Borkhataria RR, Bryan AL, Bucklin DN, Picardi S, Frederick PC. 2021. Data from: Study "Wood stork (Mycteria americana) Southeastern US 2004–2019". Movebank Data Repository. DOI: <a href="https://doi.org/10.5441/001/1.r0h6725k">10.5441/001/1.r0h6725k</a><br><br>Picardi S, Frederick PC, Borkhataria RR, Basille M. 2020. Partial migration in a subtropical wading bird in the Southeastern United States. Ecosphere. 11(2):e03054. DOI: <a href="https://doi.org/10.1002/ecs2.3054">10.1002/ecs2.3054</a><br><br>For the two particular individuals that moved to Mexico, see:<br>Picardi S, Borkhataria RR, Bryan AL, Jr, Frederick PC, Basille M. 2018. GPS telemetry reveals occasional dispersal of Wood Storks from the Southeastern US to Mexico. Caribbean Naturalist. Special Issue 2:23–29. | CC_0 |

| | | | | | | |
|---|---|---|---|---|---|---|
| Movebank | 1448409403 | Vanellus vanellus | Jelle Loonstra | Jelle Loonstra | | CC_BY_NC |
| Movebank | 1480583194 | Hydroprogne caspia | Patrik Byholm | Patrik Byholm | Byholm P, Beal M, Lötberg U, Åkesson S. 2022. Data from: Paternal transmission of migration knowledge in a long-distance bird migrant. Movebank Data Repository. <a href="https://www.doi.org/10.5441/001/1.352qf1cv" target="_blank">https://www.doi.org/10.5441/001/1.352qf1cv</a> <br><br> Byholm P, Beal M, Isaksson N, Lötberg U, Åkesson S. 2022. Paternal transmission of migration knowledge in a long-distance bird migrant. Nat Commun. https://doi.org/10.1038/s41467-022-29300-w | CC_BY |
| Movebank | 1531481854 | Phaethon aethereus,Sula dactylatra,Sula leucogaster,Fregata aquila,Onychoprion fuscatus | Diane Elizabeth Baum | Diane Elizabeth Baum | Ascension Island Government and MPIAB | CC_0 |
| Movebank | 1539583795 | Cuculus canorus | Dutch Montagu's Harrier Foundation | Dutch Montagu's Harrier Foundation | MPIAB and Dutch Montagu's Harrier Foundation | CC_0 |
| Movebank | 1542155599 | Gallinago gallinago,Cuculus canorus | Nyambayar Batbayar | Nyambayar Batbayar | Wildlife science and conservation center of Mongolia and MPIAB | CC_0 |
| Movebank | 1542192442 | Streptopelia turtur | JIGUET Frédéric | Jiguet Frédéric | MNHN and MPIAB | CC_BY_NC |
| Movebank | 1542206798 | Cuculus canorus | JIGUET Frédéric | Jiguet Frédéric | MNHN and MPIAB | CC_BY_NC |
| Movebank | 1543123177 | Onychoprion fuscatus | Christopher John Feare | Martin Wikelski | Seychelles Governmen, Chris Feare, Rachel Bristol, Christine Larose and MPIAB | CC_0 |
| Movebank | 1562253659 | Ciconia ciconia | Wolfgang Fiedler | Wolfgang Fiedler | Data from study "LifeTrack White Stork Sarralbe" in www.movebank.org by Max Planck Institute of Animal Behavior (Radolfzell, Germany) and Comune de Sarralbe | CC_0 |

| | | | | | | |
|---|---|---|---|---|---|---|
| Movebank | 1575973021 | Hydroprogne caspia | Susanne Akesson | Susanne Akesson | Åkesson S, Lötberg U, Rueda-Uribe C. 2022. Data from: Study "Tracking of Caspian Terns (Hydroprogne caspia) in the Swedish Baltic Sea 2017-2020". Movebank Data Repository. <a href="https://www.doi.org/10.5441/001/1.hg1v55ct" target="_blank">https://www.doi.org/10.5441/001/1.hg1v55ct</a><br><br> Rueda-Uribe C, Lötberg U, Ericsson M, Tesson SVM, Åkesson S. 2021. First tracking of declining Caspian Terns (Hydroprogne caspia) breeding in the Baltic Sea reveals high migratory dispersion and disjunct annual ranges as obstacles to effective conservation. J Avian Biol. 52(9). https://doi.org/10.1111/jav.02743 | CC_BY_NC |
| Movebank | 1605024900 | Chelydra serpentina,Branta canadensis,Anas platyrhynchos,Aix sponsa,Accipiter cooperii,Buteo jamaicensis,Buteo lineatus,Meleagris gallopavo,Bubo virginianus,Strix varia,Didelphis virginiana,Procyon lotor,Canis latrans | Stephen Blake | Stephen Blake | | CC_0 |
| Movebank | 1605797471 | Haematopus ostralegus | Bruno J. Ens | Peter Desmet | Dijkstra B, Dillerop R, Oosterbeek K, Bouten W, Desmet P, van der Kolk H, Ens BJ (2021) O_ASSEN - Eurasian oystercatchers (Haematopus ostralegus, Haematopodidae) breeding in Assen (the Netherlands). Dataset. <a target="_blank" href="https://doi.org/10.5281/zenodo.5653310">https://doi.org/10.5281/zenodo.5653310</a> | CC_0 |
| Movebank | 1605798640 | Haematopus ostralegus | Bruno J. Ens | Peter Desmet | Dokter AM, Oosterbeek K, Baptist M, Desmet P, van der Kolk H, Bouten W, Ens BJ (2021) O_BALGZAND - Eurasian oystercatchers (Haematopus ostralegus, Haematopodidae) wintering on Balgzand (the Netherlands). Dataset. <a target="_blank" href="https://doi.org/10.5281/zenodo.5653441">https://doi.org/10.5281/zenodo.5653441</a> | CC_0 |

| | | | | | | |
|---|---|---|---|---|---|---|
| Movebank | 1605799506 | Haematopus ostralegus | Bruno J. Ens | Peter Desmet | Oosterbeek K, Bom R, Shamoun-Baranes J, Desmet P, van der Kolk H, Bouten W, Ens BJ (2021) O_SCHIERMONNIKOOG - Eurasian oystercatchers (Haematopus ostralegus, Haematopodidae) breeding on Schiermonnikoog (the Netherlands). Dataset. <a target="_blank" href="https://doi.org/10.5281/zenodo.5653477">https://doi.org/10.5281/zenodo.5653477</a> | CC_0 |
| Movebank | 1605802367 | Haematopus ostralegus | Bruno J. Ens | Henk-Jan van der Kolk | van der Kolk H, Oosterbeek K, Jongejans E, Frauendorf M, Allen AM, Bouten W, Desmet P, de Kroon H, Ens BJ, van de Pol M (2021) O_VLIELAND - Eurasian oystercatchers (Haematopus ostralegus, Haematopodidae) breeding and wintering on Vlieland (the Netherlands). Dataset. <a target="_blank" href="https://doi.org/10.5281/zenodo.5653890">https://doi.org/10.5281/zenodo.5653890</a> | CC_0 |
| Movebank | 1605803389 | Haematopus ostralegus | Bruno J. Ens | Peter Desmet | Oosterbeek K, de Jong J, Desmet P, van der Kolk H, Bouten W, Ens BJ (2021) O_AMELAND - Eurasian oystercatchers (Haematopus ostralegus, Haematopodidae) breeding on Ameland (the Netherlands). Dataset. <a target="_blank" href="https://doi.org/10.5281/zenodo.5647596">https://doi.org/10.5281/zenodo.5647596</a> | CC_0 |
| Movebank | 1609400843 | Ichthyaetus melanocephalus | Eric Stienen | Peter Desmet | Stienen EWM, Desmet P, Milotic T, Spanoghe G, Janssens K (2022) MEDGULL_ANTWERPEN - Mediterranean gulls (Ichthyaetus melanocephalus, Laridae) breeding near Antwerp (Belgium). Dataset. <a target="_blank" href="https://doi.org/10.5281/zenodo.6599272">https://doi.org/10.5281/zenodo.6599272</a> | CC_0 |
| Movebank | 1623175929 | Rupicapra rupicapra | Martin Wikelski | Martin Wikelski | MPIAB and Miriam Wiesner | CC_0 |
| Movebank | 1671751878 | Struthio camelus | Dr Willem Burger | Willem Burger | | CC_BY_NC |
| Movebank | 1701931040 | Pernis apivorus | Wolfgang Fiedler | Wolfgang Fiedler | | CC_0 |
| Movebank | 1720694224 | Canis latrans,Puma concolor | Julie Young | Peter Mahoney | | CC_0 |

| | | | | | | |
|---|---|---|---|---|---|---|
| Movebank | 1724475305 | Larus michahellis | Marcin | Marcin | | CC_0 |
| Movebank | 1767692280 | Larus hyperboreus | Kyle Elliott | Allison Patterson | Baak JE, Patterson A, Gilchrist HG, Elliott KH. 2021. Data from: First evidence of diverging migration and overwintering strategies in glaucous gulls (Larus hyperboreus) from the Canadian Arctic. Movebank Data Repository. <a href="https://www.doi.org/10.5441/001/1.tj948m64" target="_blank">https://www.doi.org/10.5441/001/1.tj948m64</a> <br><br> Baak JE, Patterson A, Gilchrist HG, Elliott KH. 2021. First evidence of diverging migration and overwintering strategies in glaucous gulls (Larus hyperboreus) from the Canadian Arctic. Anim Migr. 8:98-109. https://doi.org/10.1515/ami-2020-0107 | CC_BY |
| Movebank | 1832666571 | Caracal caracal | Laurel Serieys | Laurel Serieys | Serieys LEK, Bishop JM. 2024. Data from: Study "Caracal movement ecology study in Cape Town, South Africa". Movebank Data Repository. <a href="https://doi.org/10.5441/001/1.317" target="_blank">https://doi.org/10.5441/001/1.317</a> <br><br> These data are described in<br> Serieys LEK, Bishop JM, Rogan MS, Smith JA, Suraci JP, O'Riain MJ, Wilmers CC. 2023. Anthropogenic activities and age class mediate carnivore habitat selection in a human-dominated landscape. iScience. 26(7):107050. https://doi.org/10.1016/j.isci.2023.107050 | CC_BY_NC |
| Movebank | 1841091905 | Numenius arquata | Geert Spanoghe | Peter Desmet | Spanoghe G, Janssens K, Nijs G, Milotic T, Desmet P (2021) CURLEW_VLAANDEREN - Eurasian curlews (Numenius arquata, Scolopacidae) breeding in Flanders (Belgium). Dataset. <a target="_blank" href="https://doi.org/10.5281/zenodo.5779130">https://doi.org/10.5281/zenodo.5779130</a> | CC_0 |
| Movebank | 1841261165 | Anas penelope | Jonas Waldenström | Marielle van Toor | van Toor ML, Kharitonov S, Švažas S, Dagys M, Kleyheeg E, Müskens G, Ottosson U, Žydelis R, Waldenström J. 2021. Migration distance affects how closely Eurasian wigeons follow spring phenology during migration. Mov Ecol. 9:61. doi: <a href="https://doi.org/10.1186/s40462-021-00296-0">10.1186/s40462-021-00296-0</a> | CC_BY |

| | | | | | | |
|---|---|---|---|---|---|---|
| Movebank | 1852945982 | Pteropus melanotus | Christopher | Christopher | Todd CM, Westcott DA, Martin JM, Rose K, McKeown A, Hall J, Welbergen JA. 2022. Data from: Study "Movements of the Christmas Island flying fox, Australia". Movebank Data Repository. <a href="https://www.doi.org/10.5441/001/1.mn019k4d" target="_blank">https://www.doi.org/10.5441/001/1.mn019k4d</a> <br><br> These data are described in<br> Todd CM, Westcott DA, Martin JM, Rose K, McKeown A, Hall J, Welbergen JA. 2022. Body-size dependent foraging strategies in the Christmas Island flying-fox: implications for seed and pollen dispersal within a threatened island ecosystem. Mov Ecol. 10:19. https://doi.org/10.1186/s40462-022-00315-8 | CC_0 |
| Movebank | 1879445455 | Chelonia mydas,Varanus salvator,Dendrocygna javanica,Anas platyrhynchos,Accipiter trivirgatus,Milvus migrans,Gyps himalayensis,Fulica atra,Numenius arquata,Tyto alba,Homo sapiens,Canis lupus,Bubo sumatranus,Nisaetus alboniger | Ratiwan Sitdhibutr | Martin Wikelski | | CC_BY |

| | | | | | | |
|---|---|---|---|---|---|---|
| Movebank | 1904388965 | Branta leucopsis,Anser albifrons,Anser anser | Bart Nolet | Nelleke Buitendijk | Buitendijk NH, de Jager M, Kruckenberg H, Kölzsch A, Moonen S, Müskens GJDM, Nolet BA. 2023. Data from: More grazing, more damage? Assessed damage to grassland relates non-linearly to goose grazing pressure. Movebank Data Repository. <a href="https://www.doi.org/10.5441/001/1.fk899541" target="_blank">https://www.doi.org/10.5441/001/1.fk899541</a> <br><br> These data are described in<br> Buitendijk N, de Jager M, Hornman M, Kruckenberg H, K ölzsch A, Moonen S, Nolet B. 2022. More grazing, more damage? Assessed yield loss on agricultural grassland relates non-linearly to goose grazing pressure. J Appl Ecol. 59(12): 2878-2889. https://doi.org/10.1111/1365-2664.14306 | CC_BY |
| Movebank | 1907973121 | Tapirus terrestris,calibration | Patricia Medici | Michael Noonan | Medici EP. 2023. Data from: Study "Lowland tapirs, Tapirus terrestris, in Southern Brazil". Movebank Data Repository. <a href="https://www.doi.org/10.5441/001/1.03ck4s52" target="_blank">https://www.doi.org/10.5441/001/1.03ck4s52</a> <br><br> These data are described in<br> Fleming CH, Deznabi I, Alavi S, Crofoot MC, Hirsch BT, Medici EP, Noonan MJ, Kays R, Fagan WF, Sheldon D, et al. 2022. Population-level inference for home-range areas. Methods Ecol Evol. 13(5):1027-1041. https://doi.org/10.1111/2041-210X.13815 <br><br> Medici EP, Mezzini S, Fleming CH, Calabrese JM, Noonan MJ. 2022. Movement ecology of vulnerable lowland tapirs between areas of varying human disturbance. Move Ecol. 10:14. https://doi.org/10.1186/s40462-022-00313-w | CC_BY_NC |
| Movebank | 1909487338 | Anas platyrhynchos | Martin Wikelski | Martin Wikelski | MPIAB and FAO | CC_0 |
| Movebank | 1941203363 | Aquila rapax,Trigonoceps occipitalis,Terathopius ecaudatus | Andre Botha | Martin Wikelski | Vultures for Africa and MPIAB | CC_0 |
| Movebank | 1958514115 | Aquila chrysaetos | Anders Tøttrup | Signe Agermose Mathiasen Andersen | | CC_BY_NC |

| | | | | | | |
|---|---|---|---|---|---|---|
| Movebank | 2105214573 | Branta canadensis | Manon Sorais | Manon Sorais | Sorais M., Patenaude-Monette M., Sharp C., Askren R., LaRocque A., Leblon B., and Giroux J.-F. | CC_0 |
| Movebank | 2190520177 | Larus michahellis | Marcin | Marcin | www.interrex-tracking.com | CC_0 |
| Movebank | 2272852276 | Larus fuscus | Marcin | Marcin | www.interrex-tracking.com | CC_BY_NC |
| Movebank | 2277774459 | Pelecanus occidentalis | Paul Leberg | Brock Geary | | CC_BY_NC |
| Movebank | 2290252202 | Branta leucopsis,Anser albifrons,Anser fabalis,Anser brachyrhynchus | Bart Nolet | Andrea Kölzsch | Kölzsch A, Lameris TK, Müskens GJDM, Schreven KHT, Buitendijk NH, Kruckenberg H, Moonen S, Heinicke T, Cao L, Madsen J, Wikelski M, Nolet BA. 2022. Data from: Wild goose chase: geese flee high and far, and with aftereffects from New Year’s fireworks. Movebank Data Repository. <a href="https://www.doi.org/10.5441/001/1.g51fs0jv" target="_blank">https://www.doi.org/10.5441/001/1.g51fs0jv</a> <br><br> These data are described in<br> Kölzsch A, Lameris TK, Müskens GJDM, Schreven KHT, Buitendijk NH, Kruckenberg H, Moonen S, Heinicke T, Cao L, Madsen J, et al. 2022. Wild goose chase: geese flee high and far, and with aftereffects from New Year’s fireworks. Conserv Lett. e12927. https://doi.org/10.1111/conl.12927 | CC_BY |
| Movebank | 2298738353 | Larus fuscus | Eric Stienen | Peter Desmet | Stienen EWM, Müller W, Lens L, Milotic T, Desmet P (2023) LBBG_ADULT - Lesser black-backed gulls (Larus fuscus, Laridae) breeding in Belgium. Dataset. <a target="_blank" href="https://doi.org/10.5281/zenodo.10055493">https://doi.org/10.5281/zenodo.10055493</a> | CC_0 |
| Movebank | 2313947453 | Platalea leucorodia | Geert Spanoghe | Peter Desmet | Spanoghe G, Janssens K, Milotic T, Desmet P (2023) SPOONBILL_VLAANDEREN - Eurasian spoonbills (Platalea leucorodia, Threskiornithidae) in Flanders (Belgium). Dataset. <a target="_blank" href="https://doi.org/10.5281/zenodo.10055132">https://doi.org/10.5281/zenodo.10055132</a> | CC_0 |

| | | | | | | |
|---|---|---|---|---|---|---|
| Movebank | 2379775078 | Hypsignathus monstrosus | Elodie Schloesing | Elodie Schloesing | Schloesing E, Caron A, Chambon R, Courbin N, Labadie M, Nina R, Mouti Mdadinga F, Ngoubili W, Sandial D, N'Kaya-Tobi , Bourgarel M, De Nys HM, Cappelle J. 2024. Data from: Foraging and mating behaviors of Hypsignathus monstrosus at the bat-human interface in central African rainforest. Movebank Data Repository. <a href="https://doi.org/10.5441/001/1.278" target="_blank">https://doi.org/10.5441/001/1.278</a> <br><br> These data are described in<br> Schloesing E, Caron A, Chambon R, Courbin N, Labadie M, Nina R, Mouiti Mbadinga F, Ngoubili W, Sandiala D, N'Kaya Tobi, Bourgarel M, De Nys HM, Cappelle J. 2023. Foraging and mating behaviors of Hypsignathus monstrosus at the bat-human interface in central African rainforest. Ecol Evol. 13(7):e10240. https://doi.org/10.1002/ece3.10240 | CC_0 |
| Movebank | 2398637362 | Falco naumanni | Javier Bustamante | Javier Bustamante | | CC_BY_NC |
| Movebank | 2425475445 | Gyps bengalensis,Gyps himalayensis | A B M Sarowar ALAM | A B M Sarowar ALAM | | CC_0 |
| Movebank | 2526306516 | Mergus merganser | Anthony Wetherhill | Anthony Wetherhill | | CC_BY_NC |
| Movebank | 2608802883 | Rangifer tarandus | Leif Egil Loe | Samantha P. H. Dwinnell | Loe, L. E., B. B. Hansen, A. Stien, S. D. Albon, R. Bischof, A. Carlsson, R. J. Irvine, M. Meland, I. M. Rivrud, E. Ropstad, V. Veiberg, and A. Mysterud. 2016. Behavioral buffering of extreme weather events in a high- Arctic herbivore. Ecosphere 7(6):e01374. 10.1002/ecs2.1374 | CC_BY_NC |

| | | | | | | |
|---|---|---|---|---|---|---|
| Movebank | 2630711281 | Morus bassanus | William A. Montevecchi | Kyle d'Entremont | d'Entremont KJN, Davoren GK, Montevecchi WA. 2023. Data from: Study "Northern Gannet Breeding Season GPS Data from Cape St. Mary's, NL, Canada: 2019 to 2022". Movebank Data Repository. <a href="https://www.doi.org/10.5441/001/1.5km7v2s3" target="_blank">https://www.doi.org/10.5441/001/1.5km7v2s3</a> <br><br> These data are described in<br> d'Entremont KJN, Davoren GK, Walsh CJ, Wilhelm SI, Montevecchi WA. 2022. Intra- and inter-annual shifts in foraging tactics by parental northern gannets Morus bassanus indicate changing prey fields. Mar Ecol Prog Ser. 698:155-170. https://www.doi.org/10.3354/meps14164 | CC_BY_NC |
| Movebank | 2636372210 | Lynx rufus,Canis latrans | Laura Prugh | Laura Prugh | Prugh LR. 2023. Data from: Study "GPS tracking of bobcats and coyotes in northern Washington". Movebank Data Repository. <a href="https://www.doi.org/10.5441/001/1.gm93267b" target="_blank">https://www.doi.org/10.5441/001/1.gm93267b</a> <br><br> These data are described in<br> Prugh LR, Cunningham CX, Windell RM, Kertson BN, Ganz TR, Walker SL, Wirsing AJ. 2023. Fear of large carnivores amplifies human-caused mortality for mesopredators. Science. 380(6646):754-758. https://doi.org/10.1126/science.adf2472 <br><br> Bassing SB, DeVivo M, Ganz TR, Kertson BN, Prugh LR, Roussin T, Satterfield L, Windell RM, Wirsing AJ, Gardner B. 2023. Are we telling the same story? Comparing inferences made from camera trap and telemetry data for wildlife monitoring. Ecol Appl. 33(1):e2745. https://doi.org/10.1002/eap.2745 | CC_BY |

| | | | | | | |
|---|---|---|---|---|---|---|
| Movebank | 2658117564 | Morus bassanus | Jana W E Jeglinski | Jana W E Jeglinski | Jeglinski JWE, Matthiopoulos J, Votier SC, Lane JV. 2024. Data from: Strong breeding colony fidelity in northern gannets following high pathogenicity avian influenza HPAIV outbreak. Movebank Data Repository. <a href="https://doi.org/10.5441/001/1.608" target="_blank">https://doi.org/10.5441/001/1.608</a> <br><br> These data are described in<br>Grémillet D, Ponchon A, Provost P, Gamble A, Abed-Zahar M, Bernard A, Courbin N, Delavaud G, Deniau A, Fort J, et al. 2023. Strong breeding colony fidelity in northern gannets following high pathogenicity avian influenza HPAIV outbreak. Biol Conserv. 286:110269. https://doi.org/10.1016/j.biocon.2023.110269<br><br> Jeglinski JWE, Lane JV, Votier SC, Furness RW, Hamer KC, McCafferty DJ, Nager RG, Sheddan M, Wanless S, Matthiopoulos J. 2024. HPAIV outbreak triggers short-term colony connectivity in a seabird metapopulation. Sci Rep. 14:3126. https://doi.org/10.1038/s41598-024-53550-x | CC_BY |
| Movebank | 2658220054 | Morus bassanus | Jana W E Jeglinski | Jana W E Jeglinski | Jeglinski JWE, Lane JV, Votier SC, Furness RW, Hamer KC, McCafferty DJ, Nager R, Sheddan M, Wanless S, Matthiopoulos J. 2024. Data from: HPAIV outbreak triggers short-term colony connectivity in a seabird metapopulation. Movebank Data Repository. <a href="https://doi.org/10.5441/001/1.607" target="_blank">https://doi.org/10.5441/001/1.607</a> <br><br> These data are described in<br> Jeglinski JWE, Lane JV, Votier SC, Furness RW, Hamer KC, McCafferty DJ, Nager RG, Sheddan M, Wanless S, Matthiopoulos J. 2024. HPAIV outbreak triggers short-term colony connectivity in a seabird metapopulation. Sci Rep. 14:3126. https://doi.org/10.1038/s41598-024-53550-x | CC_BY_NC |
| Movebank | 2830157729 | Nycticorax nycticorax | Mike Ward | Sarah Slayton | | CC_0 |
| Movebank | 2915700754 | Bubulcus ibis | Benjamin Vollot | Benjamin Vollot | | CC_0 |

| | | | | | | |
|---|---|---|---|---|---|---|
| Movebank | 2944153255 | Larus smithsonianus | Mark Mallory | Sarah Gutowsky | Mallory ML, Craik S, Allard KA, Gutowsky S. 2023. Data from: Study "American Herring Gulls - GPS - Lobster Bay, Southwest Nova Scotia, Canada". Movebank Data Repository. <a href="https://www.doi.org/10.5441/001/1.292" target="_blank">https://www.doi.org/10.5441/001/1.292</a> <br><br> These data are described in<br> Gutowsky S, Baak JE, Craik S, Mallory M, Knutson N, d'Entremont A, Allard K. 2023. Seasonal and circadian patterns of herring gull (Larus smithsoniansus) movements reveal temporal shifts in industry and coastal island interaction. Ecol Solut Evid. 4(3):e12274. https://doi.org/10.1002/2688-8319.12274 | CC_0 |
| Movebank | 2961927604 | Falco naumanni | Javier Bustamante | Javier Bustamante | Lesser kestrel movements, EBD-CSIC, projects MERCURIO-SUMHAL | CC_BY_NC |
| Movebank | 2970193504 | Falco tinnunculus | Javier Bustamante | Javier Bustamante | Common kestrel (Falco tinnunculus) movements, EBD-CSIC, projects MERCURIO & SUMHAL | CC_BY_NC |
| Movebank | 2976678046 | Odocoileus virginianus | Blaise Ashley Newman | Blaise Ashley Newman | Newman BA, Dyal JR, Miller KV, Cherry MJ, D'Angelo GJ. 2023. Data from: Influence of visual perception on movement decisions by an ungulate prey species. Movebank Data Repository. <a href="https://www.doi.org/10.5441/001/1.293" target="_blank">https://www.doi.org/10.5441/001/1.293</a> <br><br> These data are described in<br> Newman BA, Dyal JR, Miller KV, Cherry MJ, D'Angelo GJ. 2023. Influence of visual perception on movement decisions by an ungulate prey species. Biol Open. 12(10):bio059932. https://doi.org/10.1242/bio.059932 <br><br> Dyal JR, Miller KV, Cherry MJ, D'Angelo GJ. 2022. White-tailed deer movement in response to helicopter surveys. Wildlife Soc Bull. 46(5):e1383. https://doi.org/10.1002/wsb.1383 | CC_BY |

| | | | | | | |
|---|---|---|---|---|---|---|
| Movebank | 2984378217 | Anthropoides virgo | Yuriy | Ivan Pokrovskiy | Andryushchenko Y, Pokrovsky I, Bernd V, Fiedler W, Wikelski M. 2024. Data from: Study "1000 Cranes. Ukraine.". Movebank Data Repository. <a href="https://doi.org/10.5441/001/1.590" target="_blank">https://doi.org/10.5441/001/1.590</a> <br><br> These data are described in<br> Yanco SW, Oliver RY, Iannarilli F, Carlson BS, Heine G, Mueller U, Richter N, Vorneweg B, Andryushchenko Y, Batbayar N, et al. 2024. Migratory birds modulate niche tradeoffs in rhythm with seasons and life history. Proc Natl Acad Sci USA. https://doi.org/10.1073/pnas.2316827121 | CC_BY |
| Movebank | 3094468724 | Neophron percnopterus | Ron Efrat | Ron Efrat | Efrat R, Hatzofe O, Mueller T, Sapir N, Berger-Tal O. 2023. Data from: Early life and acquired experiences interact in shaping migratory and flight behaviors. Movebank Data Repository. <a href="https://www.doi.org/10.5441/001/1.298" target="_blank">https://www.doi.org/10.5441/001/1.298</a > <br><br> These data are described in<br> Efrat R, Hatzofe O, Mueller T, Sapir N, Berger-Tal O. 2023. Early life and acquired experiences interact in shaping migratory and flight behaviors. Curr Biol. https://doi.org/10.1016/j.cub.2023.11.012 | CC_BY |
| Movebank | 3131029654 | Milvus milvus | Jan Skrabal | Jan Skrabal | Škrábal J, Literák I, Raab R. 2023. Study "Milvus_milvus_Soaring_over_Adriatic_sea". Movebank Data Repository. <a href="https://doi.org/10.5441/001/1.300" target="_blank">https://doi.org/10.5441/001/1.300</a> <br><br> These data are described in<br> Škrábal J, Krejčí Š, Raab R, Sebastián-González E, Literák I. 2023. Soaring over open waters: horizontal winds provide lift to soaring migrants in weak thermal conditions. Movement Ecol. 11:76. https://doi.org/10.1186/s40462-023-00438-6 | CC_BY_NC |
| Movebank | 3133580132 | Lepus europaeus | Chloe tavernier | Chloe tavernier | | CC_BY_NC |

| | | | | | | |
|---|---|---|---|---|---|---|
| Movebank | 3179890710 | Vulpes vulpes | Tom Porteus | Tom Porteus | Porteus TA, Short MJ, Hoodless AN, Reynolds JC. 2024. Data from: Study "Red Fox (Vulpes vulpes) in UK wet grasslands". Movebank Data Repository. <a href="https://doi.org/10.5441/001/1.304" target="_blank">https://doi.org/10.5441/001/1.304</a> <br><br> These data are described in<br> Porteus TA, Short MJ, Hoodless AN, Reynolds JC. 2024. Movement ecology and minimum density estimates of red foxes in wet grassland habitats used by breeding wading birds. Eur J Wildlife Res. 70:8. https://doi.org/10.1007/s10344-023-01759-y | CC_0 |
| Movebank | 3186161092 | Anatidae ,Dendrocygna bicolor | Manuel Olivier Grosselet | Manuel Olivier Grosselet | | CC_0 |
| Movebank | 3218698288 | Rynchops niger | Manuel Olivier Grosselet | Manuel Olivier Grosselet | | CC_0 |
| Movebank | 3367931169 | Nucifraga caryocatactes | Eike Lena Neuschulz | Eike Lena Neuschulz | Graf V, Sorensen MC, Mueller T, Neuschulz EL. 2024. Data from: Study "Movement patterns of seed dispersing spotted nutcrackers (Nucifraga caryocatactes)". Movebank Data Repository. <a href="https://doi.org/10.5441/001/1.324" target="_blank">https://doi.org/10.5441/001/1.324</a> <br><br> These data are described in<br> Sorensen MC, Mueller T, Donoso I, Graf V, Merges D, Vanoni M, Fiedler W, Neuschulz EL. Scatter-hoarding birds disperse seeds to sites unfavorable for plant regeneration. Mov Ecol. 10:38. https://doi.org/10.1186/s40462-022-00338-1 <br><br> Graf V, Mueller T, Grüebler MU, Kormann UG, Albrecht J, Hertel AG, Sorensen MC, Tschumi M, Neuschulz EL. In press, 2024. Individual behavior shapes patterns of bird-mediated seed dispersal. Funct Ecol. | CC_BY |
| Movebank | 3413045568 | Streptopelia turtur | JIGUET Frédéric | Andrea Kölzsch | HABITRACK: Habitat tracking for the conservation of huntable bird species | CC_BY |
| Movebank | 3578422072 | Tyto furcata | Hermann Wagner | Hermann Wagner | | CC_0 |
| Movebank | 3791354435 | Panthera leo | Kevin MacFarlane | Kevin MacFarlane | MacFarlane, Kevin, 2014, The Ecology and Management of Kalahari lions in a Conflict Area in Central Botswana, PhD, Australian National University | CC_0 |

| | | | | | | |
|---|---|---|---|---|---|---|
| Movebank | 3809257699 | Panthera leo | Kevin MacFarlane | Robert Heinsohn | MacFarlane K. 2014. The ecology and management of Kalahari lions in a conflict area in central Botswana [dissertation]. [Canberra (Australia)]: Australian National University. https://doi.org/10.25911/5d78d54c1a50c | CC_BY |
| Movebank | 3902656356 | Nycticorax nycticorax | Mike Ward | Sarah Slayton | | CC_0 |
| Movebank | 3945977313 | Aphriza virgata | Shawn Crimmins | Sam Simon | | CC_0 |
| Movebank | 4447816442 | Rissa tridactyla | Jacob Davies | Jacob Davies | | CC_0 |
| Movebank | 4645303014 | Vultur gryphus | Hannah Williams | Hannah Williams | | CC_0 |
| Movebank | 4725153649 | Choloepus hoffmanni | Amelia Symeou | Amelia Symeou | | CC_0 |
| Movebank | 4732684199 | Bradypus variegatus | Rebecca Cliffe | Amelia Symeou | | CC_0 |
| Movebank | 4739876185 | Rallus ,Rallus elegans | Richard Temple Jr. | Richard Temple Jr. | | CC_0 |
| Movebank | 4901146318 | Connochaetes taurinus | Grant Hopcraft | Tom Morrison | Hopcraft, J.G.C., Morrison, T.A., and Stabach, J.A. (2024). Data from: Movement of resident wildebeest across the Masai Mara Ecosystem, Kenya. Movebank Data Repository. | CC_BY_NC |
| Movebank | 5175345606 | Lynx rufus,Urocyon cinereoargenteus | Laurel Serieys | Laurel Serieys | Serieys LEK, Matsushima SS, Wilmers CC. 2024. Data from: Study "Bobcat habitat connectivity study in central California". Movebank Data Repository. <a href="https://doi.org/https://doi.org/10.5441/001/1.323" target="_blank">https://doi.org/https://doi.org/10.5441/001/1.323</a> <br><br> These data are described in<br> Serieys LEK, Rogan MS, Matsushima SS, Wilmers CC. 2021. Road-crossings, vegetative cover, land use and poisons interact to influence corridor effectiveness. Biol Conserv. 253:108930. https://doi.org/10.1016/j.biocon.2020.108930 | CC_0 |
| Tucker at al 2023 | | Alces alces | Bram Van Moorter | | | |
| Tucker at al 2023 | | Alces alces | Matthew Kauffman | | | |

| | | | |
|---|---|---|---|
| Tucker at al 2023 | | Antilocapra americana | Chris Geremia |
| Tucker at al 2023 | | Antilocapra americana | Julie Young |
| Tucker at al 2023 | | Antilocapra americana | Matthew Kauffman |
| Tucker at al 2023 | | Canis aureus | Hubert Potocnik |
| Tucker at al 2023 | | Canis latrans | David Drake |
| Tucker at al 2023 | | Canis lupus | Daniel Stahler |
| Tucker at al 2023 | | Canis lupus | Jerrold Belant |
| Tucker at al 2023 | | Canis lupus | Mark Hebblewhite |
| Tucker at al 2023 | | Capreolus capreolus | Francesca Cagnacci |
| Tucker at al 2023 | | Cervus canadensis | Daniel Stahler |
| Tucker at al 2023 | | Cervus canadensis | Julie Young |
| Tucker at al 2023 | | Cervus canadensis | Kim Poole |
| Tucker at al 2023 | | Cervus elaphus | Erling Meisingset |
| Tucker at al 2023 | | Cervus elaphus | Johannes Signer |
| Tucker at al 2023 | | Cervus elaphus | Marco Heurich |
| Tucker at al 2023 | | Cervus elaphus | Peter Sunde |
| Tucker at al 2023 | | Chrysocyon brachyurus | Rogerio Cunha de Paula |
| Tucker at al 2023 | | Connochaetes taurinus | Grant Hopcraft |
| Tucker at al 2023 | | Equus hemionus | Petra Kaczensky / Buuveibaatar Bayarbaatar |

| | | | |
|---|---|---|---|
| Tucker at al 2023 | | Loxodonta africana | Morgan Hauptfleisch |
| Tucker at al 2023 | | Loxodonta africana | Robert Pringle |
| Tucker at al 2023 | | Odocoileus hemionus | Chris Geremia |
| Tucker at al 2023 | | Odocoileus hemionus | Julie Young |
| Tucker at al 2023 | | Odocoileus hemionus | Matthew Kauffman |
| Tucker at al 2023 | | Odocoileus virginianus | Guillaume Bastille-Rousseau |
| Tucker at al 2023 | | Odocoileus virginianus | Jerrold Belant |
| Tucker at al 2023 | | Odocoileus virginianus | Vickie DeNicola |
| Tucker at al 2023 | | Ovis canadensis | Chris Geremia |
| Tucker at al 2023 | | Ovis canadensis | Kevin Monteith |
| Tucker at al 2023 | | Ovis canadensis californiana | Julie Young |
| Tucker at al 2023 | | Ovis canadensis canadensis | Julie Young |
| Tucker at al 2023 | | Ovis canadensis nelsoni | Julie Young |
| Tucker at al 2023 | | Panthera leo | Jacob Goheen |
| Tucker at al 2023 | | Procapra gutturosa | Nandintsetseg Dejid |
| Tucker at al 2023 | | Puma concolor | Christopher Wilmers |
| Tucker at al 2023 | | Puma concolor | Julie Young |
| Tucker at al 2023 | | Rangifer tarandus | Allicia Kelly |
| Tucker at al 2023 | | Rangifer tarandus | Bram Van Moorter |
| Tucker at al 2023 | | Tragelaphus angasii | Robert Pringle |
| Tucker at al 2023 | | Tragelaphus strepsiceros | Robert Pringle |

| | | | |
|---|---|---|---|
| Tucker at al 2023 | | Tragelaphus sylvaticus | Robert Pringle |
| Tucker at al 2023 | | Ursus arctos | Jerrold Belant |
| Tucker at al 2023 | | Ursus arctos | Jonas Kindberg |
| Tucker at al 2023 | | Ursus arctos | Nuria Selva |

| Contact email |
| --- |
| |
| |
| |

| |
|---|
| |
| |
| |

| |
|---|
| |
| |
| |

| |
|---|
|<br><br> |
| |
| |

| |
|---|
| |
| |
| |
| |

| |
|---|
| |
| |

| |
|---|
| |
| |
| |
| |

| |
|---|
| |
| |
| |

| |
|---|
| |
| |
| |

| |
|---|
| |
| |

| |
|---|
| |
| |

| |
|---|
| |
| |
| |
| |
| |
| |
| |

| |
|---|
| |
| |
| |

| |
|---|
| |

| |
|---|
| |
| |

| |
|---|
| |
| |

| |
|---|
|<br><br><br><br><br><br><br><br><br><br><br><br><br><br><br><br><br><br><br><br><br><br><br><br><br><br><br><br><br><br><br><br><br><br> |

| |
|---|
| |
| |
| |
| |
| |

| |
|---|
| |
| |
| |

| |
|---|
| |

| |
|---|
| |
| |

| |
|---|
| |
| |
| |
| |
| |
| |

| |
|---|
| |

| |
|---|
| |
| |

| |
|---|
| |
| |
| |
| |

| |
|---|
| |
| |
| |

| |
|---|
| |
| |

| |
|---|
| |
| |
| |

| |
|---|
| |
| |
| |
| |
| |
| |
| |
| |

| |
|---|
| |
| |
| |
| |
| |
| |

| |
|---|
| |
| |
| |

| |
|---|
| |
| |
| |

| |
|---|
| |
|<br> |
| |

| |
|---|
| |
| |
| |
| |

| |
|---|
| |
| |
| |
| |

| |
|---|
| |
| |

| |
|---|
| |
| |
| |

| |
|---|
| |
| |

| |
|---|
| |
| |
| |

| |
|---|
| |
| |
| |

| |
|---|
| |
| |
| |

| |
|---|
| |
| |
| |

| |
|---|
| |
| |
| |
| |

| |
|---|
| |
| |
| |
| |
| |

| |
|---|
| |
| |

| |
|---|
| |
| |
| |
| |
| |
| |
| |

| |
|---|
| |
| |
| |
| |
| |

| |
|---|
| |
| |
| |
| |

| |
|---|
| |
| |
| |
| |
| |
| |
| |
| |
| |

| |
|---|
| |
| |
| |
| |

| |
|---|
| |
| |
| |
| |
| |
| |
| |
| |

| |
|---|
| |
| |
| |
| |
| |

| |
|---|
| |
| |

| |
|---|
| |
| |
| |
| |
| |

| |
|---|
| |
| |
| |
| |
| |
| |
| |

| |
|---|
| |
| |
| |
| |
| |

| |
|---|
| |
| |

| |
|---|
| |
| |
| |
| |

| |
|---|
| |
|<br><br> |

| |
|---|
| |
| |
| |
| |

| |
|---|
| |
| |
| |
| |
| |
| |
| |

| |
|---|
| |
| |
| |
| |
| |
| |
| |
| |
| |
| |
| |
| |

| |
|---|
| |
| |
| |
| |
| |
| |
| |
| |
| |
| |
| |
| |
| |
| |
| |
| |
| |
| |
| |

| |
|---|
| |
| |
| |
| |
| |
| |
| |
| |
| |
| |
| |
| |
| |
| |
| |
| |
| |
| |
| |
| |
| |

| |
|---|
| |
| |
| |
| |
